# A chromosome-assigned Mongolian gerbil genome with sequenced centromeres provides evidence of a new chromosome

**DOI:** 10.1101/2022.09.21.508825

**Authors:** Thomas D. Brekke, Alexander S. T. Papadopulos, Eva Julià, Oscar Fornas, Beiyuan Fu, Fengtang Yang, Roberto de la Fuente, Jesus Page, Tobias Baril, Alexander Hayward, John F. Mulley

## Abstract

Chromosome-scale genome assemblies based on ultra-long read sequencing technologies are able to illuminate previously intractable aspects of genome biology such as fine-scale centromere structure and large-scale variation in genome features such as heterochromatin, GC content, recombination rate, and gene content. We present here a new chromosome-scale genome of the Mongolian gerbil (*Meriones unguiculatus*) which includes the complete sequence of all centromeres. Gerbil centromeres are composed of four different repeats of length 6pb, 37bp, 127bp, or 1747bp which occur in simple alternating arrays and span 1-6Mb. Gerbil genomes have both an extensive set of GC-rich genes and chromosomes strikingly enriched for constitutive heterochromatin. We sought to determine if there was a link between these two phenomena and found that the two heterochromatic chromosomes of the Mongolian gerbil have distinct underpinnings: Chromosome 5 has a large block of intra-arm heterochromatin as the result of a massive expansion of centromeric repeats, while chromosome 13 is comprised of extremely large (>150kb) repeated sequences. In addition to characterizing centromeres, our results demonstrate the importance of including karyotypic features such as chromosome number and the locations of centromeres in the interpretation of genome sequence data, and highlight novel patterns involved in the evolution of chromosomes.

## Introduction

Understanding genome organization and function and how they vary has been an important goal in the field of biology since at least the 1950s. The new and relatively inexpensive long-range sequencing technologies such as PacBio HiFi and Oxford Nanopore are facilitating the sequencing and chromosome-scale assembly of the genomes of many new species (Jayakumar and Sakakibara 2017). Such high-quality genomes are an important tool to address long-standing questions about variation in the structure and function of genomes across the tree of life. Such questions include: What is the nucleotide sequence and structure of centromeres in non-model species? What is the recombination landscape and how does it influence nucleotide content variation in genes and along chromosomes? In addition and often overlooked: what new insights can be gleaned when we reinterpret cytological data, such as the banding patterns of chromosomes in a karyotype, in light of chromosome-scale assemblies?

Centromeres are crucially important during mitosis and meiosis. Functionally, they are the binding site of centromere-specific histones and other proteins which facilitate their binding to the spindle apparatus (McKinley and Cheeseman 2016). They are visible in karyotypes as constrictions in the chromosome which stain very darkly under different chemical treatments (Willard 1990). They are characterized by arrays of various repeated sequences of DNA of various lengths, where the sequence of the repeat is species-specific (Talbert and Henikoff 2020). Due to their size and repetitive nature, they have proven intractable to assembly by all but the most recent of long-range sequencing technologies, indeed it is only within the last year that human centromeres have been completely sequenced and annotated (Altemose et al. 2022). An immense amount of work has gone into studying centromeres at the functional level using visualization techniques and the like, but very little is known about the specific sequence of centromeres in most species. Sequencing and characterizing centromeres in various non-model species is and will be an important addition to understanding the variation and function of centromeres across the tree of life.

The nucleotide composition of genomes is not homogenous; it varies along chromosome arms and between chromosomes, individuals, populations, and species (Eyre-Walker and Hurst 2001). Variation in the distribution of guanine (G) and cytosine (C) bases is heavily determined by the recombination-associated process of GC-biased gene conversion (gBGC), which favours fixation of guanine and cytosine over adenine (A) and thymine (T)(Lamb 1984; Arbeithuber et al. 2015). Over evolutionary time this process results in a GC bias around recombination hotspots (Galtier et al. 2001). Gerbils and their relatives have multiple extensive regions of extremely high GC bias within their genomes, higher than that seen in any other mammal (Hargreaves et al. 2017; Dai et al. 2020; Pracana et al. 2020). Historically, this has complicated attempts to obtain high-quality contiguous gerbil genome assemblies (Leibowitz et al. 2001; Gustavsen et al. 2008). Intriguingly, there appear to be two distinct patterns of GC skew in gerbils: (i) a region associated with the ParaHox cluster and the surrounding genes, where virtually all genes in this region have very high mutation rates and an extreme GC bias, and (ii) a further set of 17 large clusters of GC-rich genes also with high mutation rates (Pracana et al. 2020). These intriguing characteristics of gerbil genomes make them an ideal system in which to examine the association between GC biased gene conversion and the organization of eukaryotic genomes.

Chromatin state is an important mechanism for the regulation of gene activity. Facultative heterochromatin is cell-type-specific and may be converted to open, active euchromatin during gene regulatory processes. In contrast, constitutive heterochromatin is marked by tri-methylation of histone H3 at the lysine 9 residue (H3K9me3) (Saksouk et al. 2015) and comprises densely compacted, gene-poor inactive regions of the genome which are condensed in all cell types at all developmental stages, such as centromeres and telomeres (Saksouk et al. 2015; Penagos-Puig and Furlan-Magaril 2020). Many gerbil species (Family Gerbillidae) have chromosomes with high levels of constitutive heterochromatin, though the specific chromosome and extent of heterochromatin vary by species. Mongolian gerbils possess distinctive karyotypic features: nearly a third of Chromosome 5 and all of Chromosome 13 appear to be composed of constitutive heterochromatin by multiple different assays: Chromosome 13 stains entirely dark in C-banding stains (Gamperl and Vistorin 1980) and is completely coated by heterochromatin histone marks in immunofluorescence assays (de la Fuente et al. 2014).(Gamperl and Vistorin 1980) The genomes of the North African Gerbil (*Gerbillus campestris*), the hairy-footed gerbil (*Gerbilliscus paeba*), and the fat sandrat (*Psammomys obesus*) all contain a single heterochromatic chromosome (Solari and Ashley 1977; Gamperl and Vistorin 1980; Knight et al. 2013).

The heterochromatic chromosomes in gerbils are present in all individuals examined to date and do not meet the criteria for classification as B chromosomes, i.e.: they are not non-essential, and do not vary in copy number among individuals and tissues without an adverse impact on fitness (Ahmad and Martins 2019). These chromosomes therefore provide a unique system to examine the impact of their heterochromatic state on genic evolution and particularly whether it is linked to the extensive number of GC-rich genes in gerbil genomes. Heterochromatin is typically gene-poor (Dimitri et al. 2005) and transcriptionally repressed (Grewal and Moazed 2003; Dillon 2004). This makes it unlikely that entire heterochromatic chromosomes would be maintained and transmitted across generations for millions of years if they did not encode any genes or are entirely selfish independent genetic elements. High GC% in certain gerbil genes could be an adaptation to a transcriptionally-repressive environment. Genes with high GC% in their coding regions and adjacent regions of DNA, and especially those with high GC% in the 3^rd^ codon position (GC_3_) can show elevated expression (Lercher et al. 2002; Vinogradov 2005). Conversely, since gBGC is a recombination-dependent process, and since all chromosomes must undergo at least one reciprocal recombination event (crossover) with their homologue during meiosis (Lydall et al. 1996), an alternative hypothesis is that the extreme GC% present in some gerbil genes is a consequence their becoming entrapped in or near a recombination hotspot. If the bulk of the extensive heterochromatin observed on these gerbil chromosomes is non-permissive to recombination, then genes in those regions where recombination can occur will become increasingly GC-rich because of continual exposure to gBGC. We may therefore reasonably expect a link between GC-rich genes and these unusual gerbil chromosomes.

A key question is how did fully heterochomatic chromosomes in gerbils arise? They may once have been “normal” chromosomes that have degenerated into gene-poor, non-functional, or silenced chromosomes by accumulation of repetitive DNA. Alternatively, they may have formed from heterochromatic pieces that broke off from other chromosomes, in the same way that the neochromosomes of tumors (Garsed et al. 2009, 2014) and some B chromosomes (Camacho et al. 2000; Dhar et al. 2002) develop from fragments of other chromosomes.

Alternatively they could be the duplicate of another chromosome, which condensed into heterochromatin a mechanism of dosage compensation in the same way that additional copies of X chromosomes are inactivated in female mammals (Lyon 1962). Finally, they may potentially have grown from a smaller chromosomal “seed”, which broke off from another chromosome, and subsequently grew by repeated segmental duplication.

Until very recently, questions such as those posed above could not be addressed in a non-model system for several key reasons. A particularly important issue was the difficulties that short read-based genome sequencing approaches face regarding the assembly of GC%-rich regions (Hron et al. 2015; Bornelöv et al. 2017; Botero-Castro et al. 2017; Tilak et al. 2018; Yin et al. 2019). Meanwhile, the current trend towards the generation of chromosome-scale assemblies has perhaps lost sight of the importance of an understanding of the karyotype of the species being studied, and of physically linking genome sequence to identified chromosomes.

Using a new chromosome-scale genome assembly for the Mongolian gerbil and methods enabling us to assign the genomic scaffolds to physical chromosomes, we first characterize gerbil centromeres and then test (i) whether GC-rich gene clusters correlate with recombination hotspots and (ii) if those genes are associated with a single heterochromatic chromosome. Our approach allows us to examine the origin and propose a new hypothesis for the evolution of some unusual and possibly unique, heterochromatic gerbil chromosomes.

## Results and Discussion

### Gerbil genome: approach and summary statistics

We sequenced and assembled the genome of the Mongolian gerbil, *Meriones unguiculatus* into 245 contigs using PacBio HiFi reads and scaffolded OmniC chromatin conformation capture data (Figure S2), Oxford Nanopore long and ultra-long read sequence data, a genetic map (Table S1) (Brekke et al. 2019), and BioNano optical mapping. We assigned scaffolds to chromosomes by flow-sorting chromosomes into pools. Each pool was sequenced with Illumina short reads, and these reads used to determine which scaffolds associated with each pool. The sorted pools were also made into FISH paint probes to identify which physical chromosome from the karyotype associated with each pool. This approach linked the physical chromosomes with sequenced scaffolds (full methods are in Supplemental Material 1, Figures S1, S3-S7, and Tables S2 and S3). The final genome assembly contains 194 scaffolds spanning 21 autosomes, the X and Y sex chromosomes, and the mitochondrial genome (Table S4). For 20 of the 23 chromosomes, a single large scaffold contains over 94% (often over 99%) of all the sequence assigned to that chromosome (Figure 1A). Only chromosome 13, with 121 scaffolds, and the X and Y chromosomes, each with 6 scaffolds, are appreciably fragmented and there are only 30 unassigned scaffolds making up 0.066% of the sequenced bases (Figure 1B). The assembly was annotated using RNAseq data and is 92% complete based on a BUSCO analysis (Complete:92.3% [Single-copy:91.7%, Duplicated:0.6%], Fragmented:1.7%, Missing:6.0%, n:13798) (Manni et al. 2021). We used the program NeSSie (Berselli et al. 2018) to calculate two measures of sequence complexity (entropy and linguistic complexity) in sliding windows across every chromosome. The complexity metrics revealed two chromosomes with unusual features: Chromosome 5 which has an extensive region where both entropy and linguistic complexity are very low, and Chromosome 13 which shows a marked homogeneity in its entropy across the length of the chromosome (Figure 2 and Figure S8). A chromosome-by-chromosome summary of all data is found in Figure 3 and a high-resolution version is in Supplement 2.

**Figure 1:**
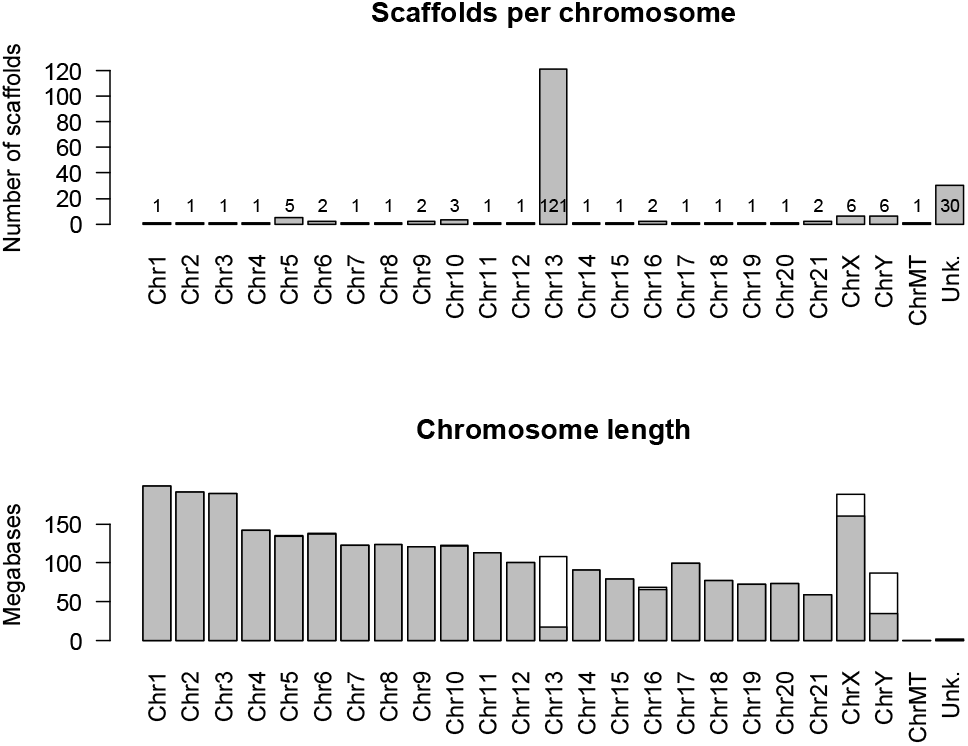
Summary statistics for the Mongolian gerbil (*Meriones unguiculatus*) genome assembly. Top: The number of scaffolds assigned to each chromosome, the mitochondrial genome, and the ‘unknown’ category. Most chromosomes are assembled into 1 or 2 scaffolds, while chromosome 13 is in 121 pieces. Bottom: The number of bases assigned to each chromosome with the single longest scaffold shaded in grey. The total amount of DNA sequence assigned to chromosome 13 is about what would be expected, showing that we are not missing data, and that the large number of scaffolds is not an artefact.

**Figure 2:**
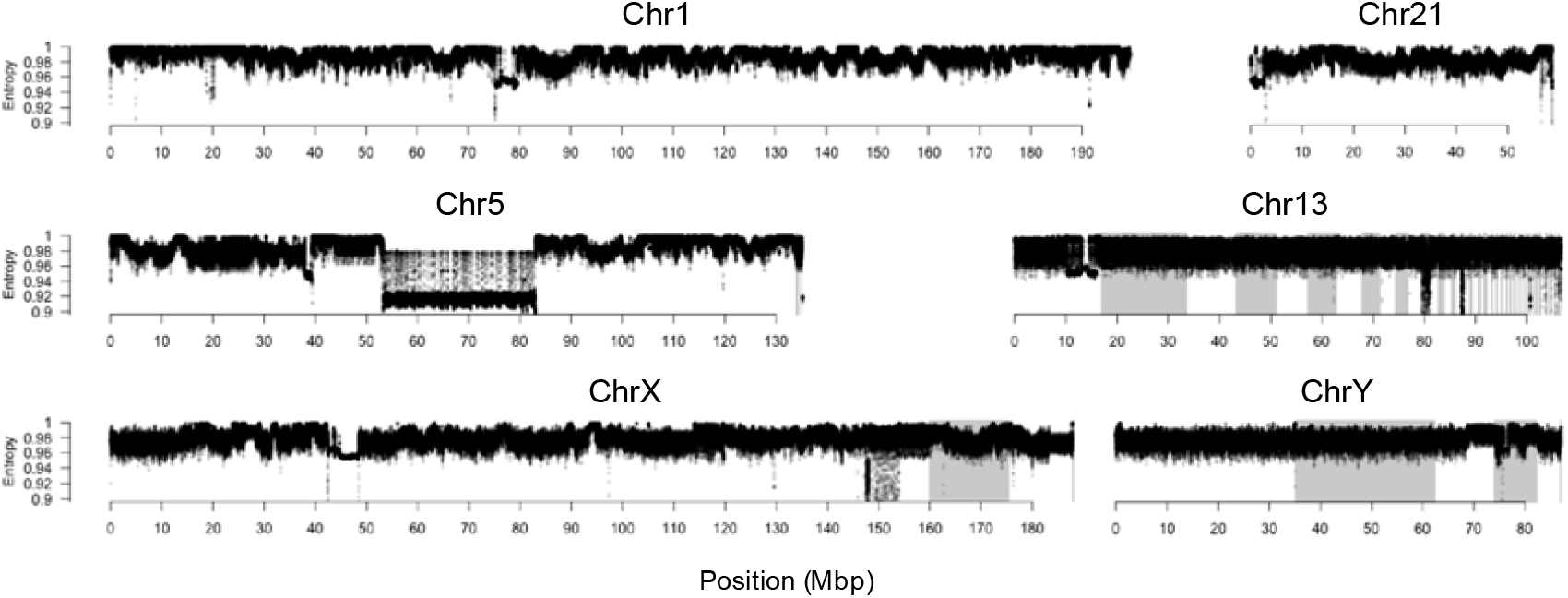
Entropy plots for a selection of chromosomes including the morphologically standard autosomes 1 and 21, the unusual autosomes 5 and 13, and both sex chromosomes. The unordered scaffolds within a chromosome are shaded alternately white and grey. Centromeres are apparent at ~75-80Mbp in Chr1, ~0-5Mbp in Chr21, ~35-40Mbp in Chr5, ~14-17Mbp in Chr13, ~45-50Mbp in ChrX, and ~75Mbp in ChrY. Note the spatial heterogeneity in chromosomes 1 and 21 that is absent in chromosome 13 and the Y. Indeed, chromosome 13 is the most homogenous chromosome in the gerbil. Entropy plots for every chromosome, as well as GC content, gene density, and linguistic complexity can be found in Figure S2.

**Figure 3:**
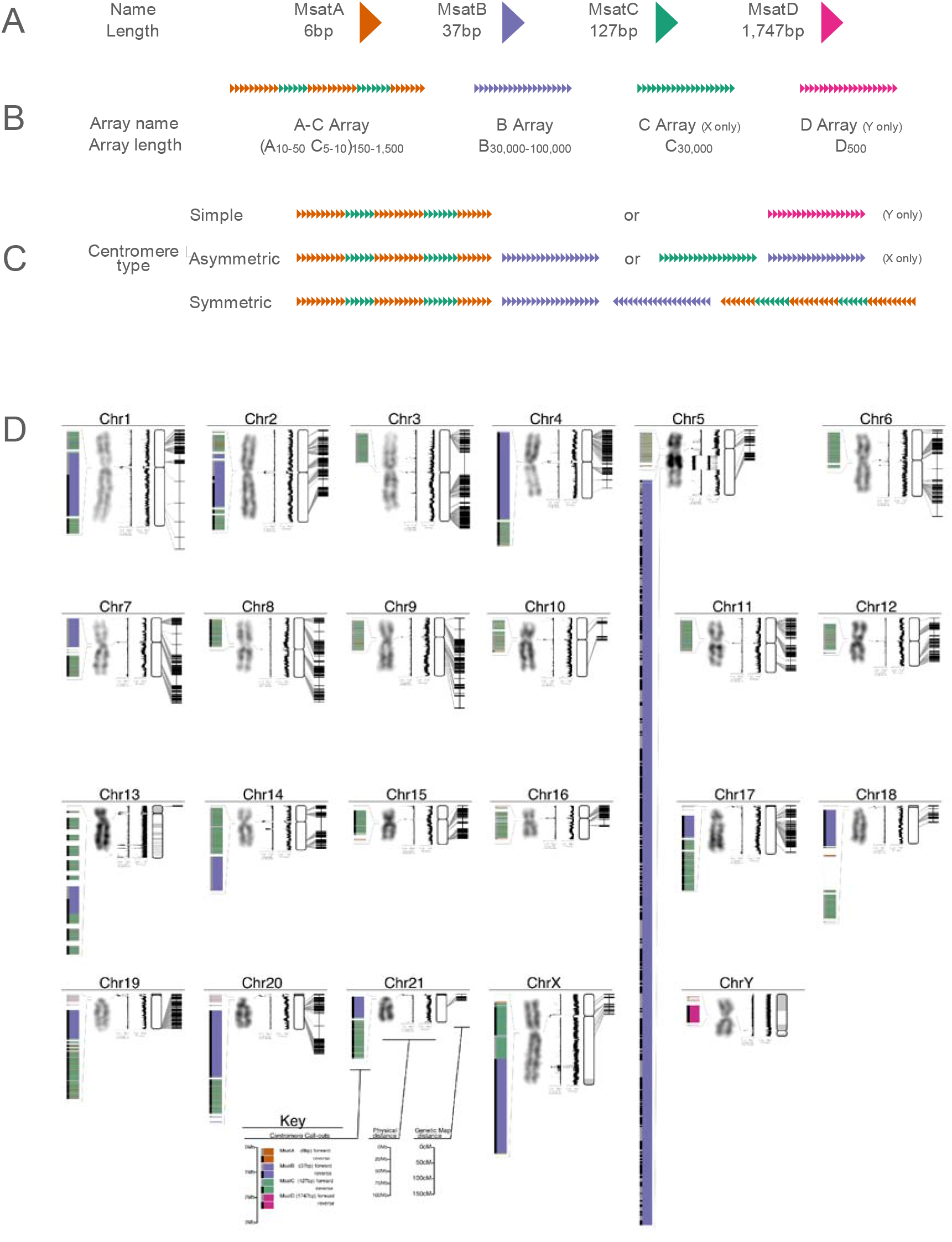
The Mongolian gerbil (*Meriones unguiculatus*) genome. Gerbil centromere types. (A) There are four different repeat types in gerbil centromeres: MsatA (6bp), MsatB (37bp), MsatC (127bp), and MsatD (1,747 bp). (B) These repeats appear in one of four repeat arrays. The A-C array consists of 10-50 copies of MsatA alternating with 5-10 copies of MsatC, all of which is repeated 150-1,500 times. The B-, C-, and D-arrays contain only multiple copies of their respective repeat. Repeat units within an array most often occur in the same orientation. In some chromosomes however both orientations occur within a single array, in which case hundreds of repeat units in the forward orientation are followed by hundreds of units in the reverse orientation (e.g. the B array of Chromosome 2 in Figure 2). (C) Centromeres consist of between one and three repeat arrays and are classed as either ‘simple’, ‘asymmetric’, or ‘symmetric’. Simple centromeres have a single array type, either an A-C array as in the autosomes, or a D array as on the Y Chromosome. Asymmetric centromeres have two arrays: either an A-C array and a B array (for the autosomes) or a C array and a B array (for the X chromosome). Symmetric centromeres consist of three arrays, a B array sandwiched between two A-C arrays which typically appear in opposite orientation to each other. (D) Genome schematic, for each chromosome we show, from left to right: (1) centromere organization, with repeats of different lengths in different colors and the orientation of the repeat array denoted by a grey or black bar on the left. Chromosome 5 has a large expansion of centromeric repeats in the long arm. All call-outs are drawn to the same scale. (2) The DAPI-banding karyotype image, showing the intra-arm heterochromatin on chromosome 5, and the entirely dark staining on chromosome 13. (3) Linguistic complexity and (4) entropy, both measured in overlapping sliding 10kb windows with a step size of 1kb. For both metrics, a low value indicates highly repetitive or predictable sequence as are characteristic of centromeres while high values indicate more complex sequence as may be found in gene-rich regions. (5) A depiction of the physical map with unplaced scaffolds organized by length and shaded alternately white and grey, and (6) a depiction of the genetic map with links between the genetic markers and their physical location. Thin grey lines link the location of similar features on adjacent plots (i.e. centromere callout to karyotype; centromere location in the karyotype to centromere in the linguistic complexity plot; genetic markers to their physical location). A high-resolution copy of panel D can be found in the supplement (Figure SC)

Two *M. unguiculatus* genome sequences have been previously published, based on short-read sequence data (Cheng et al. 2019; Zorio et al. 2019), both contain hundreds of thousands of contigs and equally large numbers of scaffolds (Table S4). One of these has recently been improved with Hi-C data (www.DNAZoo.org) into 22 chromosome-length scaffolds, and ~300,000 additional scaffolds (Cheng et al. 2019). Full-genome alignments between our genome assembly and this Hi-C assembly (Figure S9) showed that most scaffolds are colinear between the assemblies but that the “improved” Cheng et al. (Cheng et al. 2019) assembly lacks chromosome 13 entirely, hence only 22 chromosome-scale scaffolds for this n=23 species. Our highly contiguous and physically associated assembly provides the foundation for all subsequent analyses.

### Characterization of gerbil centromeres

Relatively little is known about centromere organization in non-model species, as centromeres are comprised of extensive runs of repeated sequences, which short-read technologies (and even Sanger sequencing) have struggled to cross. It is only this year that full coverage of human centromeres was obtained, from a mixture of long-read sequencing approaches applied to the genome of a hydatidiform mole cell line by the Telomere-to-Telomere (T2T) consortium (Altemose et al. 2022). Our high-quality PacBio HiFi-derived sequence data resulted in a single large scaffold per chromosome (for all but a few chromsomes) which spanned from telomere to telomere (Figure 1). Such completeness suggested that we sequenced through the centromeres of all *M. unguiculatus* chromosomes and so we set about bioinformatically identifying centromeres. Centromeres are known to be highly repetitive, occur once on each chromosome, are visually apparent as a constriction in the karyotype, and are typically on the order of a few megabases long (Talbert and Henikoff 2020). We used the entropy and linguistic complexity metrics (measures of sequence repetitiveness) to reveal a region of each chromosome that matched the above predictions: every chromosome has a single highly repetitive region ranging from ~1-10Mbp long which line up with the constriction in the karyotype (Figure 2 and Figure 3). As we did no molecular assay for centromere function, we submit these as “putative centromeres”, though for brevity, we hereafter refer to them simply as “centromeres”.

To further characterize the gerbil centromeres, we used the program NTRprism (Altemose et al. 2022) which identified four different simple repeat sequences (Figure S10). We have named these “MsatA” (for *<u>M</u>eriones* <u>sat</u>ellite <u>A</u>), “MsatB”, “MsatC”, and “MsatD” (Figure 3A): MsatA is 6bp long and has the sequence TTAGGG which is the same simple sequence repeat found in telomeres, MsatB is 37bp long, MsatC is 127bp long, and MsatD is 1,747bp long and is only found on the Y chromosome. A representative sequence of each Msat can be found in the legend of Figure S10. At the time of writing, MsatB and MsatC return no BLAST hits from NCBI’s ‘nt’ library (update 2023/01/12) and MsatD returns a single 32bp run of identity (out of 1748bp) with *Acomys russatus* suggesting that these Msats are new sequences not previously identified.

Copies of Msats are arranged into one of four variant arrays which define an intermediate-order structure in the centromeres (Figure 3B). ‘B arrays’ are formed from copies of MsatB and range in size from 1Mbp to 3Mbp long (~30,000 to ~100,000 copies). Similarly, the Y chromosome centromere is a ‘D array’ comprised of ~500 copies of MsatD spanning slightly less than a megabase. MsatA and MsatC repeats are rarely found alone, tending instead to intersperse with each other to form ‘A-C arrays’. Typically 10-50 copies of MsatA will alternate with 5-10 copies of an MsatC unit, and this alternating pattern will extend for between 100Kb and 1Mb depending on the chromosome. The only place that MsatC are found without interspersed copies of MsatA is on the X chromosome in what we term a ‘C array’. While not interspersed with MsatC, there are a number of MsatA repeats that do appear at both ends of the X centromere and are detectable by FISH (de la Fuente et al. 2014). The orientation of the Msat repeats is typically consistent across an array, however some arrays are composed of blocks of Msat repeats in alternating orientations with many copies of repeat in the forward orientation followed by many copies in the reverse orientation.

The highest-level of centromere organization is characterized by groups of between one and three arrays which fall into one of a few patterns which we term ‘simple’, ‘asymmetric’, or ‘symmetric’ (Figure 3C). Simple centromeres are comprised of a single A-C array and are present in ten of the smaller metacentric chromosomes (see chromosomes 3, 5, 6, 8-12, 15, and 16, Figure 3D). The metacentric Y chromosome also has a simple centromere, though with a D array instead of the A-C array. Asymmetric centromeres are comprised of two arrays, one of which is always a B array and the other is typically an A-C array. Eight autosomes fall into this category including all four of the small telocentric chromosomes (chromosomes 18-21, Figure 3D), three of the metacentric chromosomes (chromosomes 4, 7, and 14, Figure 3D), and one acrocentric chromosome (chromosome 17, Figure 3D). The metacentric X chromosome also has an asymmetric centromere but is the only location in the genome where a pure C array exists. Finally, symmetric centromeres are comprised of three arrays: a C array sandwiched between two A-C arrays and are found in the metacentric chromosomes 1, and 2, and the acrocentric chromosome 13. Many centromeres also contain 10Kbp–50Kbp blocks of non-repetitive, complex DNA both between and within the various arrays (see Figure 3D).

### The location of GC-rich genes

A set of over 380 genes with extreme GC content clustered in the genomes of sandrats and gerbils has previously been identified (Pracana et al 2020). It has been hypothesized that biased gene conversion has driven their GC content to extraordinary levels since they are near recombination hotspots (Pracana et al. 2020), but the resources to test this were not available so mouse gene locations had been used as an evolutionarily-informed proxy for the location of those genes in gerbils. Here we use our newly-generated chromosome-scale assembly to explicitly test how these GC-rich genes are distributed across gerbil chromosomes. We used a permutation test to show that GC-rich genes are clustered together more than is expected by chance (Figure 4A, observed = 1.71Mbp, mean = 2.89Mbp, n=1,000,000 permutations, p < 0.000001). We used our genetic map (Brekke et al. 2019) to locate recombination hotspots which were defined as regions with 5x higher recombination rate than the genome average (as per (Katzer et al. 2011). Hotspots were found on 18 of 22 chromosomes (21 autosomes and the X chromosome, we omit the Y chromosome here as it does not recombine) with 2.4+/-2.2(sd) hotspots per chromosome (Figures S11, S12, S13). Chromosomes 2, 18, 21, and the X lack recombination hotspots. We tested proximity of GC-rich genes to hotspots in two ways, first by comparing the GC-rich genes with the entire gene set (Figure S14) and secondly by performing a permutation test (Figure 4B). In both cases, GC outlier genes were found to lie significantly closer to recombination hotspots than expected by chance (Figure 4B, observed = 21.68Mbp, mean = 27.29Mbp, n = 1,000,000, p < 0.00058). These results demonstrate a clear association of GC rich gene clusters with recombination hotspots as expected under gBGC.

**Figure 4:**
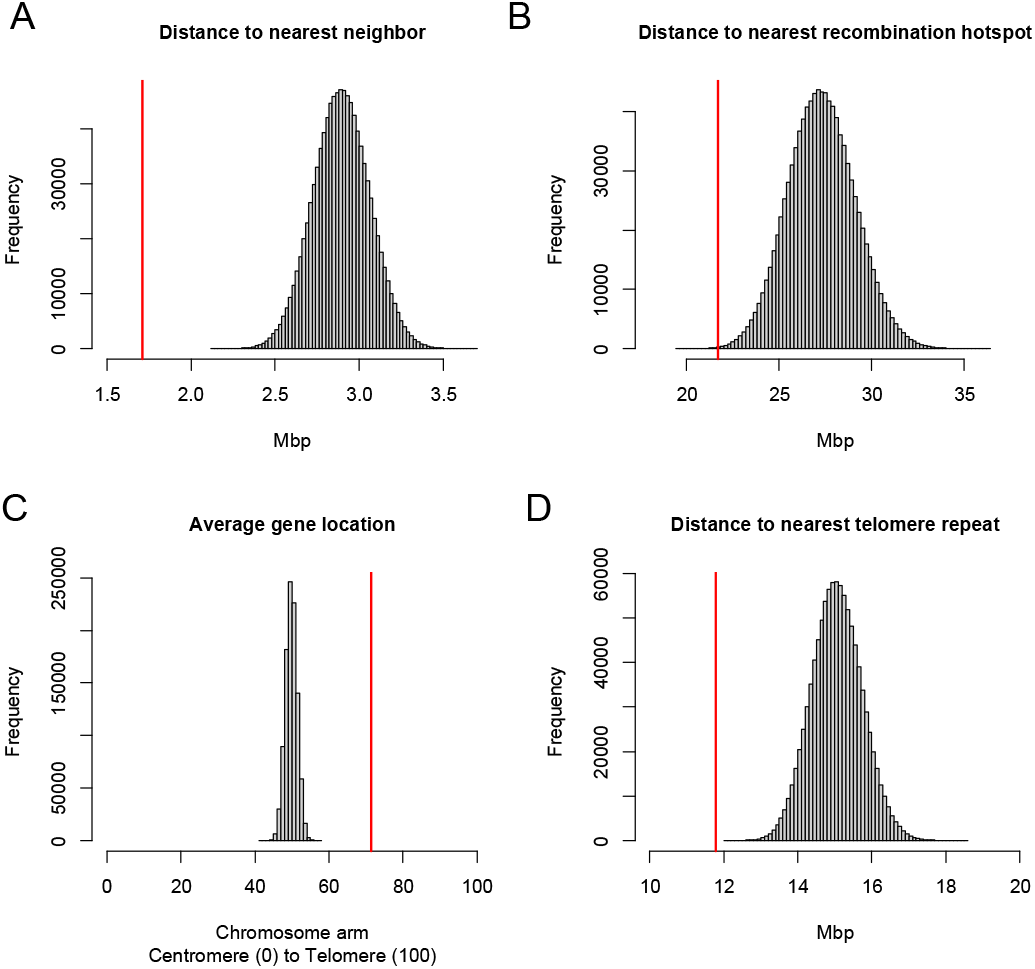
GC-rich genes are nonrandomly distributed in the *M. unguiculatus* genome. We compared the location of the 410 GC rich genes (Pracana et al. 2020)in relation to each other, the nearest recombination hotspot, their location along the chromosome arm, and their proximity to telomere repeats both interstitial and at the ends of chromosome arms. These comparisons were done once against the entire gene set (Figures S14, S15, and S17) and here using a permutation test with 1,000,000 draws of a random set of 410 genes where the red line indicates the observed value. (A) GC rich genes are clustered in the genome. The observed distance between each outlier gene and its nearest outlier gene neighbor is significantly shorter than those distances between a random group of genes (observed = 1.71Mbp, mean = 2.89Mbp, n=1,000,000 permutations, p < 0.000001). (B) GC-rich genes occur closer to recombination hotspots than expected by chance (observed = 21.68Mbp, mean = 27.29Mbp, n = 1,000,000, p < 0.00058). (C) GC rich genes are found closer to the telomere end of chromosome arms than expected by chance (observed = 71.46%, mean = 49.78%, n=1,000,000, p < 0.000001). (D) GC-rich genes are clustered nearer telomere repeats (interstitial or otherwise) than expected by chance (observed = 11.79Mbp, mean = 15.06Mbp, n=1,000,000, p < 0.000001).

While a genetic map shows the location of current recombination hotspots, hotspots move through evolutionary time due to large-scale chromosomal rearrangements and the mutational load caused by crossing over (Paigen and Petkov 2010; Tiemann-Boege et al. 2017). Consequently, we next tested whether GC outliers are associated with proxies of ancestral hotspots. Recombination rate is not uniform across a chromosome and is typically higher near the telomeres (Nachman 2002; Martinez-Perez and Colaiácovo 2009), thus we tested whether GC outliers are correlated with position along the chromosome arm. We found that whether considering the full distribution of gene locations (Figure S15) or 1,000,000 draws of the same number of random genes in a permutation test (Figure 4C), the GC outliers are found to lie much closer to the telomere than expected by chance (Figure 4C, observed = 71.46%, mean = 49.78%, n=1,000,000, p < 0.000001). Furthermore, gerbils have many interstitial telomere sites (de la Fuente et al. 2014) which are caused by chromosomal fusions embedding what was an ancestral telomere within a chromosome arm, typically near the centromere. Thus, interstitial telomere repeats are proxies for the ends of ancestral chromosomes and their associated ancient recombination hotspots. We identified interstitial telomere sites as arrays of the 6bp “MsatA” with at least 70 tandem copies (Figure S16). We therefore tested whether GC outlier genes are closer to these telomere repeats (which could be interstitial or otherwise) than expected by chance and found that they are (Figure 4D, observed = 11.79Mbp, mean = 15.06Mbp, n=1,000,000, p < 0.000001; Figure S17). In short, GC outlier genes are found in clusters across the genome and are nearer to recombination hotspots (current or ancient) and telomere/interstitial telomere sites than expected by chance, strongly supporting the hypothesis that GC-biased gene conversion is driving the extreme GC content of these genes. Figure S18 shows the distribution of centromeres, recombination hotspots, high GC genes, and telomere sites that were used in these analyses.

However, we did not find that all GC-rich genes are located on heterochromatic chromosomes and find instead that they are distributed on the order of 19.5±13.7 GC-rich genes per chromosome across the genome. The tendency for genes to become highly GC-rich in and around recombination hotspots in gerbils therefore seems unrelated to their unusual chromosomes and may instead be the result of greater recombination hotspot stability, where hotspots stay in one place for longer in gerbils compared to other species. Similarly stable hotspot location has previously been reported for birds (Singhal et al. 2015) though in birds the absence of PRDM9 correlates with greater hotspot stability. The gerbil genome encodes a full-length *Prdm9* gene on chromosome 20, and so this hotspot stability in gerbils must arise via some other mechanism.

We next sought to understand the genomic basis of the heterochromatic appearance of the chromosomes 5 and 13 in *M. unguiculatus*.

### Chromosome 5: the relevance of centromeric drive

Chromosome 5 is characterized by an enormous centromeric repeat expansion which is visible as a dark band on the q arm (Figure 3D). Our data shows that the repeat expansion is a 29Mb long B array, which comprises approximately 22% of the entire chromosome. This repeat expansion is distinct from the centromere which is a simple A-C array 1.5Mb long. In contrast to the B arrays in the centromeres of other chromosomes, the orientation of MsatB repeats on chromosome 5 switches far more frequently. With a few exceptions, B arrays in centromeres maintain their orientation across the entire array, or in the case of the symmetric centromeres, have a few large blocks in opposite orientations; the centromeric B arrays maintain orientation for 1-3Mb. Repeats in the Chromosome 5 expansion however, switch orientation over 200 times across the 29Mb, so the average block length is just 140Kb.

There is a similar large expansion of a centromeric repeat found in human chromosome 9 (Altemose et al. 2022). However, while it is similar in size to the expansion on gerbil chromosome 5, the human expansion is polymorphic in the population (Craig-Holmes and Shaw 1971). The dark band on the q arm of gerbil chromosome 5 is visible in all published karyotypes dating back to the 1960s which derive from many different individuals and laboratory colonies (Pakes 1969; Weiss et al. 1970; Gamperl and Vistorin 1980) suggesting that in contrast, the gerbil expansion is fixed at this massive size.

The repeat expansion is absent in karyotypes of many closely related Gerbillinae species, including representatives from the genera *Desmodilus, Gerbillurus, Gerbillus, Tatera*, and *Taterillus*, and is even absent in other species of *Meriones*. (Gamperl and Vistorin 1980; Benazzou et al. 1982, 1984; Qumsiyeh 1986b,a; Dobigny et al. 2002; Aniskin et al. 2006; Volobouev et al. 2007; Gauthier et al. 2010). The expansion is also absent in the sequenced genome assemblies of the closely related fat sandrat (*Psammomys obesus*) and fat-tailed gerbil (*Pachyuromys duprasi*). Alignment with the *Psammomys* genome assembly shows that the location of the repeat expansion on *M. unguiculatus* chromosome 5 is homologous to the *Psammomys* chromosome 10 centromere (Figure S19), suggesting that the region in *M. unguiculatus* is an ancestral centromere that has expanded. The centromere-drive hypothesis (Malik 2009) may explain the distribution of array types in the autosomal centromeres under the following model: the ancestral gerbil centromeres were predominately B arrays and at some point after the *Meriones – Psammomys* split, centromeric drive triggered a massive repeat expansion of the B array on what would become *Meriones* chromosome 5. This runaway expansion was the catalyst for genome-wide centromere turnover, where A-C arrays replaced B arrays as the new functional centromeres and many B arrays were evolutionarily lost, with those that remained being non-functional relics. Indeed, the centromere expansion on chromosome 5 does not bind CENT proteins, although it preserves other heterochromatic marks, (such as H3K9me3) and excludes recombination events, as assessed in male meiosis by the localization of the recombination marker MLH1 (Figure 5). While the heterochromatic state of a large portion of chromosome 5 can therefore be explained by the massive expansion of a centromeric repeat, this is not the case for chromosome 13.

**Figure 5.**
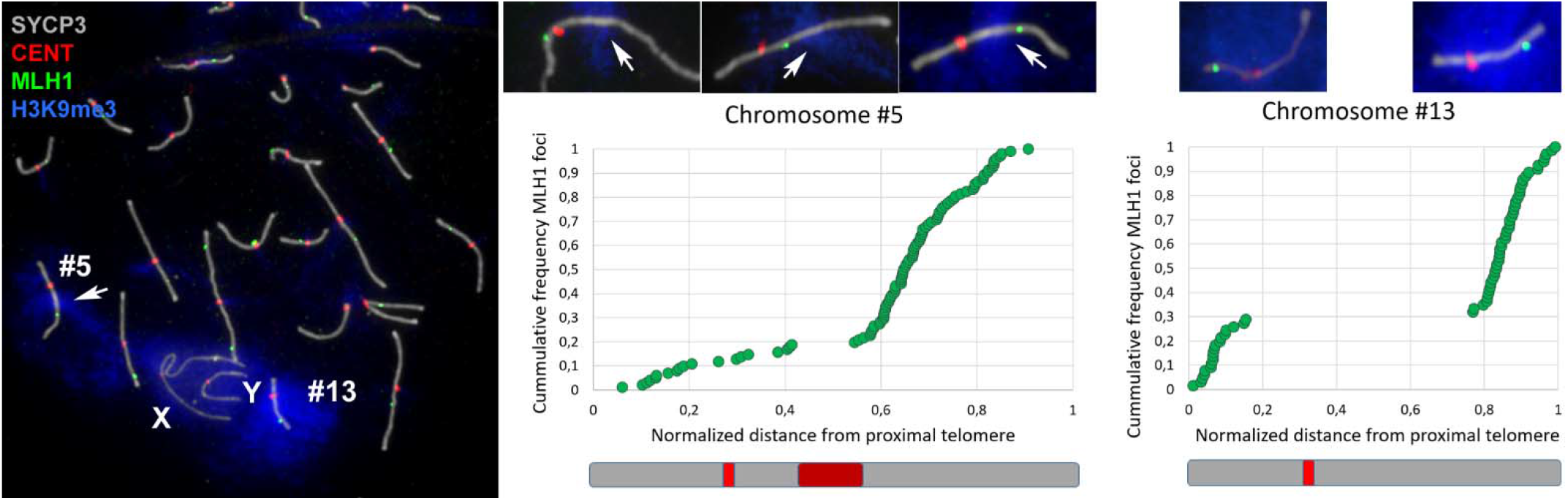
Distribution of recombination events in gerbil spermatocytes. (A) Immunolocalization of SYCP3 protein (grey) on meiotic chromosomes marks the trajectory of the synaptonemal complex along bivalents; trimethylation of histone H3 at lysine 9 (H3K9me3, blue) marks heterochromatin; CENT (red) stains centromeres; and MLH1 (green) marks the sites of crossovers. H3K9me3 is associated with the entirety of chromosome 13 (#13), a large intra-arm region of chromosome 5 (#5), and, to a lesser extent, the X and Y. The anti-CENT antibody (red) stains centromeres on all chromosomes but is not specifically associated with the large centromeric expansion of the long arm of chromosome 5. MLH1 foci can be located proximally, interstitially, or distally along bivalent 5 (central details, selected from three different spermatocytes), but they are never found within the centromere repeat expansion on this chromosome. Chromosome 13 shows either proximal or distal location of MLH1 foci (details on the right). (B) and (C) Graphs of MLH1 frequency against distance from the nearest telomere for bivalents 5 and 13, respectively. Each dot represents the location of the MLH1 focus along the synaptonemal complex on a single spermatocyte. The locations of centromeres and the chromosome 5 expansion are indicated as red and maroon boxes, respectively, on the schematic chromosomes below each graph. The graphs and drawings preserve the relative size of both chromosomes. For chromosome 5, most crossovers (over 80%) are located from the heterochromatic expansion towards the distal end. For chromosome 13, MLH1 foci are conspicuously accumulated towards the chromosomal ends, with an approximate 70:30 distribution on the long and short arms respectively.

### Chromosome 13: origin of a new autosome

Chromosome 13 is the most unusual chromosome in the gerbil genome for a variety of reasons. Karyotypically, it stains very dark and appears heterochromatic in G (Figure 3D) and C-banding images (Gamperl et al. 1977; Gamperl and Vistorin 1980). It also displays delayed synapsis during the first meiotic prophase, when compared to all other chromosomes (de la Fuente et al. 2007, 2014). On a technical level, it is the only chromosome that failed to assemble into a single chromosome-length scaffold (Figure 1), and even optical mapping was unable to improve the assembly. In a phylogenetic context there is no ortholog of chromosome 13 in mouse and rat, but similarity in G-banding patterns suggests that it may share ancestry with chromosome 14 in the fat sandrat (*P. obesus*). Short reads assigned to chromosome 13 have very low mapping quality as they map to multiple locations. As a result, chromosome 13 has very few genetic markers and a very short relative genetic map length compared to the other chromosomes (Table S1) and we suspect this is what prevented the OmniC data and HiRise pipeline from successfully assembling this chromosome. The centromere of chromosome 13 is unique in that the A-C arrays have more non-repetitive blocks interspersed within them than the other chromosomes (Figure 3D), and in terms of sequence complexity, there is no fine-scale variation in entropy across the chromosome (Figure 3, Figure S8) as on the other autosomes, suggesting very low sequence diversity. Indeed, the entropy of chromosome 13 appears even more homogenous than that of the Y chromosome (Figure 3, Figure S8). Chromosome 13 has more than the expected number of genes based on its size (Figure 6A), but far fewer unique genes (Figure 6B), demonstrating high levels of gene duplication: of the 1,990 genes on chromosome 13 annotated as something other than “Protein of unknown function”, 566 are copies of a viral *pol* protein (and so represent either endogenous retrovirus or LET retrotransposon sequences), 406 are Vmn2r (olfactory receptor) genes (of which 337 are copies of *Vmn2r116*) and 331 are Znf (Zinc finger) genes (257 of which are *Znf431*). There are more GC-rich genes located on chromosome 13 than expected based on its size (Figure 6C) and Chromosome 13 houses the original high-GC cluster (including the ParaHox genes) identified by (Hargreaves et al. 2017; Pracana et al. 2020). Chromosome 13 has a far higher repetitive sequence content (Figure 6D), as measured by the EarlGrey pipeline (Baril et al. 2022) which is clearly visible in comparison with other chromosomes in a self-alignment plot (Figure 5E-M). In fact, after filtering out alignments under 1,000bp, over 93% of bases on chromosome 13 are found in multiple copies on the chromosome, compared with ~10% on other autosomes (e.g. 11.5%, 8.2%, and 12.7% on the similarly sized chromosomes 10, 11, and 12 respectively). The bulk of chromosome 13 consists of around 400 copies of a block of DNA 170kb long, the periodicity and variable orientation of which can easily be seen in Figures 5H, 5I, and 5J. While we find no evidence of a link between high GC% genes and this chromosome generally, chromosome 13 does encode the set of genes previously identified as being the most extreme outliers in gerbil and sandrat genomes (Pracana et al. 2020). These genes surrounding the ParaHox gene cluster include *Pdx1, Cdx2, Brca2* and others crucial for proper embryogenesis and cell function (Withers et al. 1998). The cluster is contained within an ancient genomic regulatory block (Kikuta et al. 2007), where genes are locked together by the presence of overlapping regulatory elements. The presence of the most unusual genes on the most unusual chromosome is very interesting and is consistent with a model where the selective pressure to keep this block of genes intact may have had a role in the formation of the chromosome.

**Figure 6.**
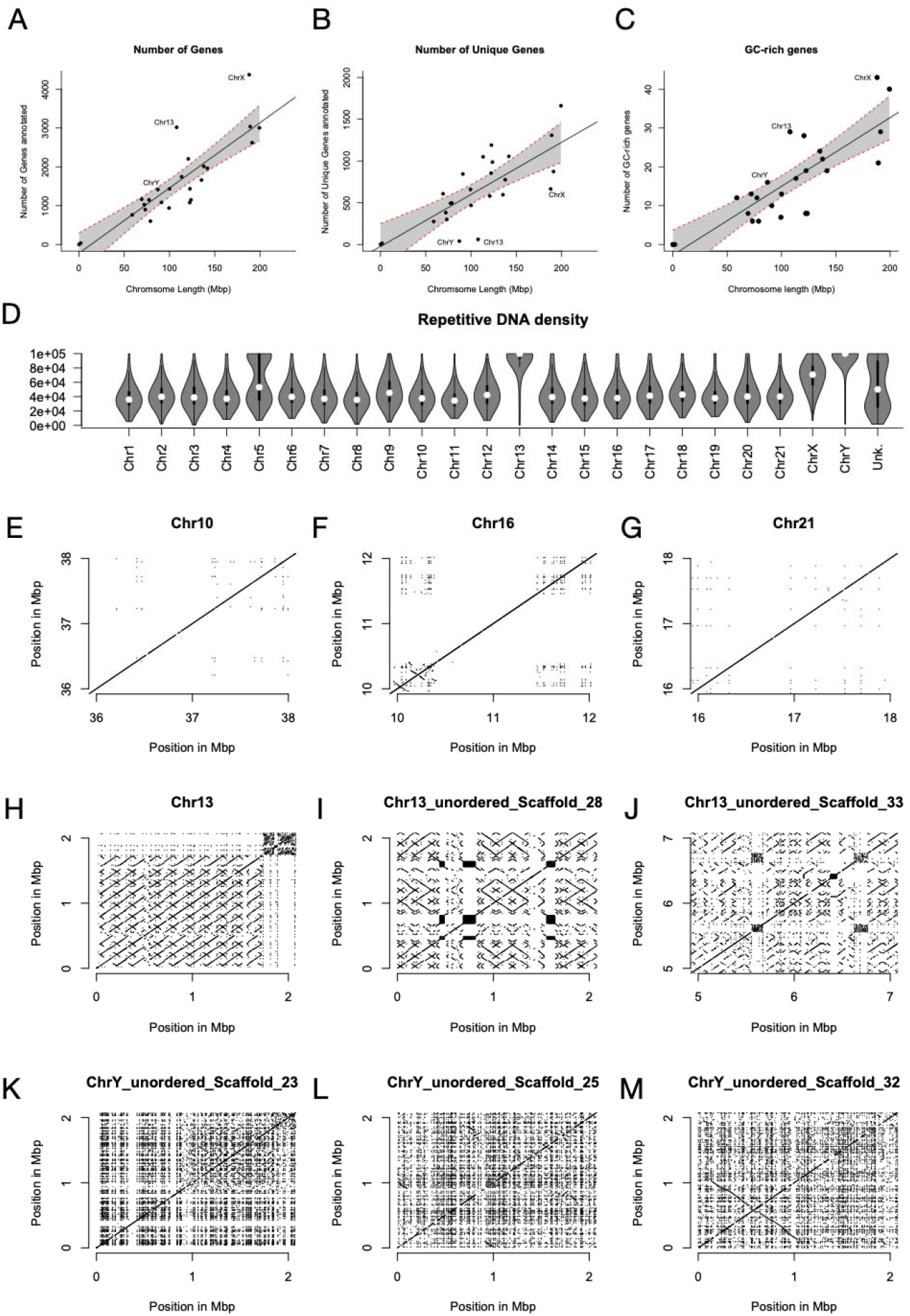
Chromosome 13 is unusual in terms of gene content and repetitive DNA density. (A) There is a strong relationship between chromosome length and gene number, but both chromosome 13 and the X have more genes than expected for their length. (B) When duplicate genes are removed, chromosome 13 and both sex chromosomes have far fewer genes than expected based on their length (error bars show the 95% confidence interval). (C) Chromosome 13 is enriched for GC-rich genes. (D) Chromosome 13 has far higher repetitive DNA content than the other autosomes and is rivaled only by the Y. Panels E-M show a self-alignment of a selection of “typical” chromosomes (E: Chr10; F: Chr16; G: Chr21), as well as three of the longer scaffolds from the highly repetitive chromosome 13 (H, I, J) and the Y (K, L, M). Each panel shows a 2Mbp section of chromosome and only alignments longer than 1,000 bases are plotted. The primary alignments are clearly visible as diagonal lines at y=x. All alignments off of the 1:1 line are repetitive sequence. The prevalence of repetitive sequence on chromosome 13 is much higher than other autosomes, and is most similar to the situation on the Y chromosome (D). However, repeats on chromosome 13 (H, I, J) are much longer than those on the Y (K, L, M), as expected based on their fundamentally different evolutionary history.

We propose the following model to explain the origin of chromosome 13: a chromosomal fragment approximately 5 million bases long which included the ParaHox cluster (Hargreaves et al. 2017) broke off from an ancestral chromosome, perhaps during a genome rearrangement.

The ParaHox genes and many of their neighbours are crucially important during development and so could not be lost altogether. For example: *Pdx1*-/- mice die shortly after birth (Jonsson et al. 1994; Offield et al. 1996) as do those lacking *Brca2* (Evers and Jonkers 2006) *Insr* (Accili et al. 1996) or *Hmgb1* (Calogero et al. 1999) function; *Cdx2*-/- mice die within the first 5 or 6 days of development (Chawengsaksophak et al. 1997); 75% of *Gsx1*-/- mice die within 4 weeks of birth, and none live beyond 18 weeks (Li et al. 1996); and *Flt1*-/- mice die *in utero* (Fong et al. 1995). These are just a small selection of genes in this region, but they demonstrate the selective pressure(s) that must exist for its maintenance within the genome. While the simplest option might have been for this fragment to have joined onto or into another chromosome, this does not appear to have happened, and instead we propose that this chromosomal fragment became the seed for the growth of an entirely new chromosome. In some species, the evolutionary fate of such a fragment may be long-term persistence as a microchromosome: a small, gene-dense, repeat-poor, GC-rich chromosome of ≤30Mb with a high recombination rate. But while microchromosomes are common in birds, reptiles, and fish, they do not persist in mammals over evolutionarily time (Srikulnath et al. 2021; Waters et al. 2021). Efficient transmission of mammalian chromosomes between generations and into daughter cells therefore seems to require a minimum size, and in the case of *M. unguiculatus* chromosome 13, we propose the hypothesis in which the fragment grew rapidly via a breakage-fusion-bridge mechanism, (McClintock 1938, 1941; Bignell et al. 2007; Campbell et al. 2010; Greenman et al. 2012), where the chromatid ends without a telomere fuse, and then are pulled apart at anaphase, breaking randomly and resulting in long inverted repeats. The patterns apparent in the Chromosome 13 self-alignments (Figure 6G, 6H, 6I) are consistent with what would be expected under this model and may explain how a 170kb region at the end of the chromosome was repeatedly duplicated, at multiple scales, until a 107Mb chromosome was formed. The high similarity of these duplicated regions explains our difficulty in assembling this chromosome, the multimapping of short reads, and the failure of BioNano optical mapping to improve our assembly.

While the repetitive nature of Chromosome 13 is consistent with it arising and evolving under the model described above, that does not explain why Chromosome 13 houses genes with the most extreme GC content. Previous authors (Gamperl and Vistorin 1980) have described that chromosome 13 forms ring-like structures during meiosis, suggesting that the bulk of the heterochromatic material on this chromosome does not, or possibly cannot, form chiasma, and therefore cannot undergo recombination. However, based on localization of the recombination marker MLH1, we have found evidence of recombination during male meiosis (Figure 5). Bivalent chromosome 13 presents a recombination event in most spermatocytes, although a small proportion (around 23%) lack MLH1 foci. Strikingly, MLH1 are not evenly distributed along this chromosome, as previously reported for other chromosomes (de la Fuente et al. 2014). Instead, recombination events are strongly concentrated at the chromosome ends. We therefore propose that the extreme GC skew of the ParaHox-associated genes in gerbils is the result of the inability of recombination hotspots to move out of this genomic region, leading to runaway GC-bias.

## Conclusion

The two heterochromatin-rich chromosomes of Mongolian gerbils have distinct origins. Chromosome 5 has undergone a massive expansion of a centromeric repeat, most likely as a result of meiotic drive, and Chromosome 13 has likely arisen *de novo* from an initially small seed via multiple breakage-fusion-bridge cycles. In general, these results show the importance of karyotypic knowledge of study species and serve as a warning for large-scale genome sequencing programs such as the Vertebrate Genomes Project (VGP) or the Darwin Tree of Life Project (DToL) that we must not neglect knowledge of chromosome number and morphology. Had we not known the diploid chromosome number for *M. unguiculatus*, and had we not performed chromosome sorting and FISH, we likely would have binned the 121 fragments corresponding to chromosome 13 into the “unknown” category and deduced that gerbils had one fewer chromosome than they actually have. We applied what are becoming the standard approaches for genome sequencing and assembly to the *M. unguiculatus* genome (PacBio HiFi, chromatin conformation capture, Oxford Nanopore long reads, and Bionano optical mapping), and incorporated chromosome sorting, FISH, and a SNP-based linkage map, and were still unable to assemble chromosome 13 into a single scaffold. The huge size and high similarity of the chromosome 13 repeats suggest that only ultra-long Oxford Nanopore reads, on the order of several hundred kilobases, might be able to achieve the telomere-to-telomere coverage of this enigmatic chromosome.

## Materials and Methods

The complete details of the methods are available at the end of Supplemental Material 1, here follows a brief overview.

For sequencing and assembly, we extracted DNA from gerbil liver and sequenced to a depth of 34X using PacBio HiFi technology. Genome assembly was done with the program HiFiAsm (Cheng et al. 2020). Scaffolding was done using a combination of Dovetail OmniC, Oxford Nanopore Ultra-long sequencing, Bioano Optical Mapping and a genetic map from (Brekke et al. 2019). The genome was annotated using RNAseq from kidney and testis from three individuals. Repeats were annotated using the EarlGrey pipeline (Baril et al. 2022).

For cell culture, chromosome sorting, and FISH we cultured cells from the gerbil fibroma cell line IMR-33 and chromosomes extracted for cell sorting after being arrested in mitosis.

Chromosome sorting was done with a BD Influx Cell sorter into the 17 pools containing one or two chromosomes. Each pool was sequenced with Illumina MiSeq. FISH paints were made from each pool as well.

For mitotic chromosome preparation and FISH we cultured fresh spleen cells which were then arrested in mitosis for chromosome spreads. These were stained with DAPI and the FISH probes derived from the chromosome sorting and visualized on a confocal microscope.

For meiotic chromosome preparation and immunofluorescence, we extracted meiotic cells from fresh testis and processed them for spreads and immunofluorescence. Slides were incubated with the primary antibodies goat anti-SYCP3 to mark the synaptonemal complex, rabbit anti-histone H3 trimethylated at lysine 9 to mark heterchromatin, human anti-centromere, and mouse anti-MLH1 to mark meiotic crossovers. Then slides were incubated with secondary antibodies and visualized with an Olympus BX61 microscope equipped with appropriate fluorescent filters and an Olympus DP72 digital camera.

We assigned the sequenced scaffolds with the chromosomes in the karyopye by aligning the reads from the sequenced pools to the scaffolds and identifying which pools’ reads most often aligned to each scaffold. Then we linked the pools to the karyotype by staining mitotic chromosome spreads with the FISH probes derived from each pool.

We calculated GC content and gene density for each chromosome in sliding windows of size 1kb and 1Mb respectively with step size 1kb. We calculated recombination rate with a sliding window of 8 markers with a step of 1 marker and regressed marker position against physical position. Hotspots were identified as a region whose recombination rate was 5x the genome average. Entroy and Linguistic complexity were calculated with the program NeSSie (Berselli et al. 2018) using a sliding window of size 10kb with a step of 1kb.

Centromeres were located at the trough of the linguistic complexity plot and the fine-scale structure was analysed with NTRprism (Altemose et al. 2022) and TandemRepeatFinder (Benson 1999). Interstitial telomeres were identified as those with >70 copies of the telomere sequence in the TandemRepeatfinder data. Self-alignments were done with mummer (Kurtz et al. 2004).

## Supporting information

Supplemental material 2 - genome sketch

## Acknowledgements

The authors would like to thank two anonymous reviewers and the editor for their comments as well as Aaron Comeault, Martin Swain, Yichen Dai, Adam Hargreaves, Peter Holland, and Roddy Pracana for helpful discussions pertaining to the project, and Becca Snell for help with animal care. Also David Thybert for providing the *Psammomys* genome assembly. TDB would like to thank Kris Crandell. This work was supported by the Leverhulme Trust grant entitled “Decoding Dark DNA” (grant number RPG-2018-433) and by the National Environmental Research Council of the UK (grant number NE/R001081/1 to A.S.T.P) and by grant CGL2014-53106-P from Ministerio de Economía y Competitividad (Spain to J.P.).

Unpublished genome assemblies for *Meriones unguiculatus* are used with permission from the DNA Zoo Consortium (<u>dnazoo.org</u>).

## Authors contributions

O.F. and E.J.– chromosome sorting, editing manuscript

F.Y. and B.F. – FISH, editing manuscript

T.B. and A.H. – EarlGrey, repeat annotations, editing manuscript

J.P and R. d. l. F. – recombination/histone analyses, editing manuscript

T.D.B., J. F. M., and A. S. T. P. – conceived study, genome sequencing, assembly, analysis, overall coordination, writing and editing manuscript

## Data and materials availability

All sequencing data and the genome is available under SRA BioProject PRJNA397533. Specific accession numbers can be found in Supplemental Material 1.

