## Supplemental Material 1 for "A chromosome-assigned Mongolian gerbil genome with sequenced centromeres provides evidence of a new chromosome"

**List of Supplementary materials:**
Supplemental Material 1 (this file): Figures, Tables, and Methods

Figure S1: C-banding and Dapi-banding karyotypes of *Meriones unguiculatus.* Page 3

Figure S2: Dovetail OmniC contact map. Page 4.

Table S1: Genetic map statistics: Page 5.

Methods for chromosome sorting and the logic linking the fasta records to the karyotype. Pages 6-23. Including Figures S3-S7, and Tables S2 and S3.

Figure S3: Flow karyotype of the Mongolian gerbil. Page 8.

Table S2: Gerbil chromosomes in each sorted pool. Page 10.

Figure S4: Gerbil FISH probes hybridized to gerbil chromosomes. Page 11-16.

Figure S5: Flow chart showing the logic of assigning chromosomes to scaffolds. Page 18.

Figure S6. Comparative DAPI-banding between sandrat and gerbil. Page 19.

Figure S7. FISH images of mouse chromosome paints on sandrat chromosomes. Page 20.

Table S3: Summary of gerbil/sandrat/mouse chromosome homology. Page 21.

Table S4: Comparison of three published gerbil genomes. Page 24.

Figure S8: GC content, gene density, entropy, and linguistic complexity in sliding windows across each chromosome. Page 25-48.

Figure S9: Whole-genome alignment of the genome presented here and Cheng et al (2019)’s HiC scaffolded version. Page 49.

Figure S10: Repeat density plots of centromeres. Page 49-57.

Figure S11: Recombination rates for each chromosome showing hotspots. Page 58.

Figure S12: Marey maps for each chromosome showing hotspots. Page 59.

Figure S13: Recombination hotspot distribution across chromosomes. Page 60.

Figure S14: GC rich outlier genes occur closer to recombination hotspots than expected by chance. Page 61.

Figure S15: GC rich outlier genes are found closer to telomeres than expected by change. Page 62.

Figure S16. Interstitial telomere repeat array lengths. Page 63.

Figure S17: GC rich genes are closer to interstitial telomeres than expected by chance. Page 64.

Figure S18: Schematic of centromere location, gc rich genes, recombination hotspots, and interstitial telomeres. Page 65.

Figure S19: A whole genome alignment of *Meriones* *unguiculatus* chromosome 5 with *Psammomys* *obesus* chromosome 10 showing the expansion of *Meriones* *unguiculatus* Chr5. Page 66.

Detailed Methods. Page 67-76.

Data availability. Page 77.

References. Page 78-81.

Supplemental Material 2: A very high-resolution image of Figure 3D to facilitate close inspection of chromosomal features.

Supplemental Material 3: Code base. All in-house scripts and dataframes, including vcf, gff, and genetic map files. See do_it_all_genome_polish_v6.sh for a step-by-step call sequence. Also includes some metadata files needed to run the pipeline and some Rdata packets to skip some time-consuming analyses. This supplement can be found in the Dryad repository here: <https://doi.org/10.5061/dryad.1vhhmgqws>

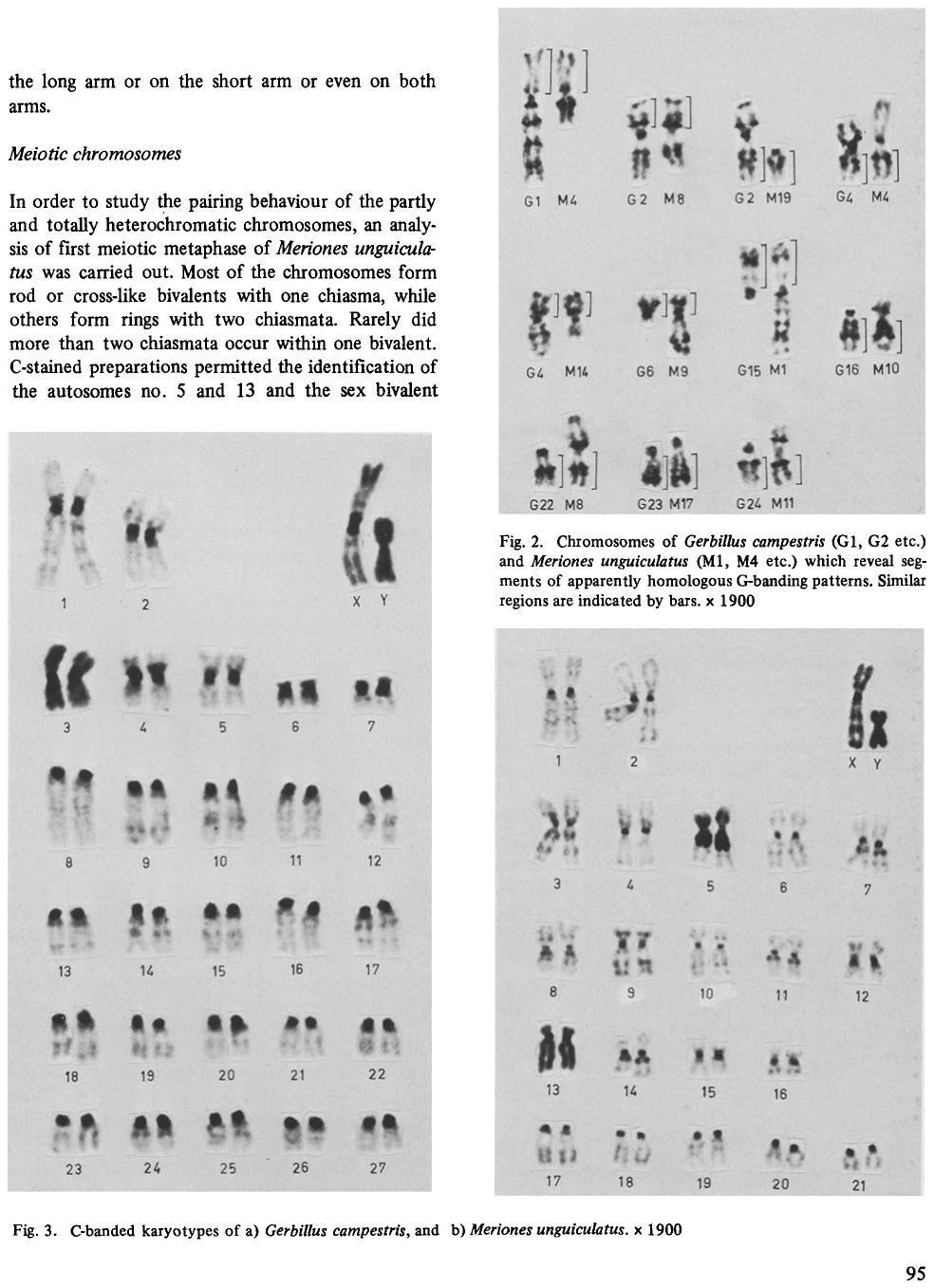
Figure S1A: C-banding karyotype of *Meriones unguiculatus*. Dark bands show heterochromatic regions, which are most often centromeres. Originally published as “Figure S3B” in (Gamperl and Vistorin 1980), and reprinted here with permission. Original caption reads: *“Fig. 3. C-banded karyotypes of a)* Gerbillus campestris*, and b)* Meriones unguiculatus*. x 1900”*. Note the dark-staining of the sex chromosomes, much of chromosome 5, and all of chromosome 13, which indicates heterochromatin, while only centromeric regions are stained in all other chromosomes.

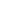

Figure S1B: DAPI-banding karyotype of *Meriones unguiculatus*. Mongolian gerbil DAPI-banding karyotype. Gerbils have 21 autosomes and an X-Y sex determination system.

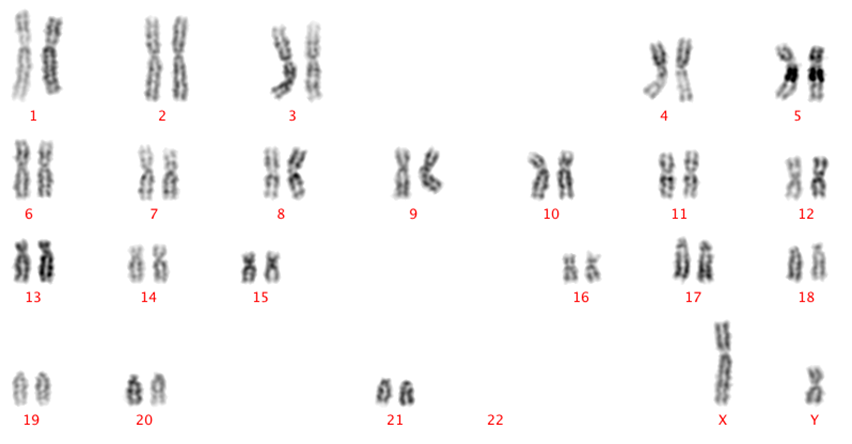

Figure S2: Dovetail OmniC contact map. This was the an early stage in the scaffolding pipeline. Further merges of scaffolds not shown here were done after this step using Oxford Nanopore ultra-long reads and a genetic map and optical mapping data.

The reason that OmniC reads were unable to scaffold Chromosome 13 is that all the reads that mapped to Chr13 mapped multiple times and so had very poor mapping quality and were removed from the analysis in the HiRise pipeline.

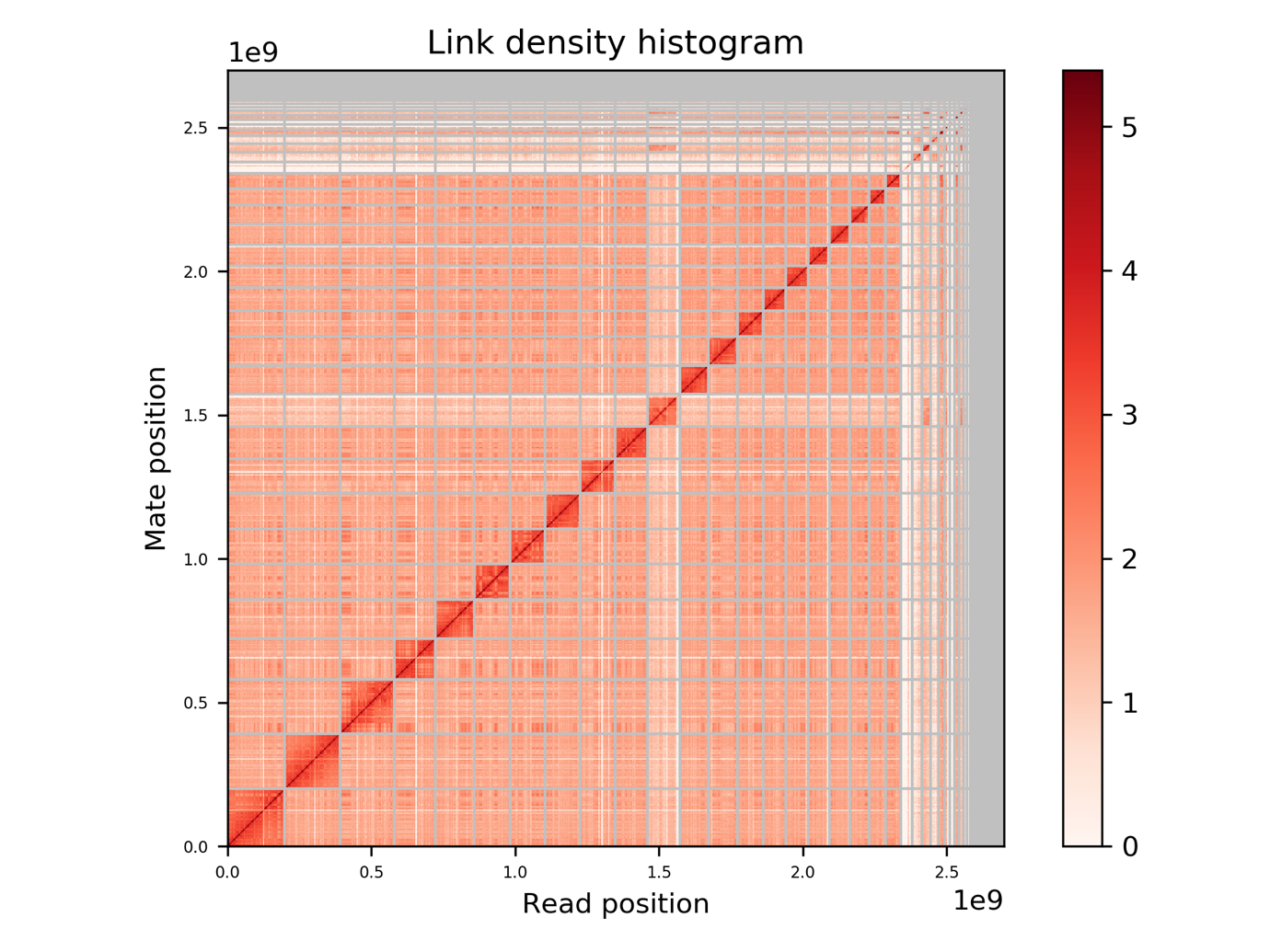

Table S1: Genetic map statistics

| Linkage Group | Number of Markers | Total centiMorgans |
| --- | --- | --- |
| 1 | 89 | 380.9 |
| 2 | 173 | 211.4 |
| 3 | 290 | 315.1 |
| 4 | 170 | 185.6 |
| 5 | 72 | 96.5 |
| 6 | 141 | 276.7 |
| 7 | 173 | 271.9 |
| 8 | 190 | 268.7 |
| 9 | 164 | 290.9 |
| 10 | 26 | 72.0 |
| 11 | 127 | 168.9 |
| 12 | 133 | 139.8 |
| 13 | 6 | 3.4 |
| 14 | 95 | 135.6 |
| 15 | 91 | 104.2 |
| 16 | 77 | 66.8 |
| 17 | 140 | 147.9 |
| 18 | 34 | 53.6 |
| 19 | 122 | 109.7 |
| 20 | 124 | 190.9 |
| 21 | 18 | 21.1 |
| X | 37 | 64.3 |
| Total | 2492 | 3575.7 |

Original data published in (Brekke et al. 2019)

**Using Chromosome sorting and FISH to link fasta records with karyotypes via sorted pools: a logical puzzle**

**Introduction**

Goal: Link the scaffolds in the reference fasta of the *Meriones* *unguiculatus* genome with a specific chromosome image and banding pattern in the *Meriones unguiculatus* karyotype.

Approach: Flow-sort *Meriones* chromosomes and make chromosome-specific FISH probes to paint chromosome spreads, thereby linking the sorted pools to the chromosome banding pattern for each chromosome. Then sequence the pools and align the reads to the scaffolds to link the pools to the scaffolds in the reference fasta.

Mongolian gerbils have 21 autosomes, an X, and a Y chromosome (Figure S1) and these were sorted into 17 pools (Figure S3).

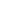

**Methods**

**BIVARIATE FLOW KARYOTYPING FOR CHROMOSOME SORTING OF GERBIL FIBROBLASTS IMR-33 CELL LINE: Chromosome suspension preparation**

*Meriones* *unguiculatus* chromosome suspensions were prepared based on the method described by (Kuderna et al. 2019), using the classic method of Chromosomes double stained with Hoechst 33258 (AT-bp specific dye) and A3-Chromomycin (CG-bp specific dye).

Briefly, cells were cultured in standard culture conditions and near the 50% of confluence Colcemid was added to the cell culture at final concentration of 0.1 µg/ml for 6 hours to block cells in mitosis. After incubation with Colcemid, mitotic cells were harvested by shaking the flask and tapping the side of the flask firmly. Cell suspension was transferred to a tube to be centrifuged for 5min at 300xg at RT.

To swell and stabilize mitotic cells, the cell pellet was disturbed by flicking the tube and resuspended in hypotonic solution (HPA solution:75mM KCl, 10mM MgSO_4_.7H_2_O, 0.2 mM spermine and 0.5mM spermidine) pH 8.0 and incubated for 20 min at 37°C.

Swollen cells were centrifuged for 5 min at 300 × g at RT. To release the chromosomes, the cell pellet was disturbed by flicking the tube once resuspended in 1ml of ice-cold polyamine isolation buffer pH 8.0 (PAB buffer: 15mM Tris pH 8.0, 2mM EDTA, 0.5mM EGTA, 80mM KCl, 3mM DTT, 0.25% Triton X-100, 0.2mM Spermine tetrahydrochloride, 0.5mM Spermidine trihydrochloride) and incubated for 40 min on ice. Then, sample was vortex vigorously for 30 seconds to liberate chromosomes from mitotic cells. The vortex time is vortex force and cell type dependant.

Chromosome staining was performed adding Hoechst 33258 (final concentration, 5 µg/ml) and chromomycin A3 (final concentration, 40 µg/ml) in presence of divalent cations (MgSO4.7H2O 10mM) and incubated for at least 8 hours at 4°C. Finally, to stabilize the staining, sodium citrate (final concentration, 10mM) and sodium sulphite (final concentration, 25mM) per ml of chromosome suspension was added and incubated for at least 2 hours on ice before analysis on the sorter.

**BIVARIATE FLOW KARYOTYPING FOR CHROMOSOME SORTING OF GERBIL FIBROBLASTS IMR-33 CELL LINE: Chromosome sorting**

Flow karyotyping and chromosome sorting was performed and described by (Kuderna et al. 2019), using a BD Influx cell sorter (Becton Dickinson, San Jose, CA) with 100 μm nozzle configuration.

For A3-Chromomycin and Hoechst-33258 detection, deep-blue laser (457nm at 300 mW) and UV laser (355 nm laser at 100 mW) were used respectively. Blue laser (488nm 200 mW) was only used for primary instrument alignment with Rainbow beads.

Chromomycin-A3 fluorescence was detected through a 500 nm long pass (LP) filter and collected by a 550/50 nm bandpass filter, while Hoechst-33258 was detected through a 400-LP filter and by 460/50 nm bandpass filter.

Instrument optimization, setup and performance were optimized by using standard 8-peaks Rainbow beads (SpheroTM Rainbow Calibration Particles) for blue laser, 1-peak UV beads (AlignflowTM Flow Cytometry Alignment) for UV laser and 1-peak 457nm (FluoresbriteTM Plain YG Microspheres) for deep-blue laser.

The threshold for chromosome sorting was set triggering in chromomycin-A3 fluorescence on 457nm. All parameters were collected in lineal mode and analyzed with the BD FACSTM Software (v. 1.0.0.0.650, Becton Dickinson, San Jose, CA). Bivariate flow karyotyping was directly setup in a biparametric dot-plot of Chromomycin-A3 versus Hoechst-33258 (Figure S3).

After chromosomes were sorted, we dialysed each pool using a Pur-A-Lyzer Maxi Dialysis kit (Sigma, PURX50005-1KT) following the manufacturer's instructions to remove any dye that remained bound to the DNA from sorting.

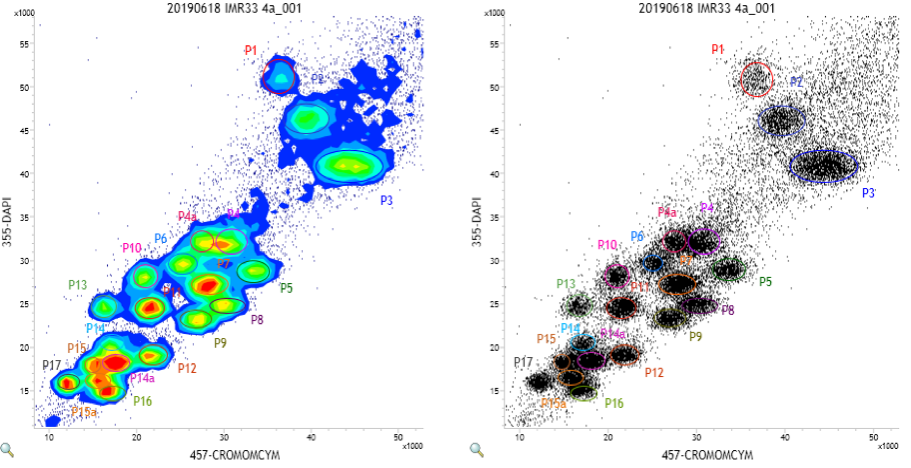

Figure S3: Flow karyotyping for Mongolian gerbil chromosomes into 17 pools labelled P1 to P17. Pools with a letter were combined with the pool of the same number (i.e. P4 and P4a were combined) for FISH probes and sequencing. Left and right panels are the same data, just plotted as a heatmap (left) of points (right). The threshold gates were set based on visual inspection of these plots.

**Sequencing of the sorted pools**

We sequenced each pool with an Illumina. MiSeq reads were aligned to scaffolds of the reference fasta with bwa (Li and Durbin 2009) to link fasta records with pools. For the alignment of each pool, we counted the number of reads mapping to each scaffold and calculated the reads mapped per scaffold length. Each scaffold then has a read-mapping density from each pool making it possible to associate every scaffold with the pool to which it belonged. We calculated the 99.99% confidence interval of the read mapping density and the scaffold was assigned to the pool that fell outside the confidence interval.

**Chromosome-specific paint probes**

We made FISH probes by first doing a whole-genome amplification of each chromosome pool using a whole genome amplification (WGA) kit (Sigma, WGA2) following the manufacturer's instructions. The amplified products were labelled using a WGA reamplification kit WGA3 but using home-made dNTP mixture with either Atto-488 or Texas red dUTP as described in(Murchison et al. 2012). Fluorescence in situ hybridisation, image capturing, and processing were the same as Murchison et al. (2012) and (Yang et al. 2017). DAPI-banding karyotypes were compared with published karyotypes (Weiss et al. 1970) to identify the chromosomes.

**Mitotic Chromosome spreads**

Complete spleens were harvested from euthanised animals and spleen cells were cultured in complete RPMI with 10% FCS and 1% Pen-strep. LPS from *E. coli* was used to stimulate the growth of B cells. Cell culture was incubated for 48 hours at 37c in a 5% CO_2_ incubator. Cells were arrested at mitotic metaphase with colcemid (final concentration of 0.1µg/mL) and incubated for a further 90 minutes. The cell culture was then spun down and resuspended in 0.075M KCl and further incubated in at 37c for 12 minutes at which point a few drops of fixative (3:1 methanol: glacial acetic acid) were added. Cells were spun down again, resuspended in fixative and stored at -20c.

Chromosome spreads for all species were made by dropping 20ul of fixed cell culture onto a cleaned slide followed by a drop of fixative, incubated in a humid 55c water bath, washed 10 min in acetone, baked 30 minutes at 65c, washed 1 minute in 50:50 mix of 0.5M NaOH and 1.0M NaCl, 1 minute in Tris HCl (pH 7.5), and dehydrated in an ethanol series: 2 minutes each of 70%, 70%, 90%, 100% EtOH.

**Results**

**Assigning chromosomes sequences with karyotypes**

We sorted 23 chromosomes into 17 pools, with 11 pools having a single chromosome and 6 pools having two chromosomes (Figure S3 and Table S2). We linked the karyotype with the fasta record by sequencing the pools and aligning the reads to the genome. We were thus able to identify the scaffold(s) that associated with each pool by comparing the depth of coverage on each scaffold. Then we used the pools to create FISH probes and painted a chromosome spread with each pool’s paint (Figure S4). By comparing the DAPI-banding of the chromosome(s) that lit up with each pools paint with the banding pattern of the original karyotype, we were able to link pools with Chromosome names. These results are described in table S2.

Table S2: Gerbil chromosomes in each sorted pool

| Pool | Chromosome |
| --- | --- |
| P1 | X |
| P2 | 2 |
| P3 | 1, 3 |
| P4 | 5, 6 |
| P5 | 4 |
| P6 | 9 |
| P7 | 8, 10 |
| P8 | 7 |
| P9 | 11 |
| P10 | 13 |
| P11 | 12, 17 |
| P12 | 14 |
| P13 | Y |
| P14 | 15, 18 |
| P15 | 19, 20 |
| P16 | 16 |
| P17 | 21 |

Figure S4: Gerbil FISH probes hybridized to gerbil chromosome spreads.

Figure S4A: Gerbil pool P1 hybridized to the gerbil X Chromosome. There is also some cross-hybridization to Chr13 likely due to its repetitive nature.

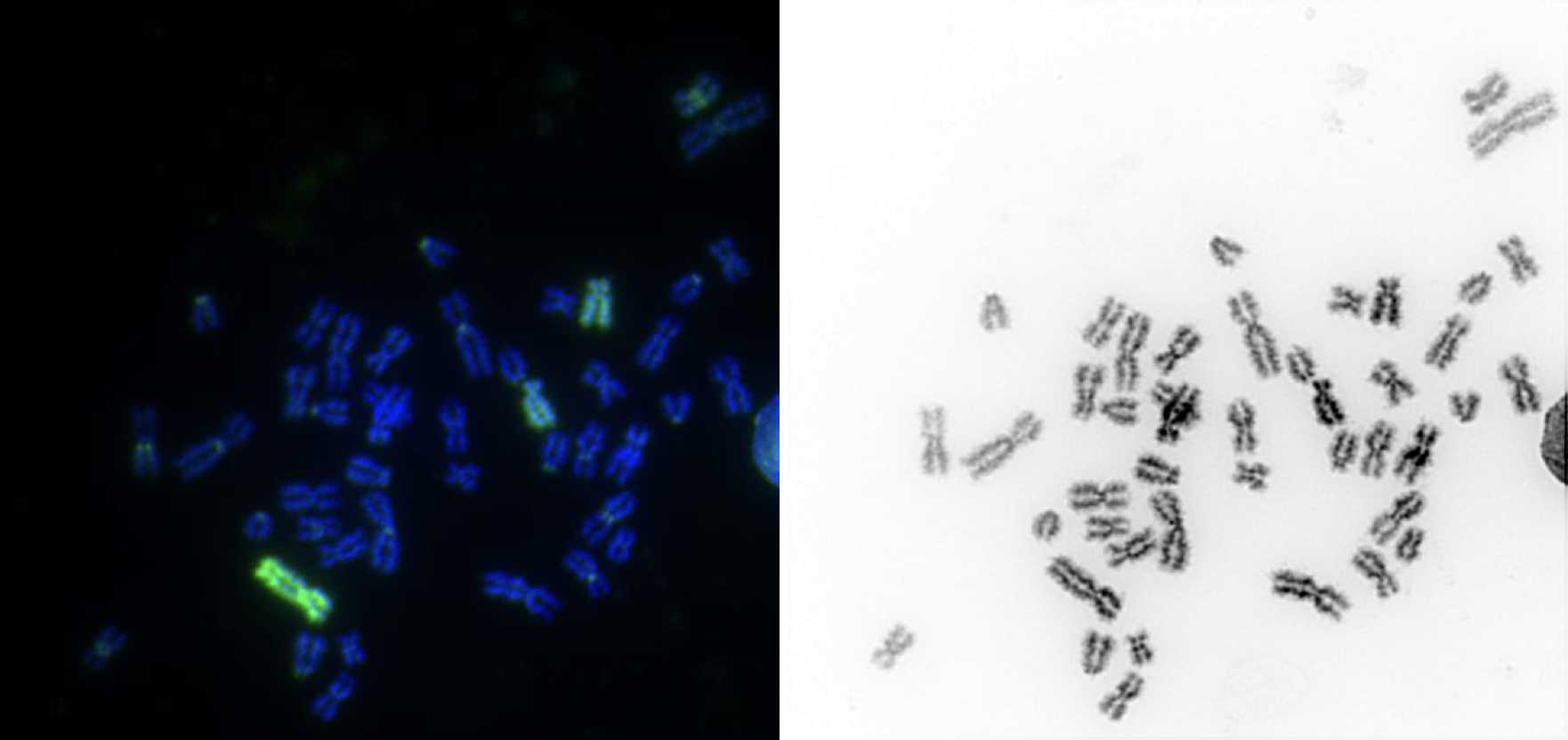

Figure S4B: Gerbil pool P2 hybridized to gerbil Chromosome 2 and also to gerbil Chromosome 13. We suspect that Chromosome 13 dimerized in the hypotetraploid cell line used for sorting and some of these molecules sorted out among Chromosome 2. This would explain why Chromosome 13 shows up here along with Chromosome 2 but why Chromosome 2 does not show up along with Chromosome 13 in P10. Alternatively, this could be due to cross-hybridization of shared repeats.

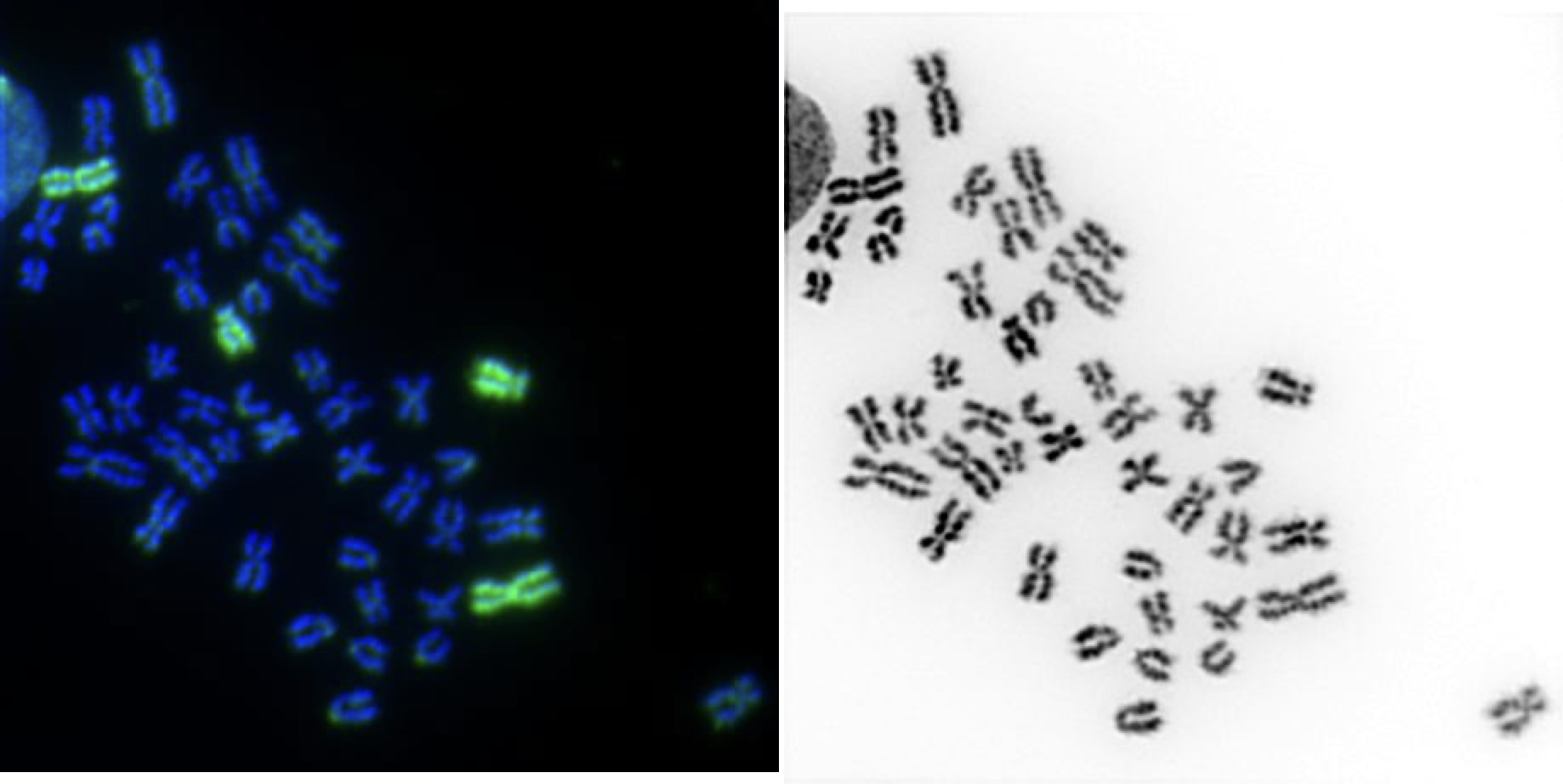

Figure S4C: Gerbil pool P3 (red) hybridized to gerbil Chromosomes 1 and 3. Also Pool 6 (green) hybridized to Chromosome 9.

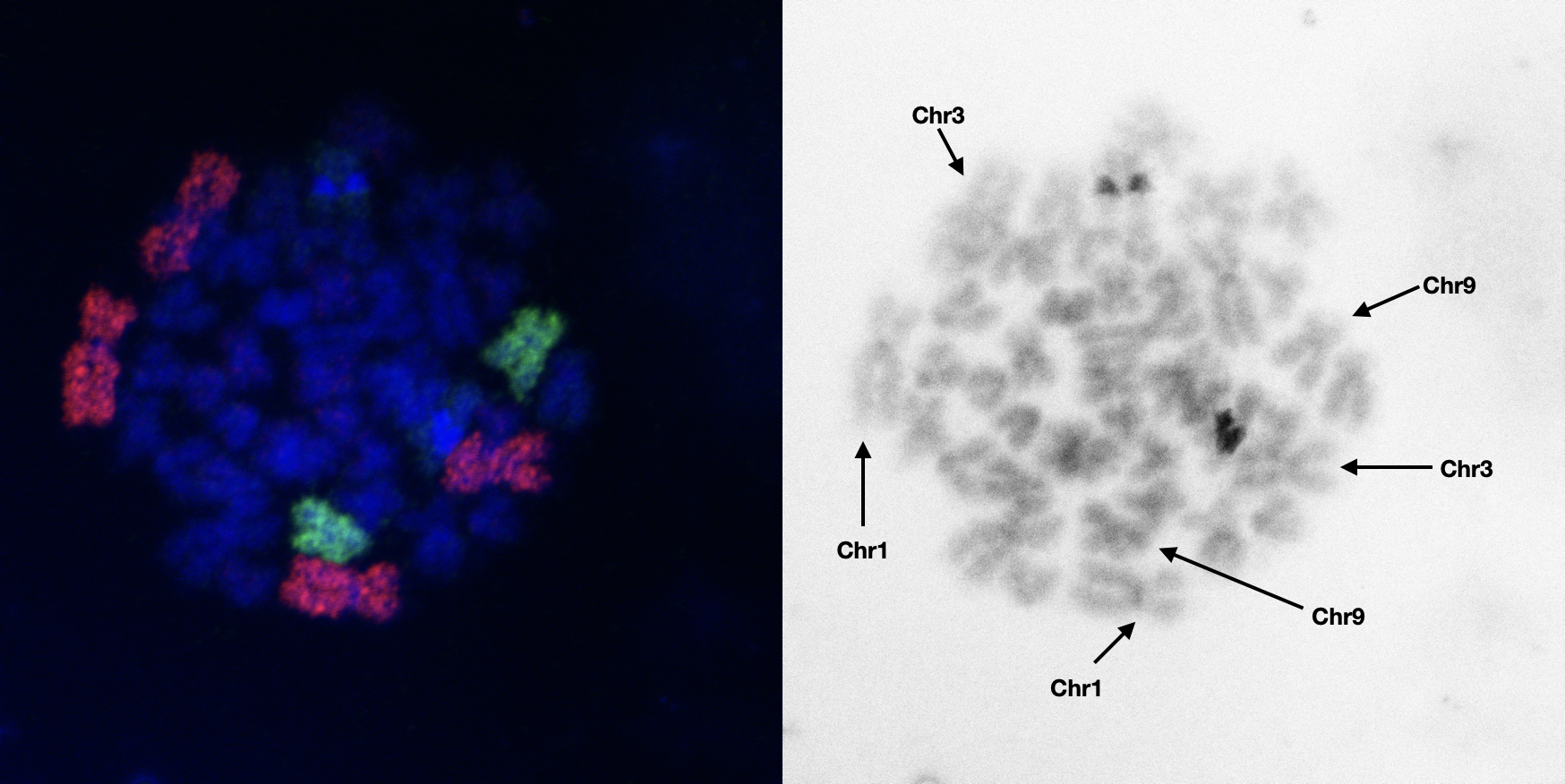

Figure S4D: Gerbil pool P4 hybridized to gerbil Chromosome 5 and 6.

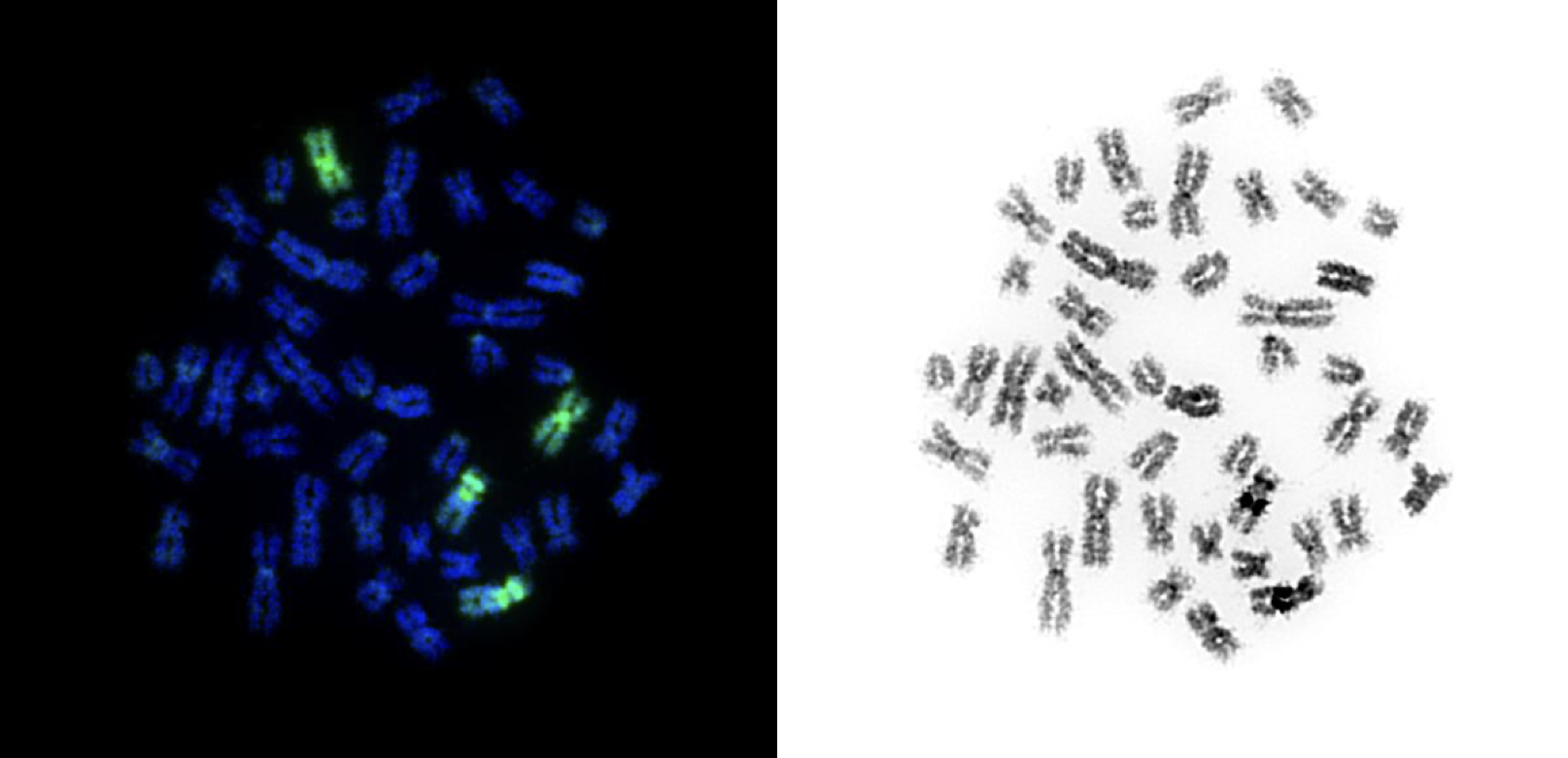

Figure S4E: Gerbil pool P5 (red) hybridized to gerbil Chromosome 4. Also gerbil P8 (green) hybridized to Chromosome 7.

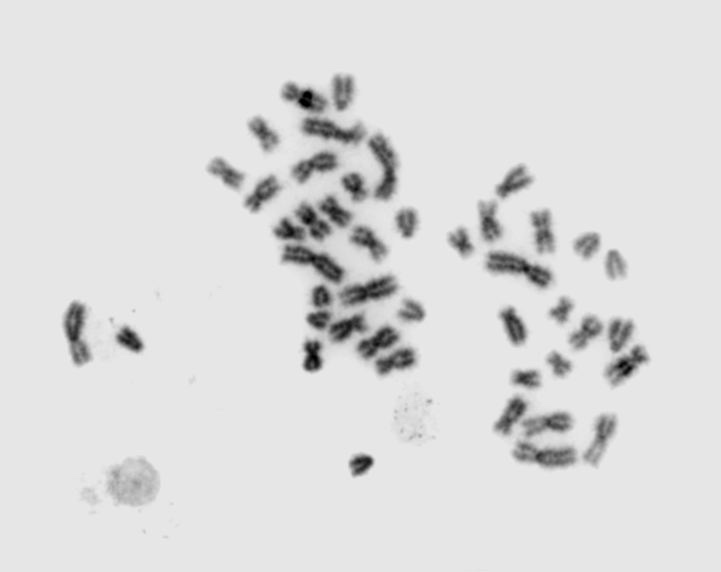

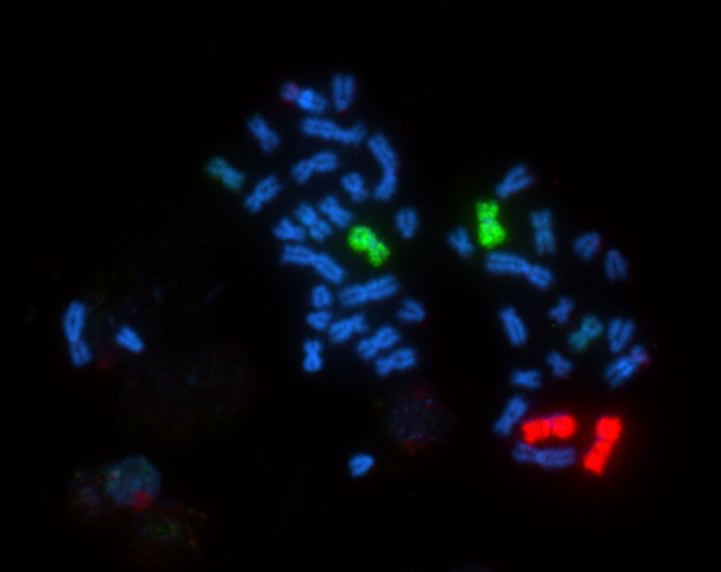

4

12

12

Figure S4F: Gerbil pool P7 hybridized to gerbil Chromosomes 8 and 10.

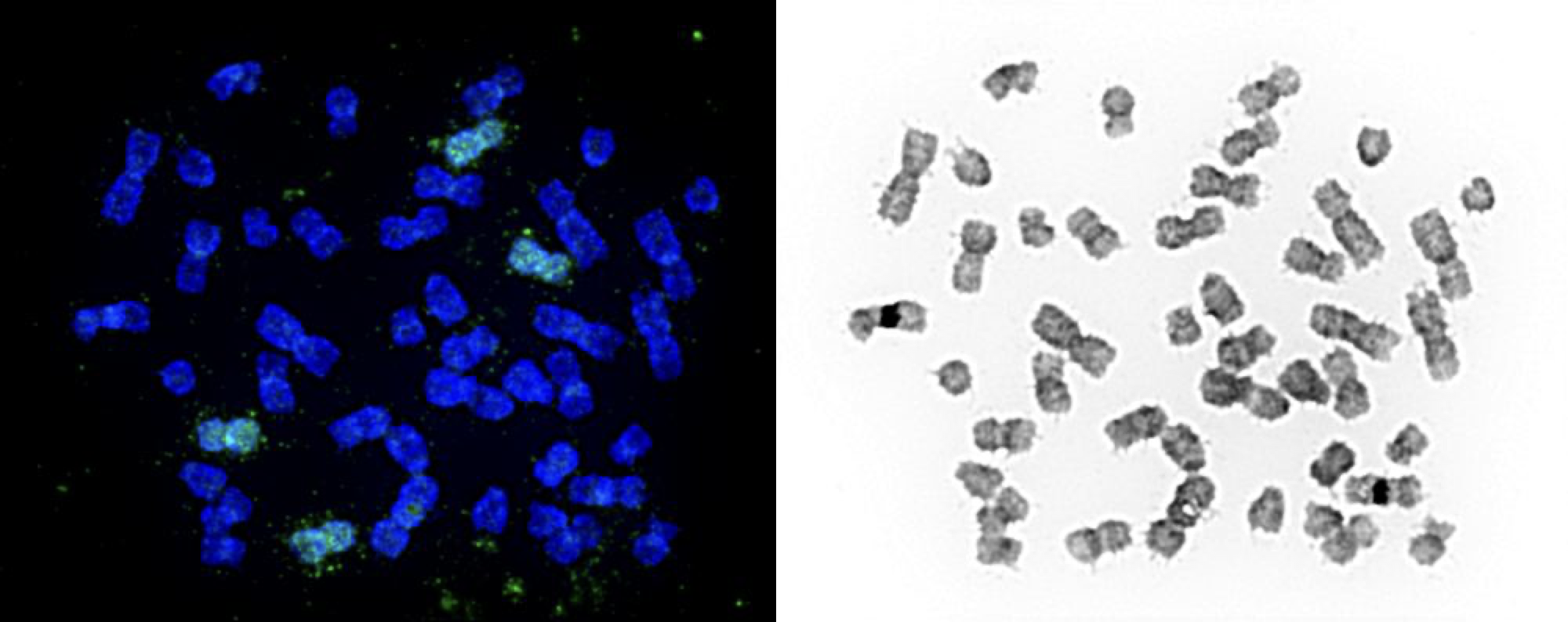

Figure S4G: Gerbil pool P9 (red) hybridized to gerbil Chromosome 11. Also Pool 12 (green) hybridized to Chromosome 14.

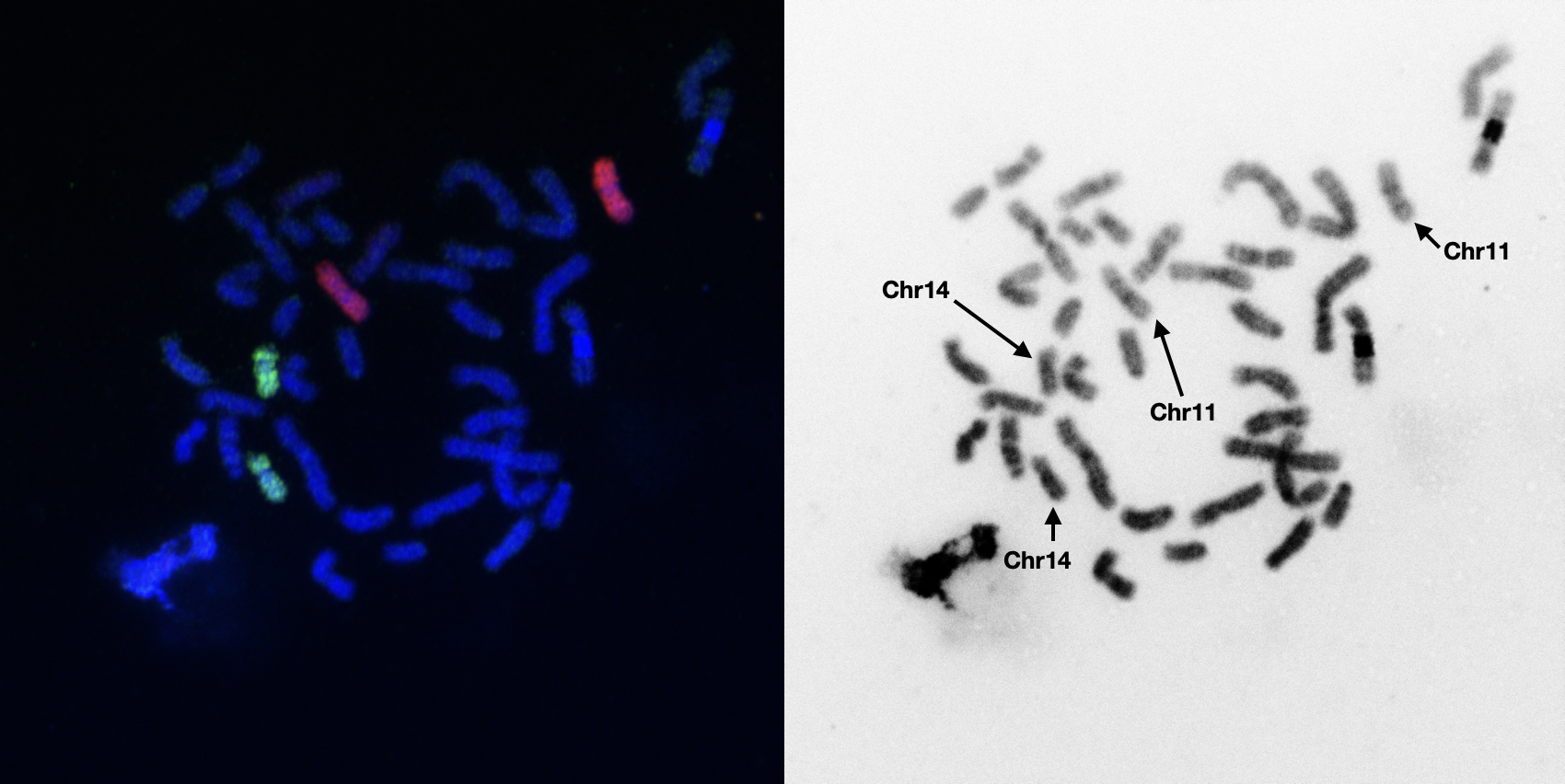

Figure S4H: Gerbil pool P10 hybridized to gerbil Chromosome 13.

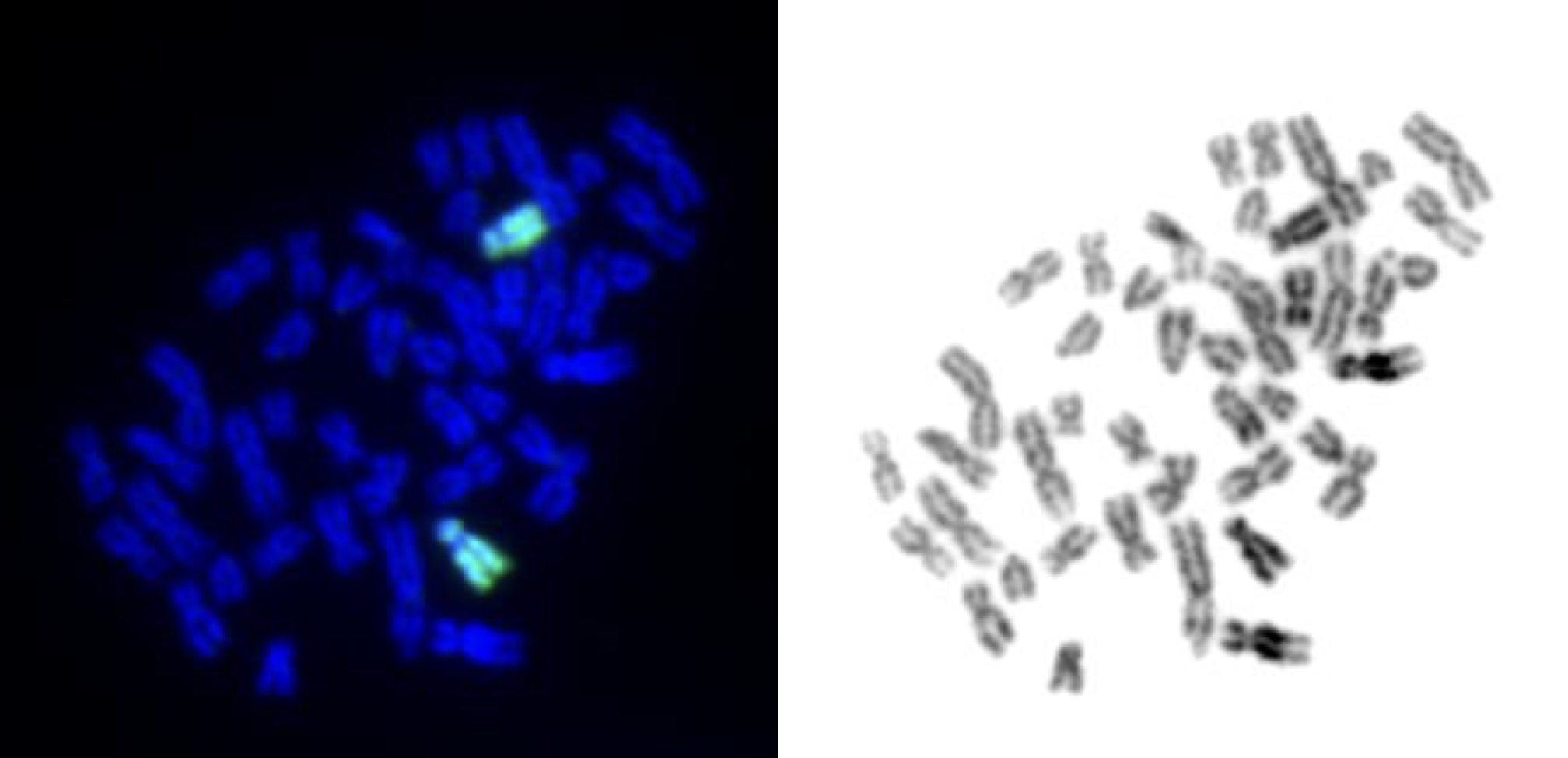

Figure S4I: Gerbil pool P11 hybridized to gerbil Chromosome 12 and Chromosome 17

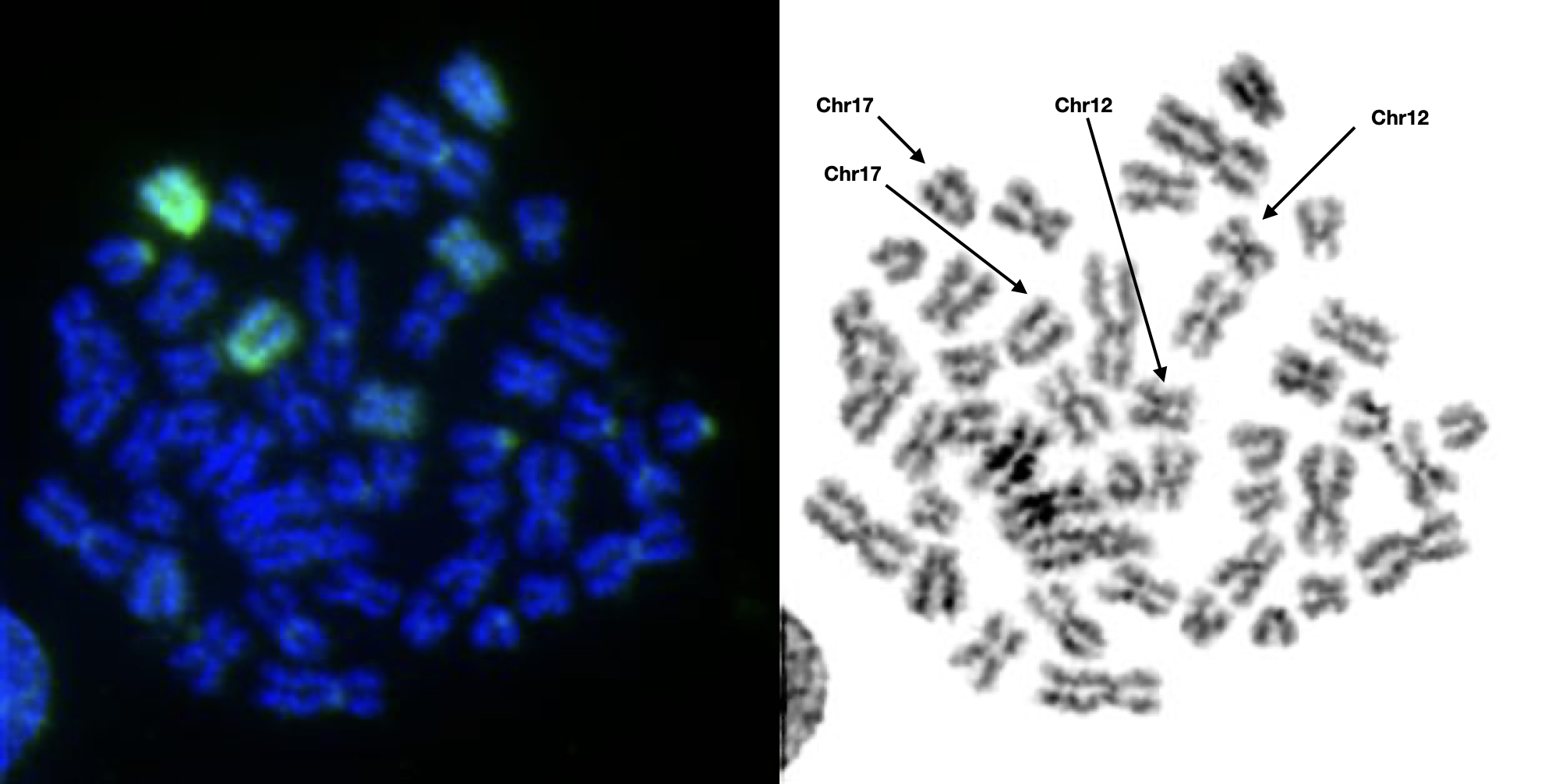

Figure S4J: Gerbil pool P13 hybridized to gerbil Y chromosome.
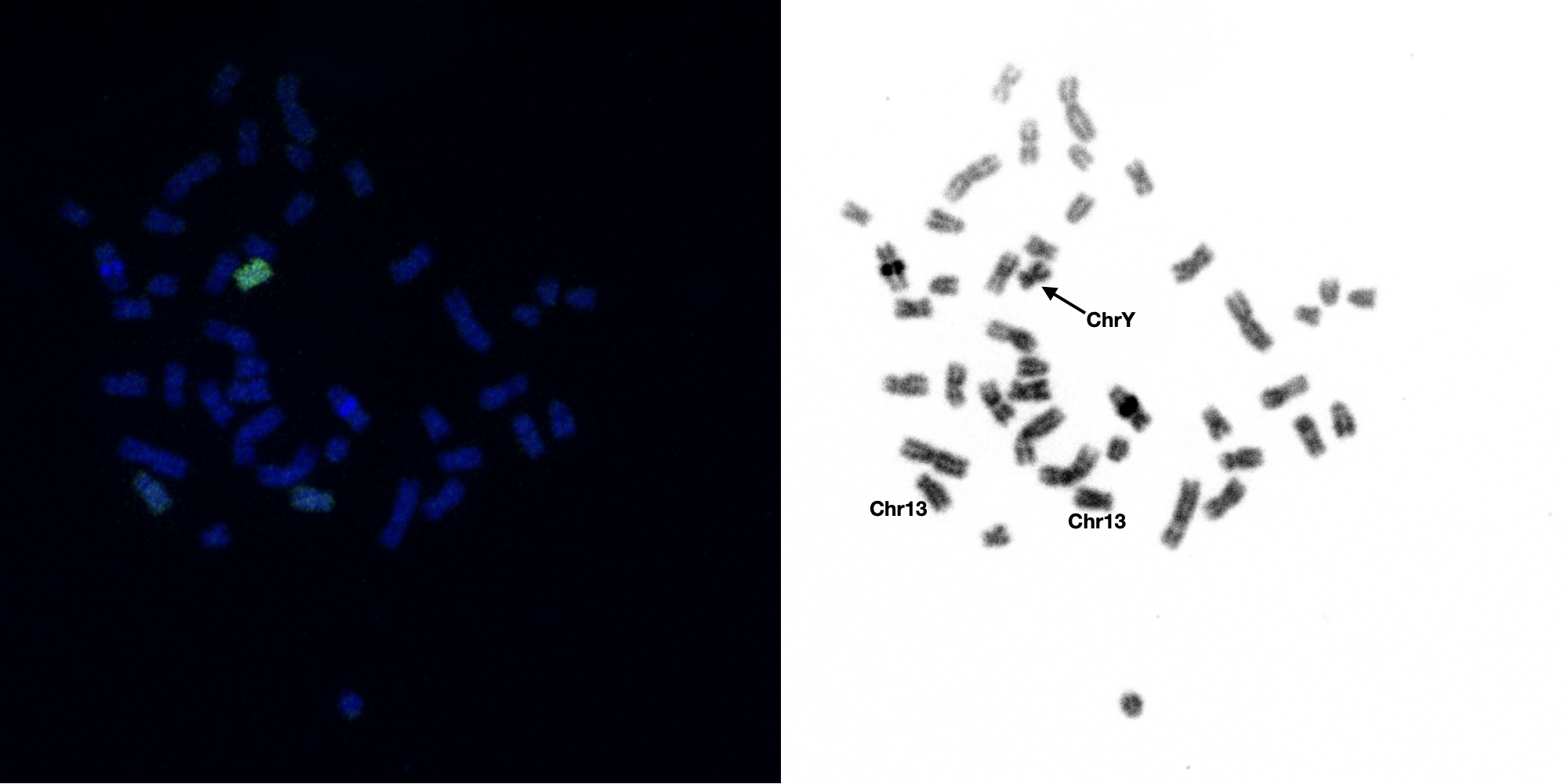

Figure S4K: Gerbil pool P14 hybridized to gerbil chromosome 15 and 18.

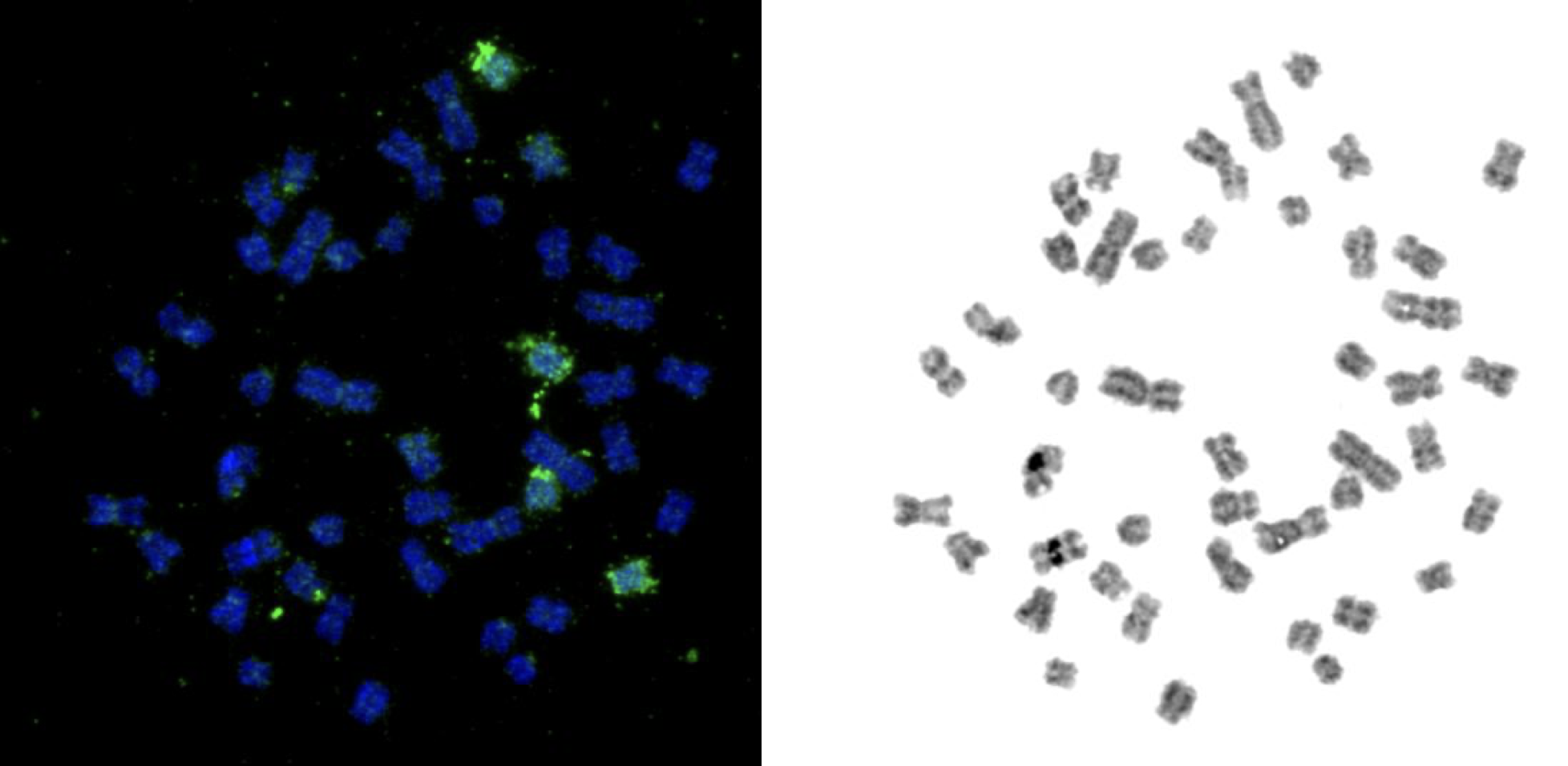

Figure S4L: Gerbil pool P15 hybridized to gerbil chromosome 19 and 20.
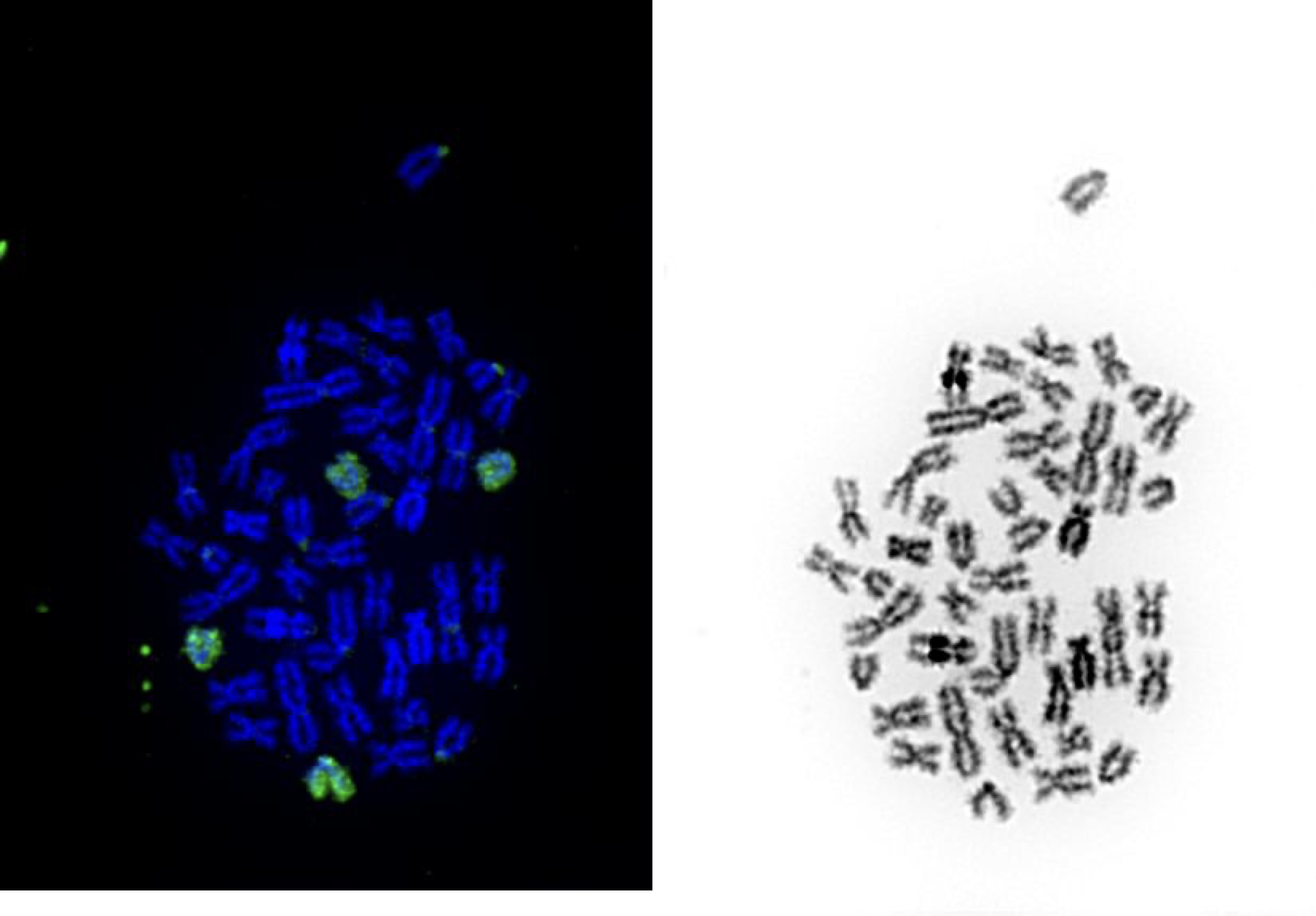

Figure S4M: Gerbil pool P16 (green) hybridized to gerbil chromosome 16. Also gerbil pool P17 (red) hybridized to gerbil chromosome 21.
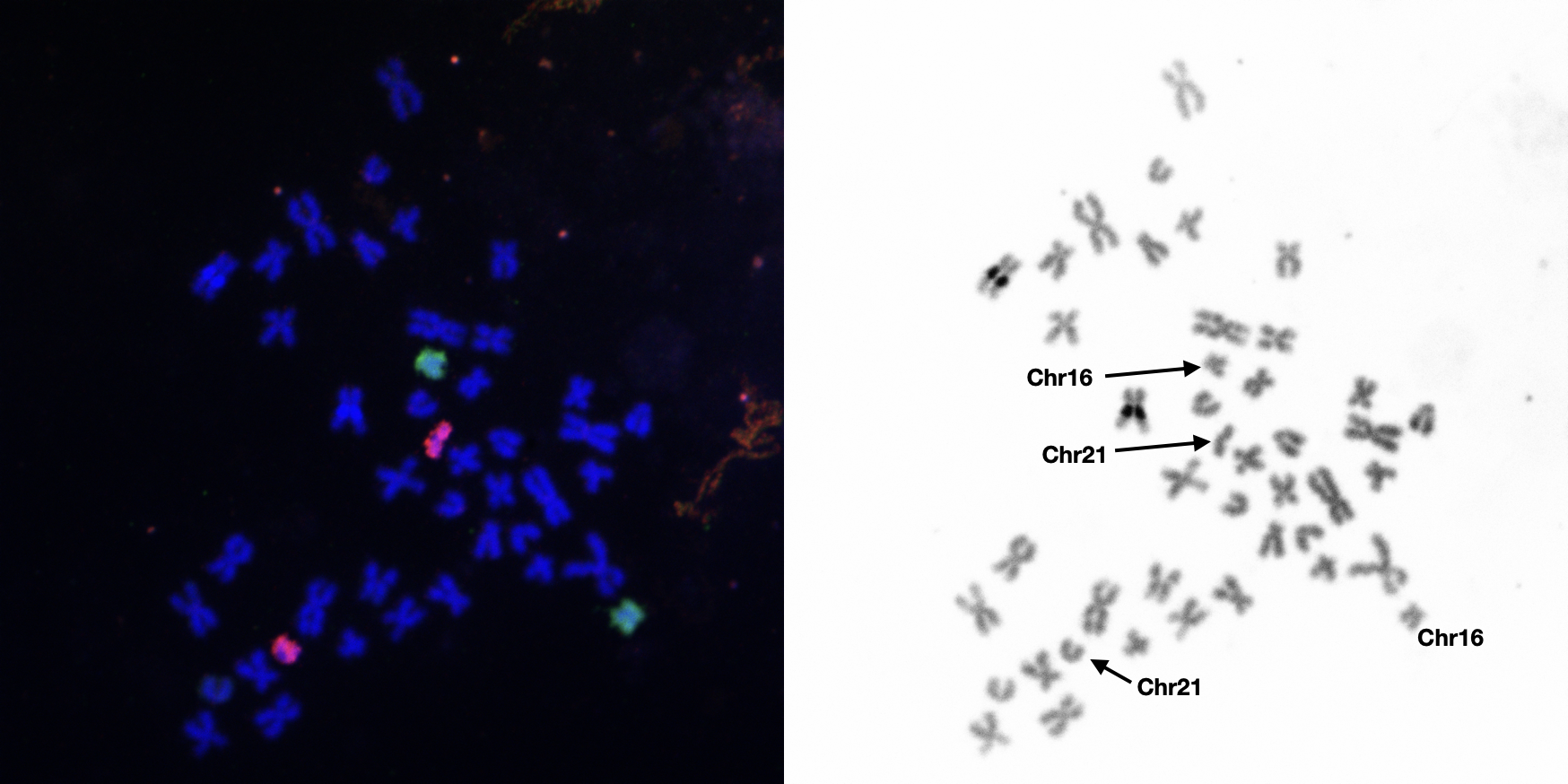

**Distinguishing chromosomes in the same pool**

The final step is to distinguish between scaffolds for the six cases where multiple chromosomes sorted into the same pool. These were pools 3, 4, 7, 11, 14, and 15. This ended up using a complicated chain of logic which is diagrammed in Figure S4 using Pool11 as an example, and the specific inferences for each pool are described below. For each pool we link the *Meriones* physical chromosome with one of the two possible *Meriones* scaffolds using the logic that if A = B, B = C, and C = D then A = D. The full chain of comparisons first links *Meriones* chromosomes to fat sandrat (*Psammomys* *obesus*) chromosomes using comparative DAPI-banding (Figure S5, Link 1; Figure S6), second it links *Psammomys* chromosomes to house mouse (*Mus* *musculus*) chromosomes using a cross-species FISH paint (Figure S5, Link 2, Figure S7). Third, it bioinformatically links *Mus* chromosomes with the unknown *Meriones* scaffolds using shared gene content (Figure S5, Link 3). For this comparison, we used biomart to gather a list of protein-coding genes from mouse chromosomes 2 and 4. From the *Meriones* gff annotation file, we extracted all entries categorised as ‘gene’ for each potential scaffold, extracted the gene name, sorted it and extracted the unique set of gene names to remove duplicates. Then we used the grep case-insensitve and count functions to compare the mouse and *Meriones* gene lists and find the number of shared genes. Once these three links were made, we could infer which *Meriones* scaffold went with which *Meriones* chromosome by stepping back through the chain. We used the sandrat intermediate as the data linking sandrat to mouse had previously been generated and is in preparation by our collaborators in another manuscript. Here, we present the smallest slice of these data necessary to make the logical links. These data are summarised in Table S3.

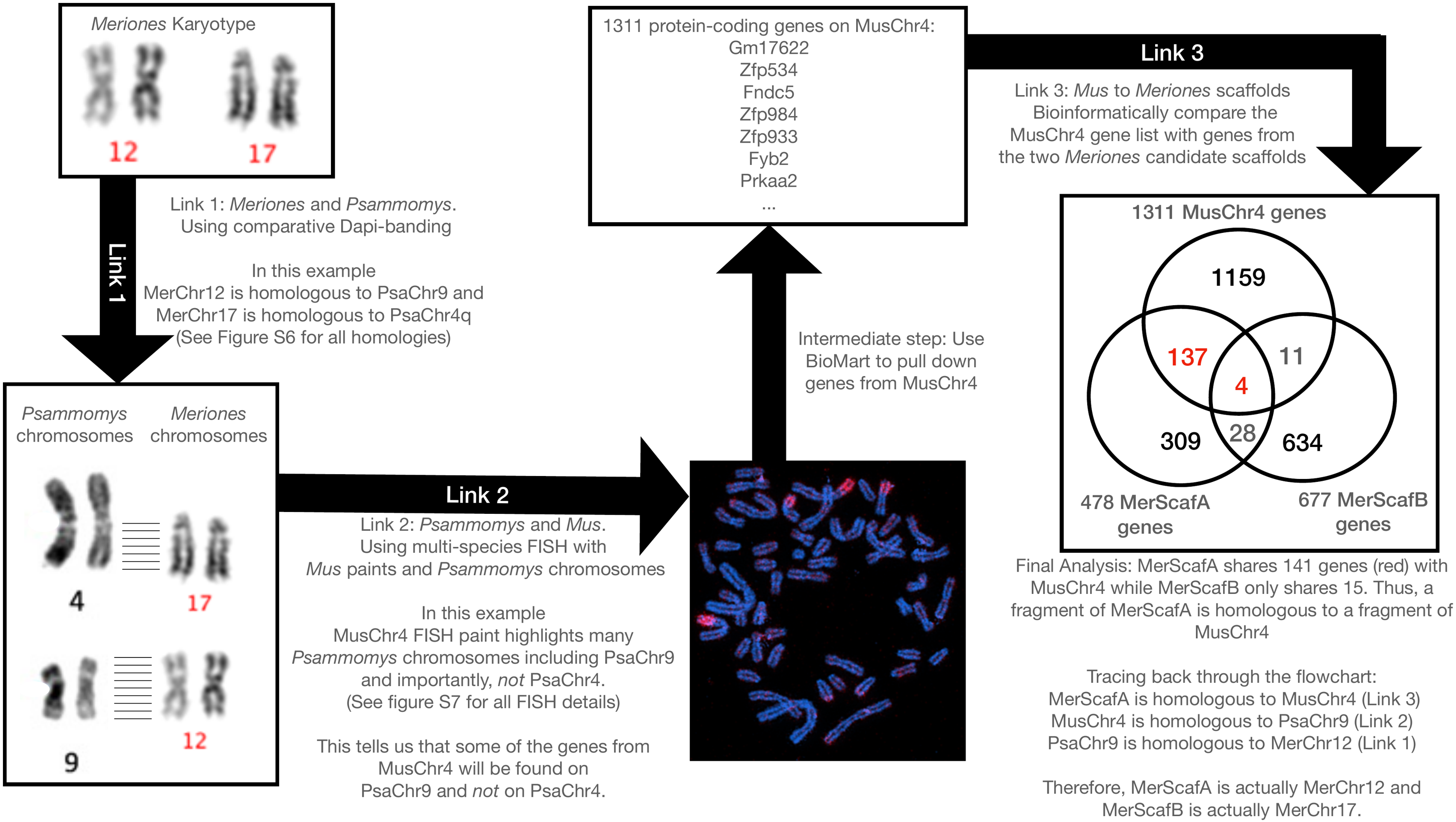

Figure S5: The logic of the chromosome assignments for the pools containing two chromosomes using Pool 11 as an example. Pool 11 contains *Meriones* chromosomes 12 and 17 and *Meriones* scaffolds A and B. The goal is to determine which chromosome goes with which scaffold.

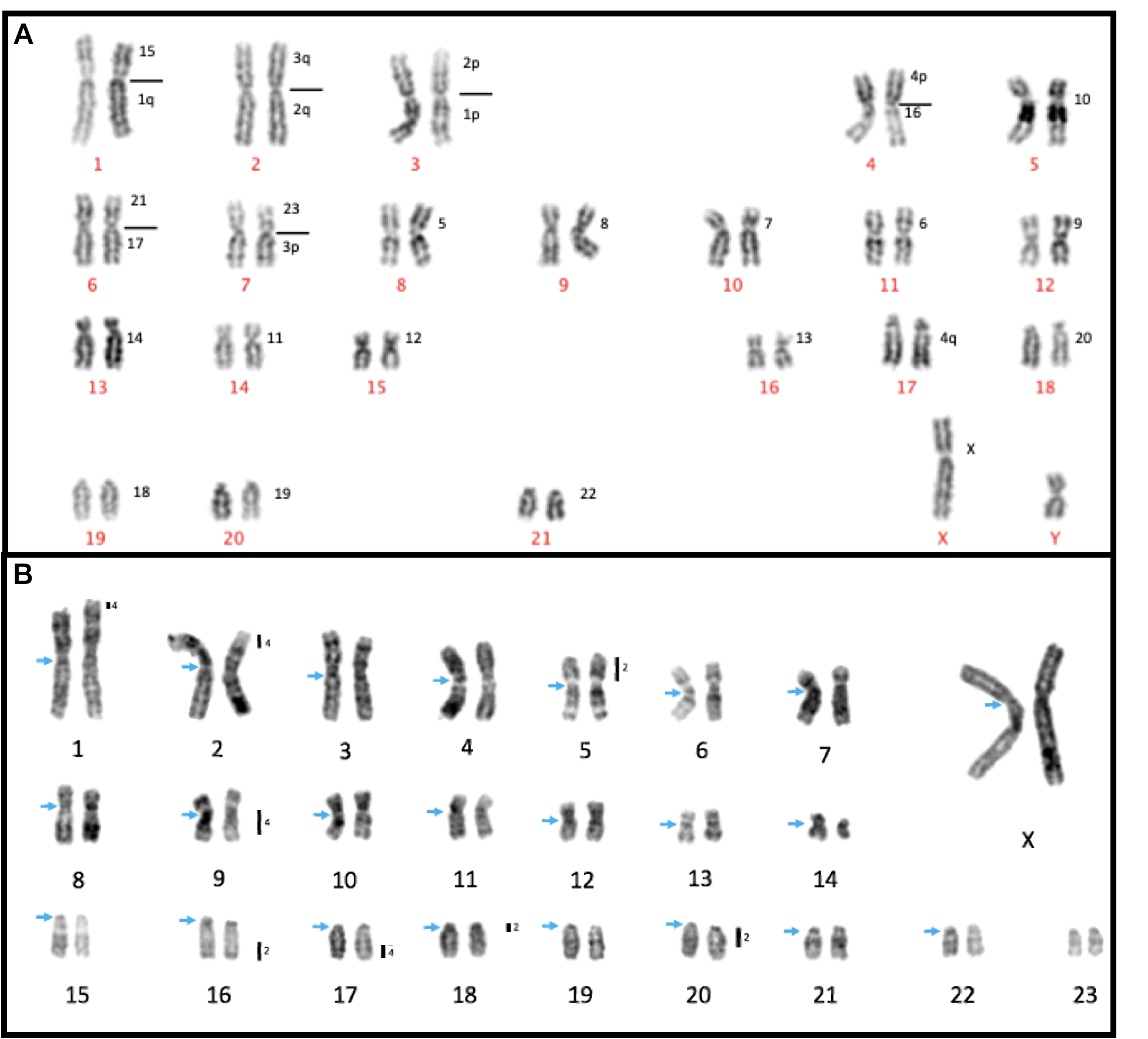

Figure S6: Comparative DAPI-banding between Sandrat and Gerbil. A) a Dapi-banded *Meriones* karyotype with chromosomes labelled in red and Sandrat homologies labelled alongside in black. B) A DAPI-banded *Psammomys* karyotype with chromosomes labelled in black and centromeres marked in blue. Mouse chromosome 2 and 4 paint locations are noted alongside Sandrat chromosomes as black bars.

Figure S7A: *Mus* Chromosome 2 paints *Psammomys* Chromosomes 5, 16, 18, and 20.

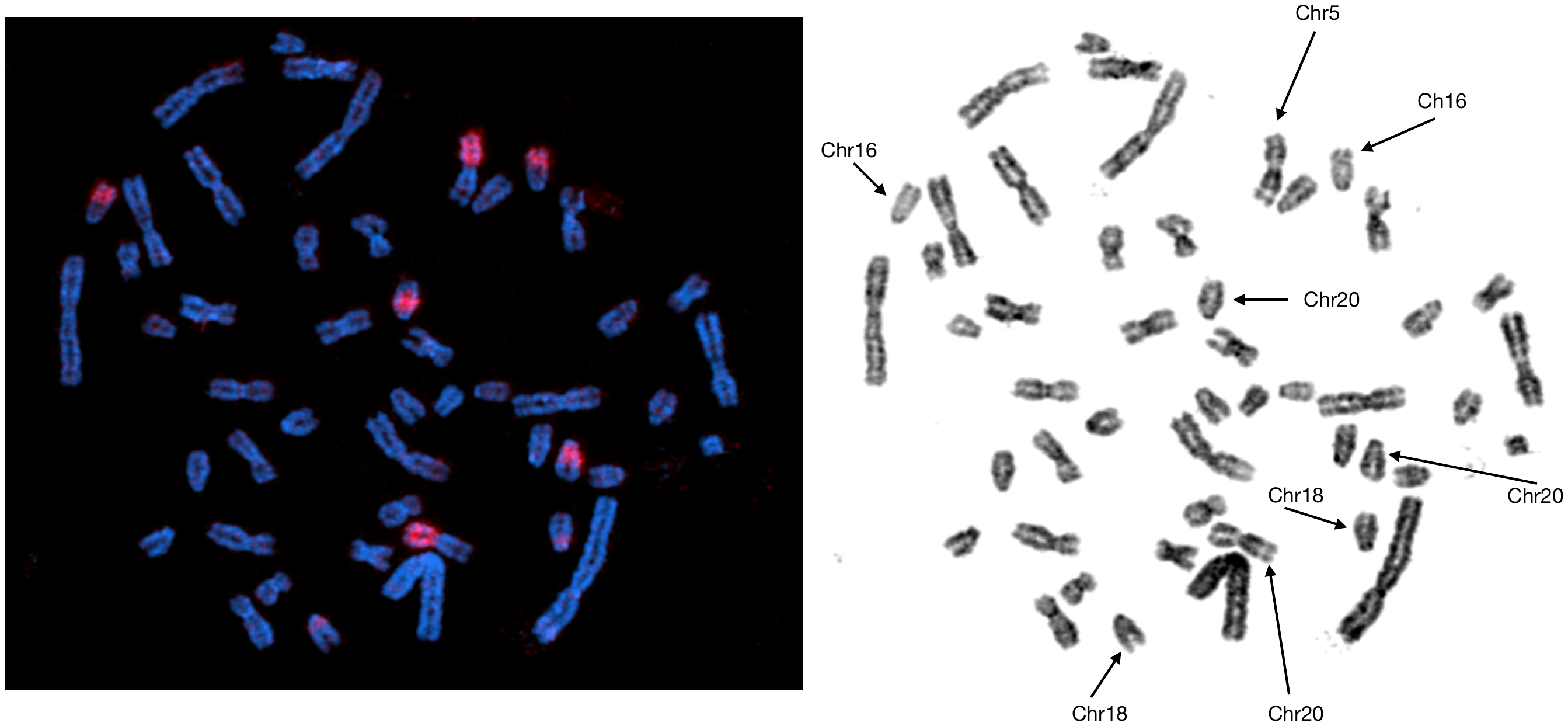

Figure S7B: *Mus* Chromosome 4 paints *Psammomys* Chromosomes 1, 2, 9 and 17.

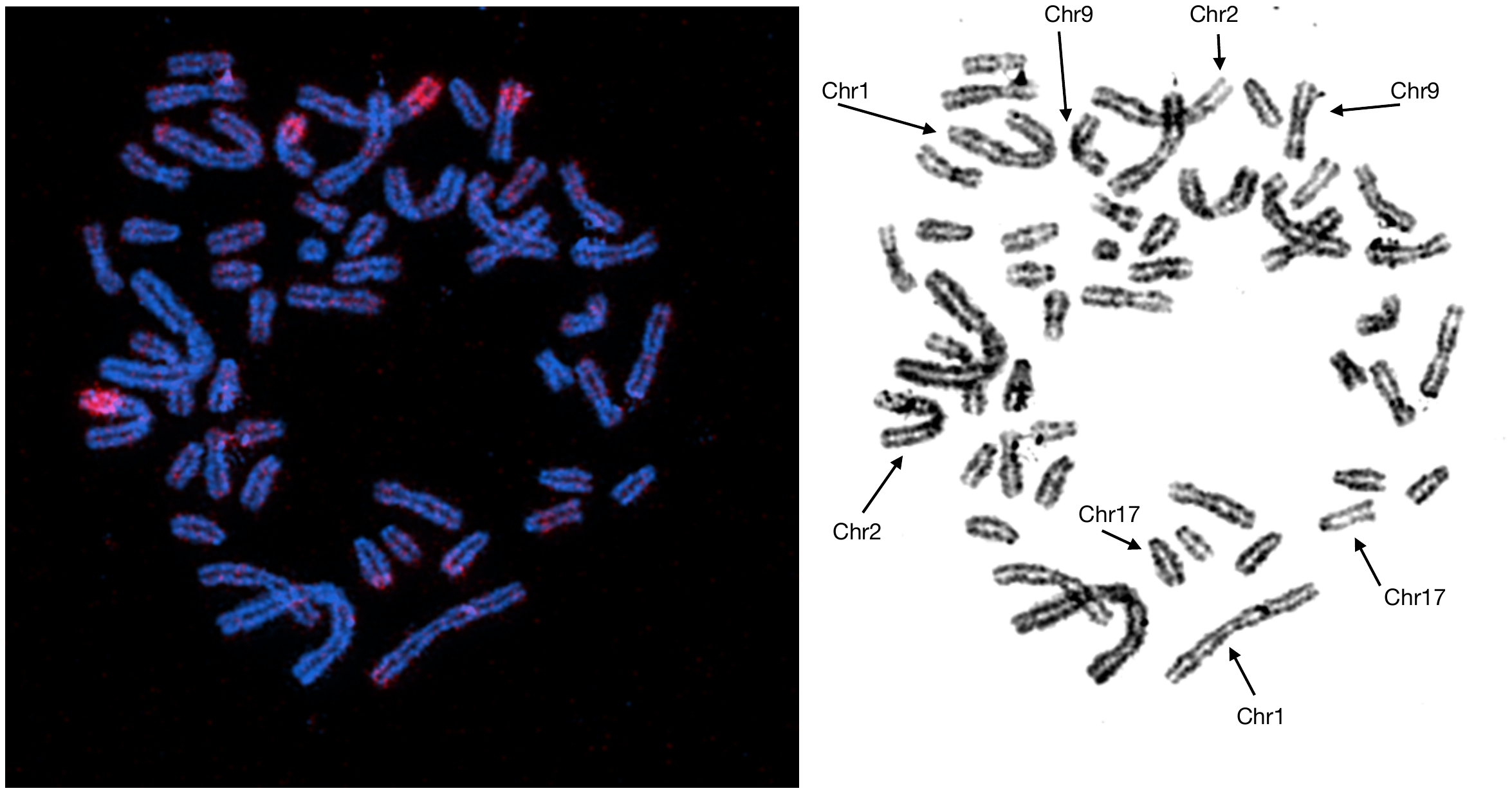

Table S3: Summary of Gerbil-Sandrat-Mouse homologies for specific pools.

| **Gerbil Pool** | **Gerbil Chromosome** | **Sandrat Chromosomes** | **Mouse Chromosomes 2 or 4?** |
| --- | --- | --- | --- |
| P3 | 1 | 1q, 15 | - |
|  | 3 | 1p, 2p | 4 |
| P4 | 5 | 10 | - |
|  | 6 | 17, 21 | 4 |
| P7 | 8 | 5 | 2 |
|  | 10 | 7 | - |
| P11 | 12 | 9 | 4 |
|  | 17 | 4q | 2 |
| P14 | 15 | 12 | - |
|  | 18 | 20 | 2 |
| P15 | 19 | 18 | 2 |
|  | 20 | 19 | - |

**Pool3**

Pool3 contains MerChr1 and MerChr3.

Link 1: MerChr3 is homologous to PsaChr1p and 2p (Figure S6A).

Link 2: PsaChr1q and PsaChr2p both have homology with MusChr4 (Figure 7B).

Link 3: MusChr4 shares 91 genes with one potential *Meriones* scaffold and 612 with the other.

We assigned ‘MerChr3’ to the scaffold that shared more genes with MusChr4.

**Pool4**

Pool4 contains MerChr5 and MerChr6.

Link 1: MerChr6 is homologous to PsaChr17 and 21 (Figure S6A).

Link 2: PsaChr17 has homology with MusChr4 (Figure S7B).

Link 3: MusChr4 shares 20 genes with one potential *Meriones* scaffold and 83 with the other.

We assigned ‘MerChr6’ to the scaffold that shared more genes with MusChr4.

An additional line of evidence for this pools is that one of the scaffolds has a run of repeats 35Mb long in the middle of the scaffold. This repetive expansion happens to be the same location as the dark stain of Meriones Chr5 suggesting they are the same. This line of evidence agrees with the logic above.

**Pool7**

Pool7 contains MerChr8 and MerChr10.

Link 1: MerChr8 is homologous to PsaChr5 (Figure S6A).

Link 2: PsaChr5 has homology with MusChr2 (Figure S7B).

Link 3: MusChr2 shares 644 genes with one potential *Meriones* scaffold and 333 with the other.

We assigned ‘MerChr8’ to the scaffold that shared more genes with MusChr2.

**Pool11**

Pool11 contains MerChr12 and MerChr17.

Link 1: MerChr12 is homologous to PsaChr9 (Figure S5, Link 1; Figure 6A).

Link 2: PsaChr9 has homology with MusChr4 (Figure S5, Link 2; Figure 7B).

Link 3: MusChr4 shares 141 genes with one potential *Meriones* scaffold and 15 with the other (Figure S5, Link 3).

We assigned ‘MerChr12’ to the scaffold that shared more genes with MusChr4.

**Pool14**

Pool14 contains MerChr15 and MerChr18.

Link 1: MerChr18 is homologous to PsaChr20 (Figure S6A).

Link 2: PsaChr20 has homology with MusChr2 (Figure S7B).

Link 3: MusChr2 shares 209 genes with one potential *Meriones* scaffold and 366 with the other.

We assigned ‘MerChr18’ to the scaffold that shared more genes with MusChr2.

**Pool15**

Pool15 contains MerChr19 and MerChr20.

Link 1: MerChr19 is homologous to PsaChr18 (Figure S6A).

Link 2: PsaChr18 has homology with MusChr2 (Figure S7B).

Link 3: MusChr2 shares 184 genes with one potential *Meriones* scaffold and 124 with the other.

We assigned ‘MerChr19’ to the scaffold that shared more genes with MusChr2.

Table S4: Comparison of three published *Meriones* genomes.

| Citation | Primary Sequencing | Scaffolding | Bases | Number of N’s | Contigs | Scaffolds | Any chromosome-length scaffolds? | Scaffolds assigned to chromosome? | BUSCO completeness |
| --- | --- | --- | --- | --- | --- | --- | --- | --- | --- |
| Zorio  et al (2019) | Illumina | Mate-pair | 2,620,810,971 | 120,553,581 | 384,902 | 384,902 | No | No | 93.6% (n=6,192) |
| Cheng  et al (2019) | Illumina | Dovetail Hi-C | 2,543,403,711 | 30,559,892 | 413,334 | 297,728 | Yes | No | 86% (n=3,023) |
| The genome presented here | PacBio HiFi | Dovetail Omni-C, Oxford NanoPore Ultra-long,  a genetic map,  and BioNano optical mapping | 2,702,217,419 | 5,301 | 245 | 194 | Yes | Yes | 92.3% (n=13,798) |

Citations: (Cheng et al. 2019; Zorio et al. 2019)

Figure S8: GC content, Gene density, Entropy, and Linguistic complexity for all chromosomes. GC content is calculated in 1kb sliding windows with a step size of 1kbp. Gene density is calculated in 1Mbp sliding windows with a step size of 1kb. Entropy and Linguistic complexity are both calculated with 10kbp sliding windows with a step size of 1kbp using NeSSie (Berselli et al. 2018). For chromosomes with multiple scaffolds, different scaffolds are shown in alternating white and grey bands.

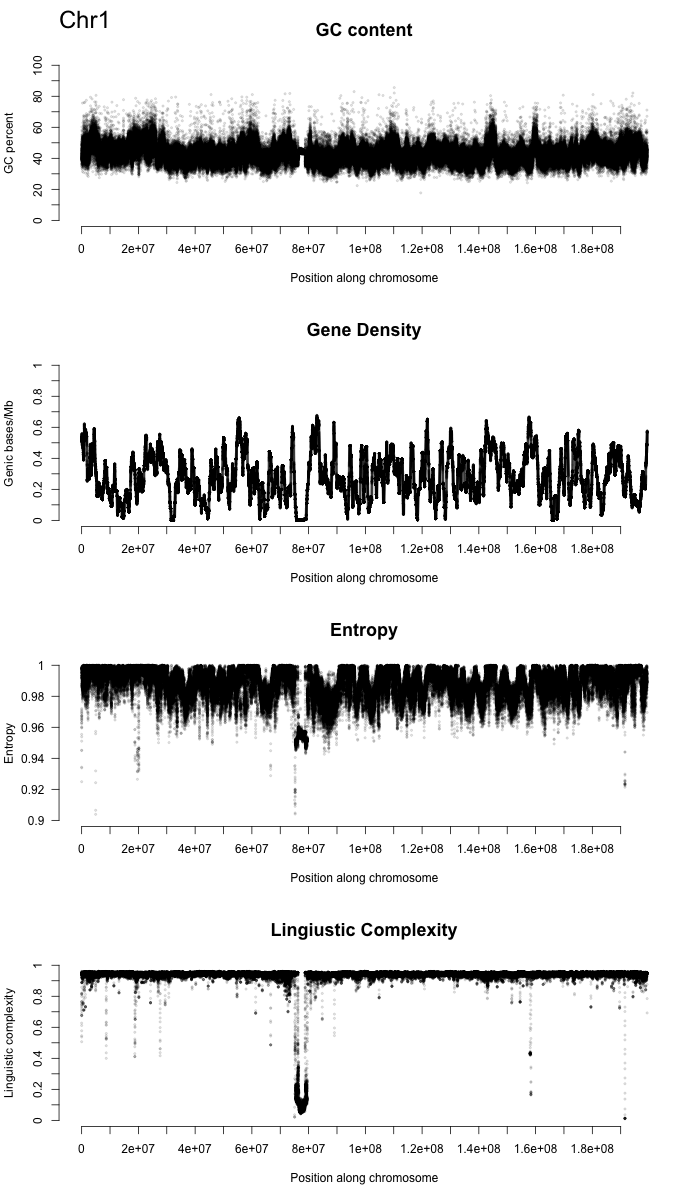

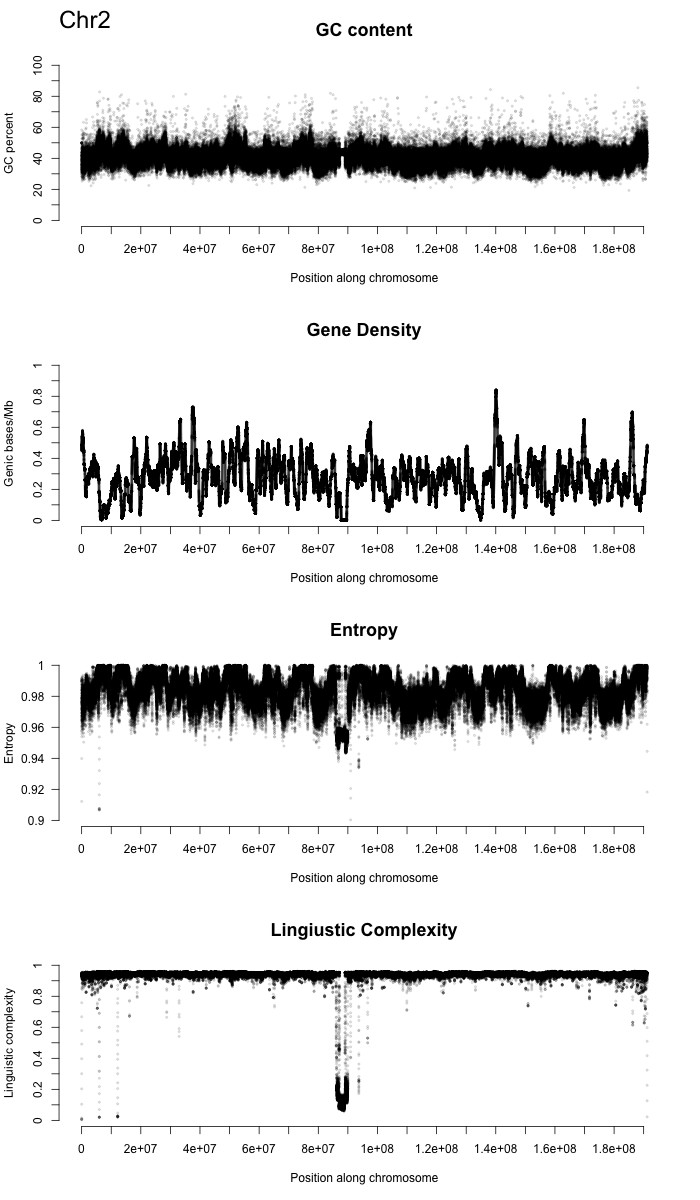

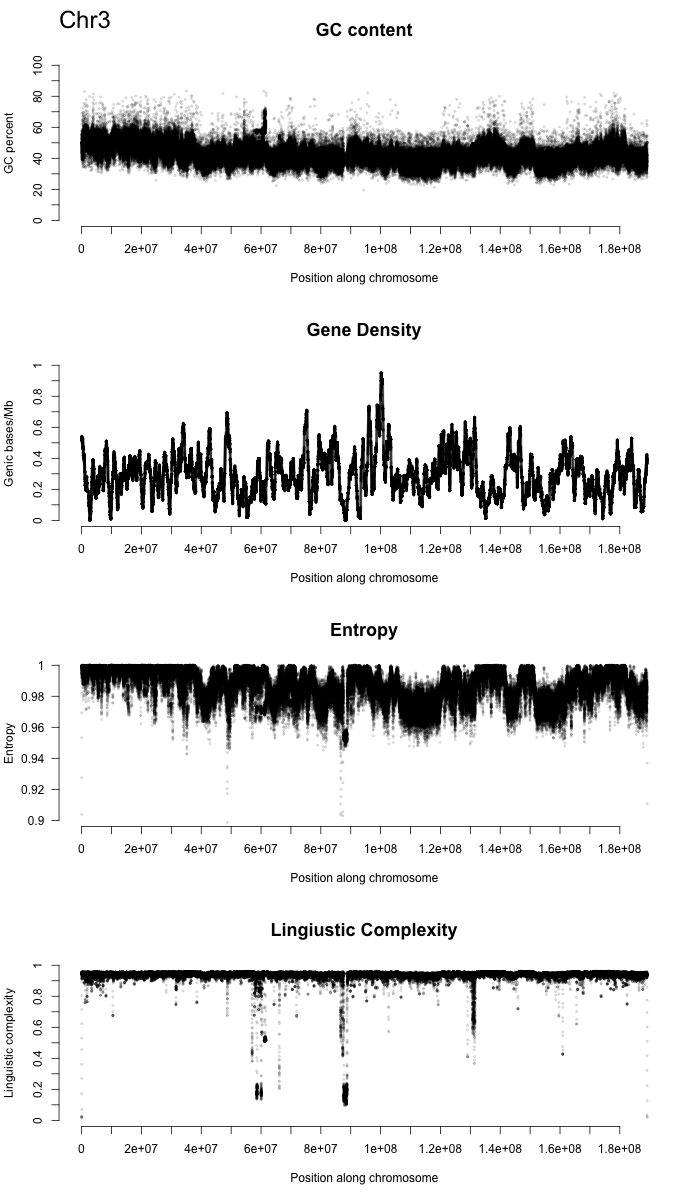

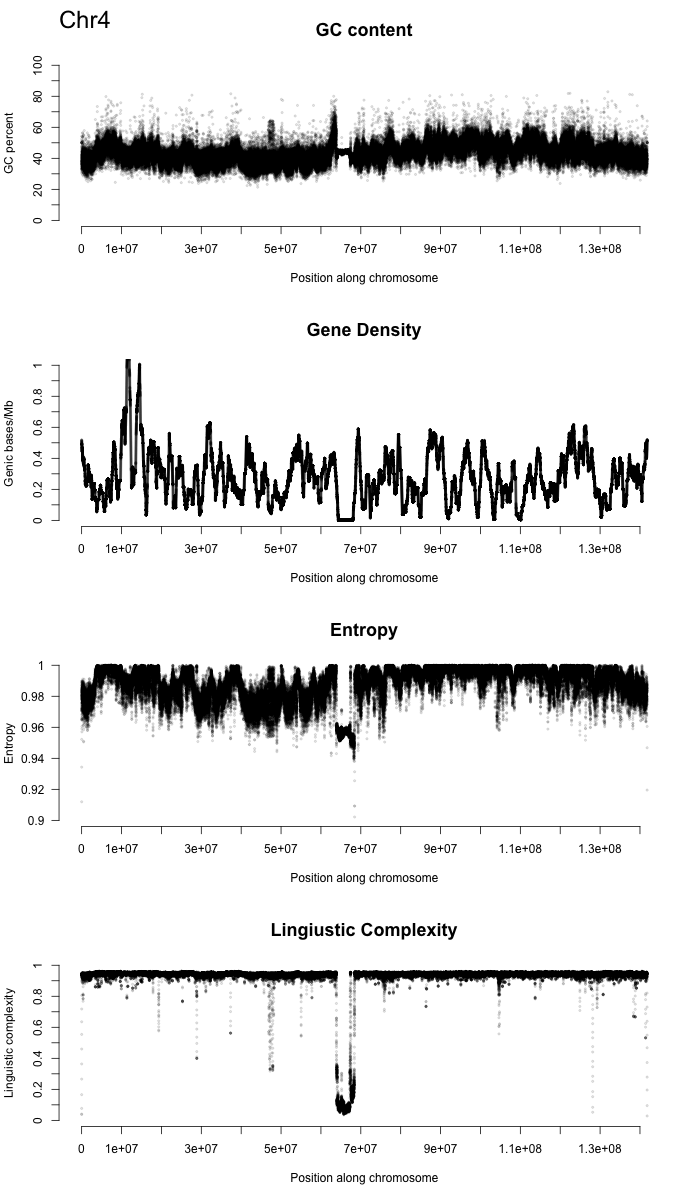

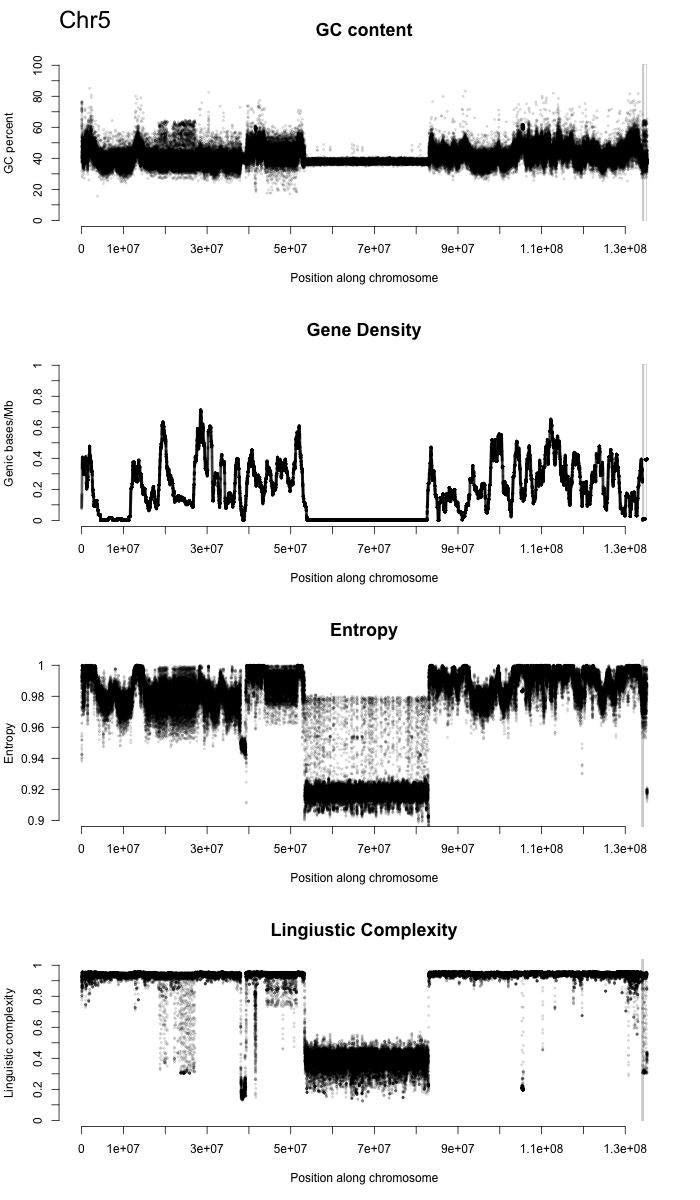

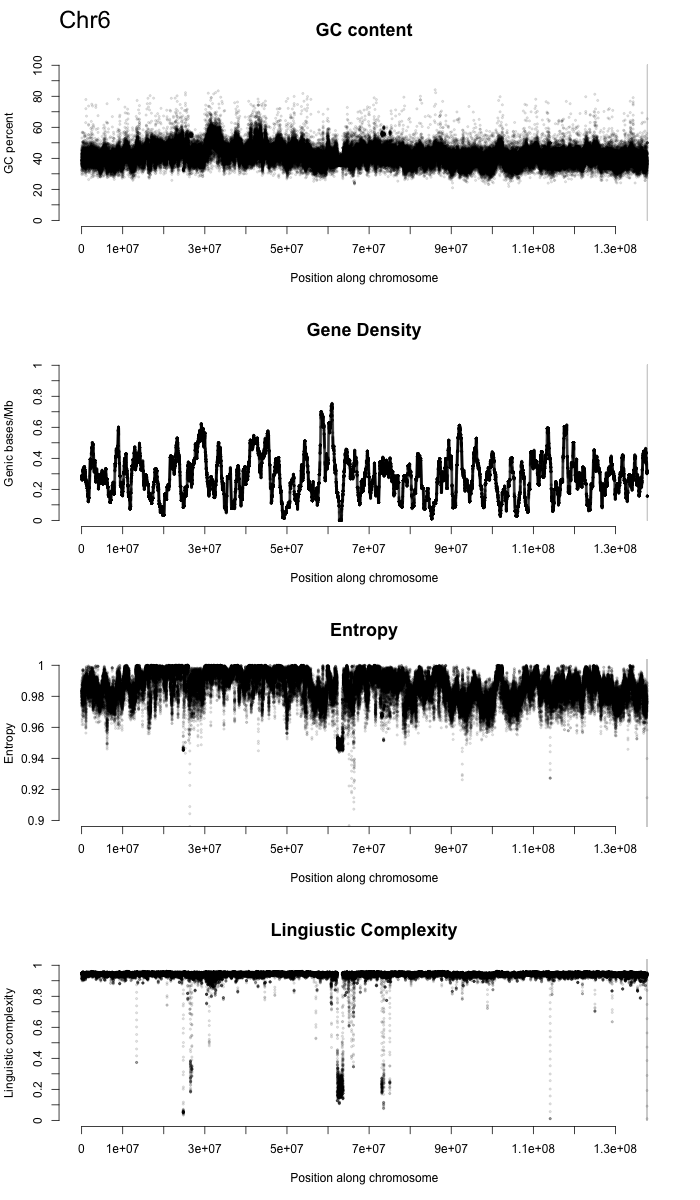

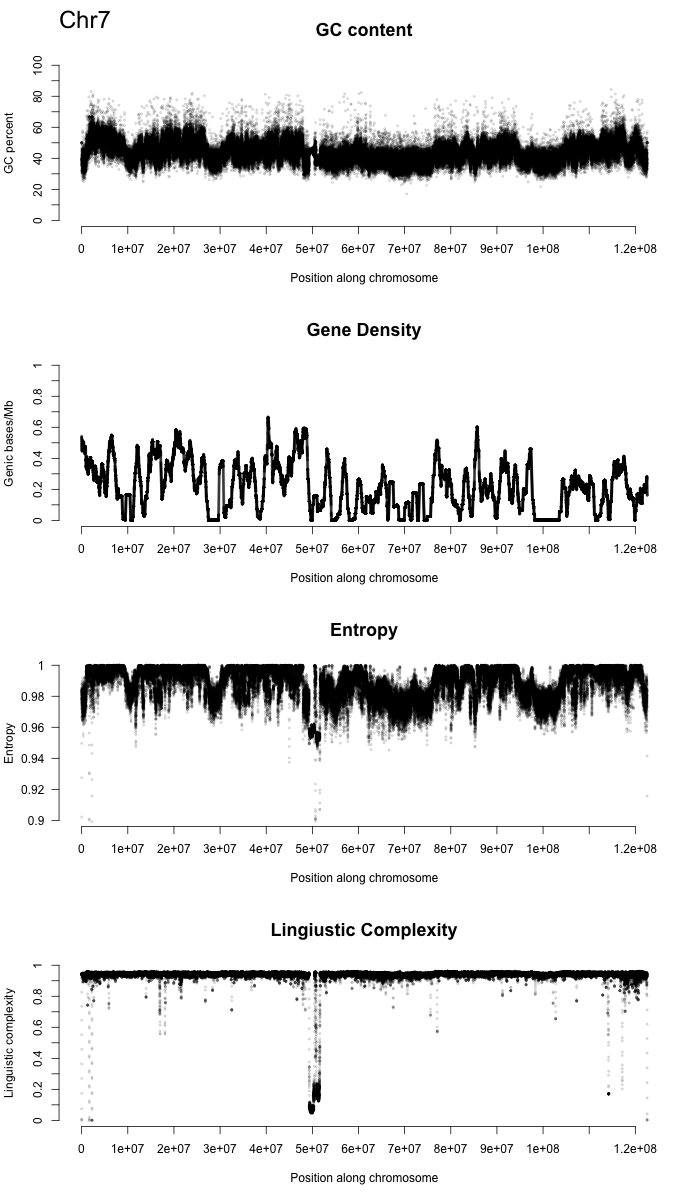

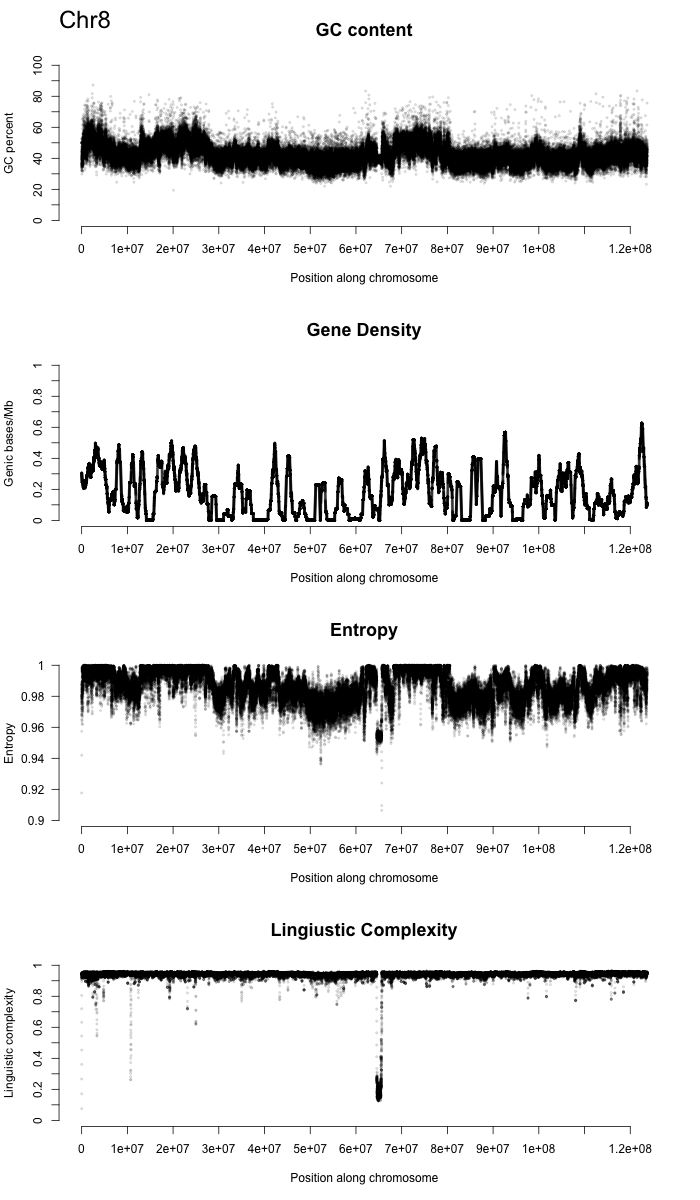

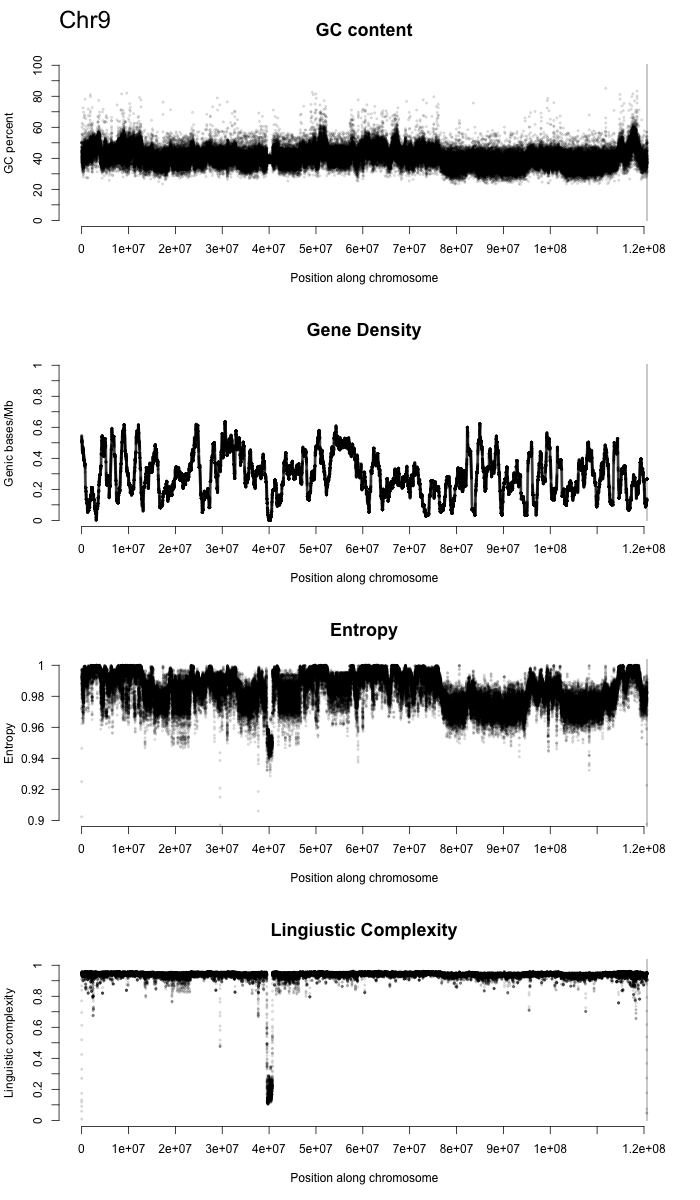

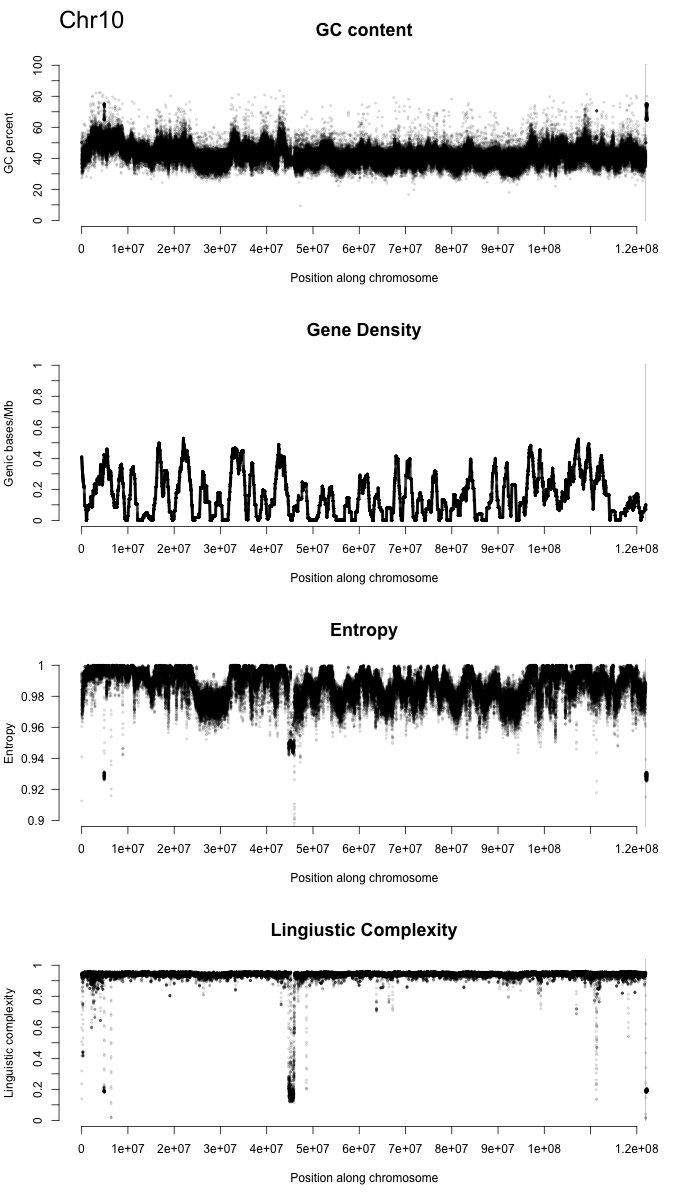

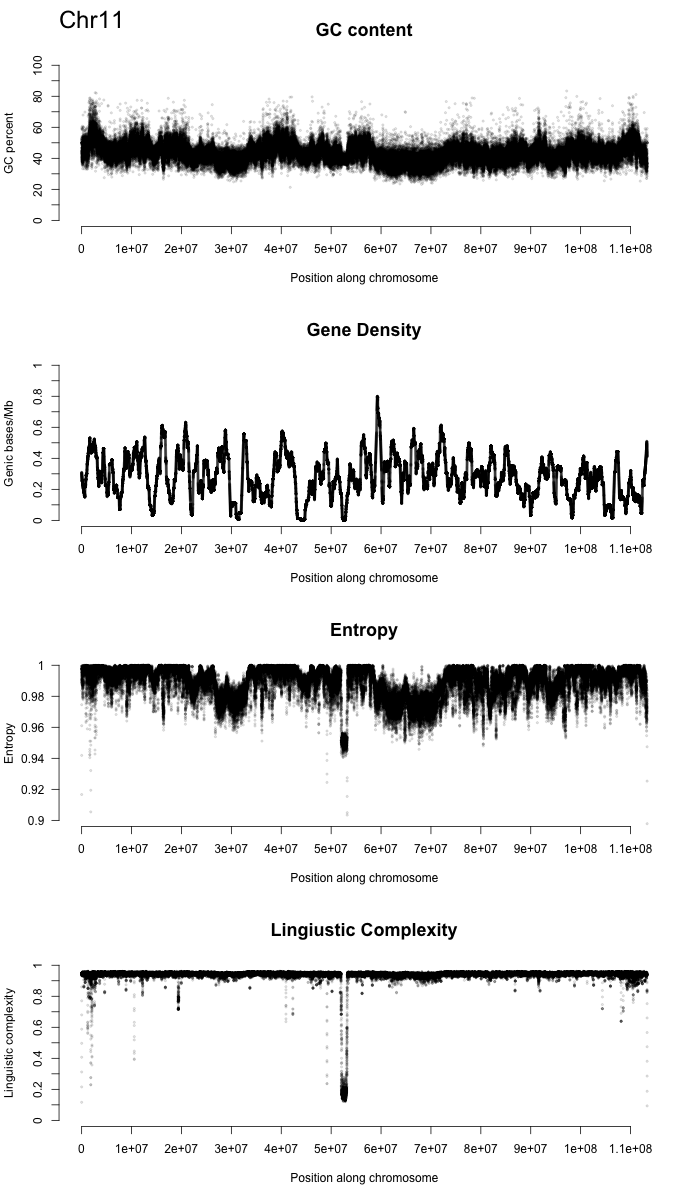

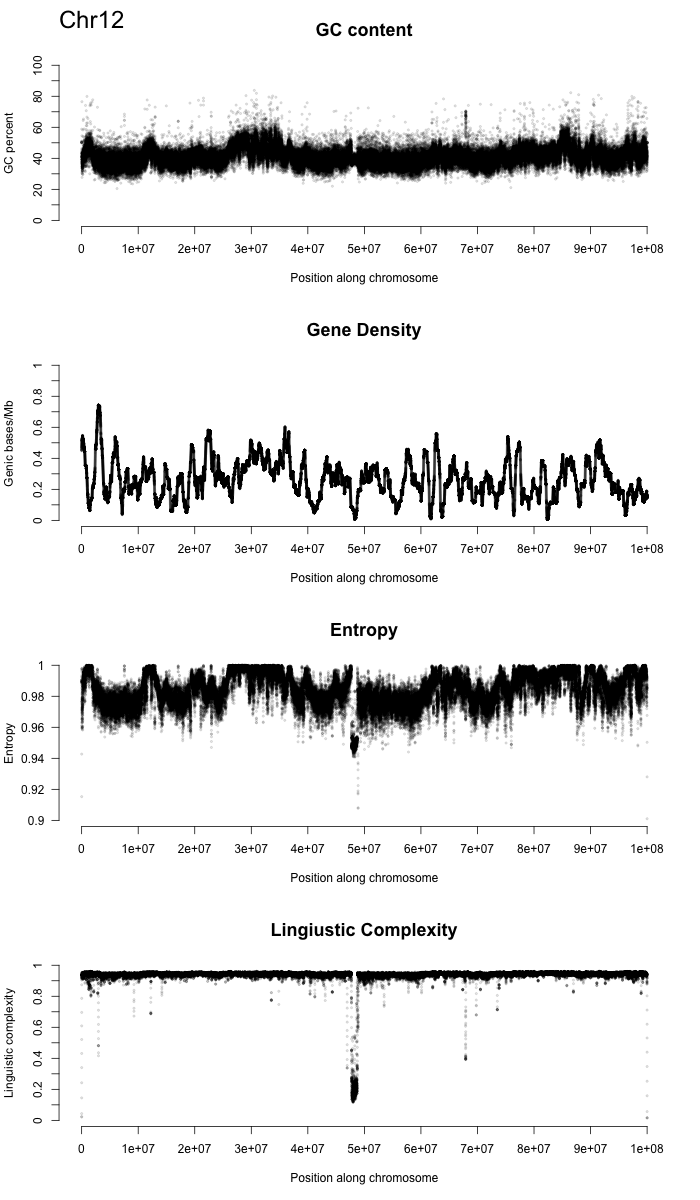

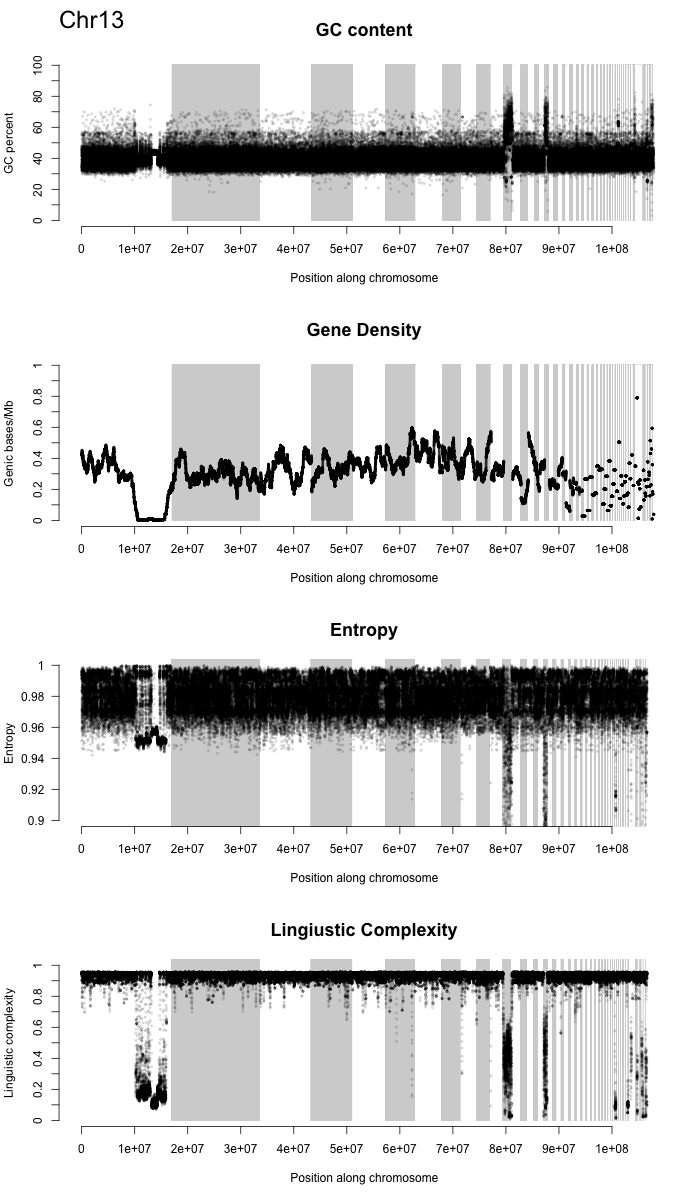

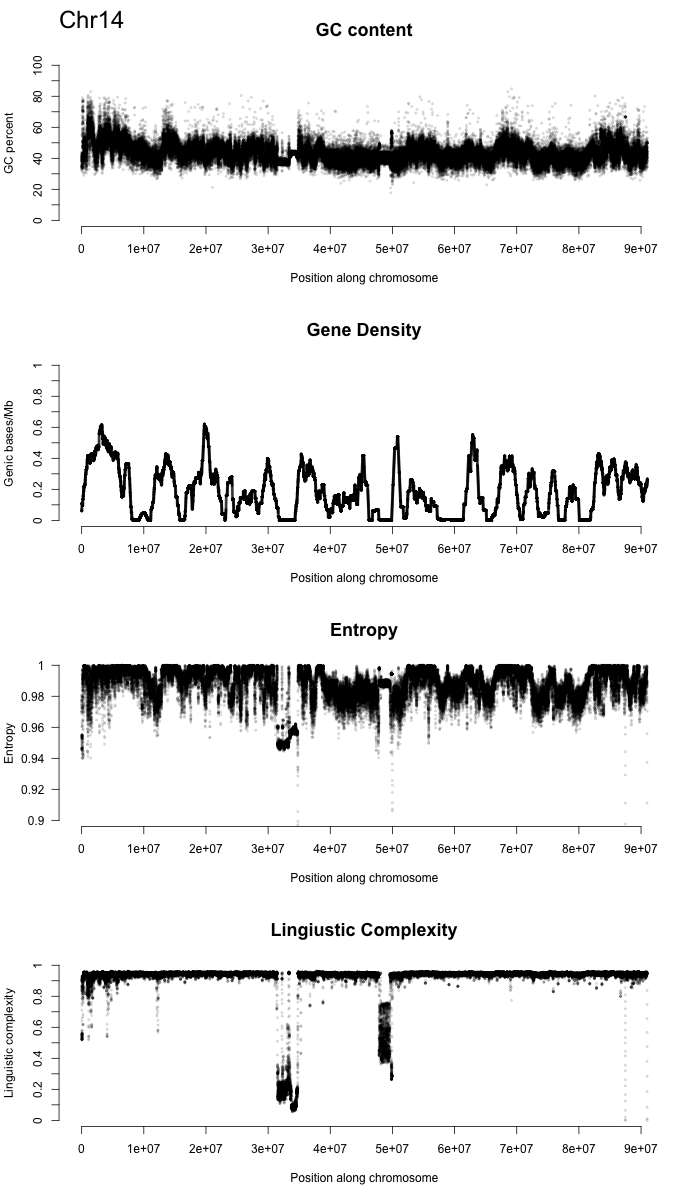

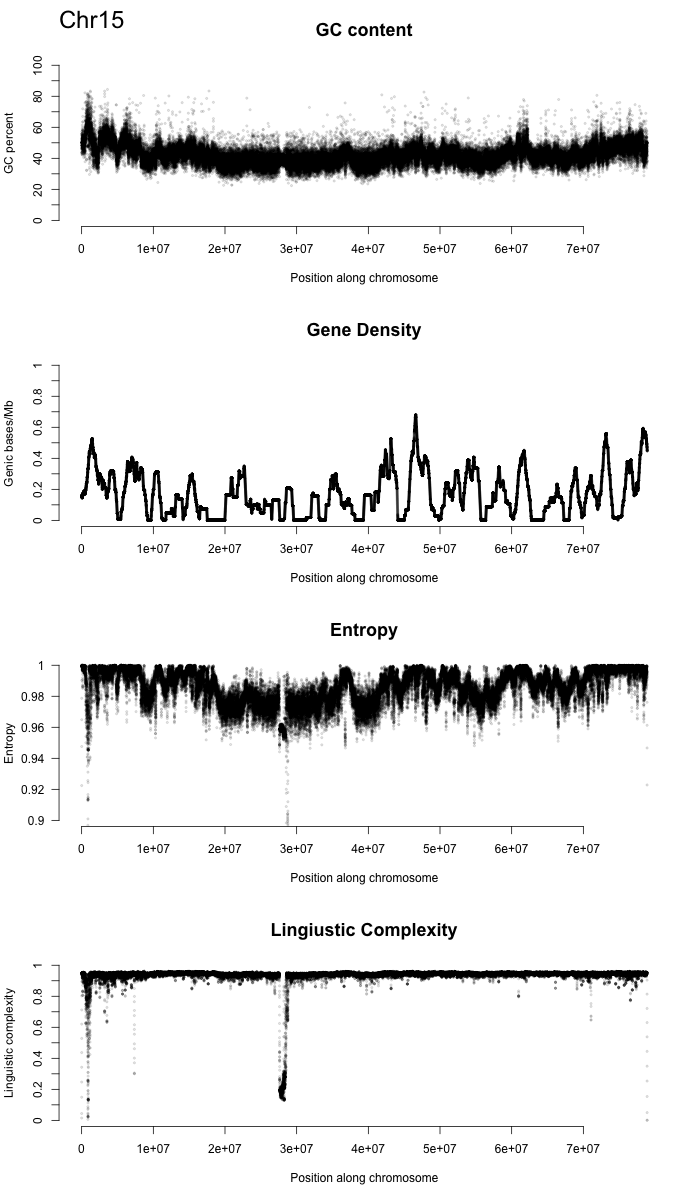

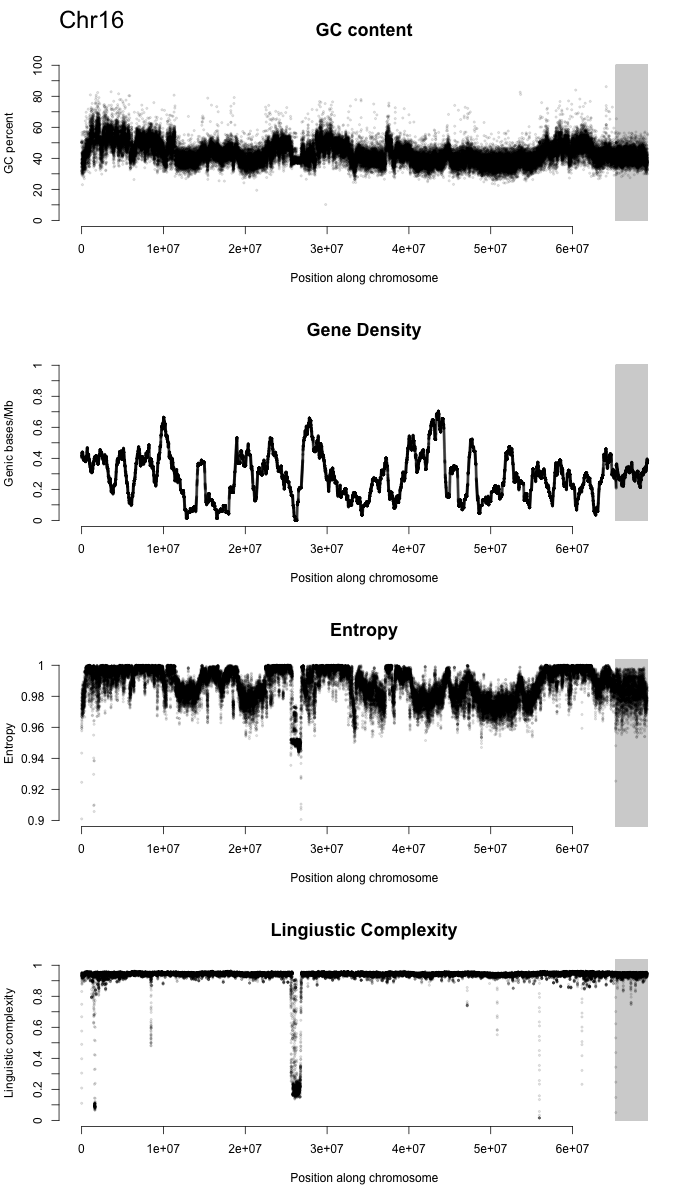

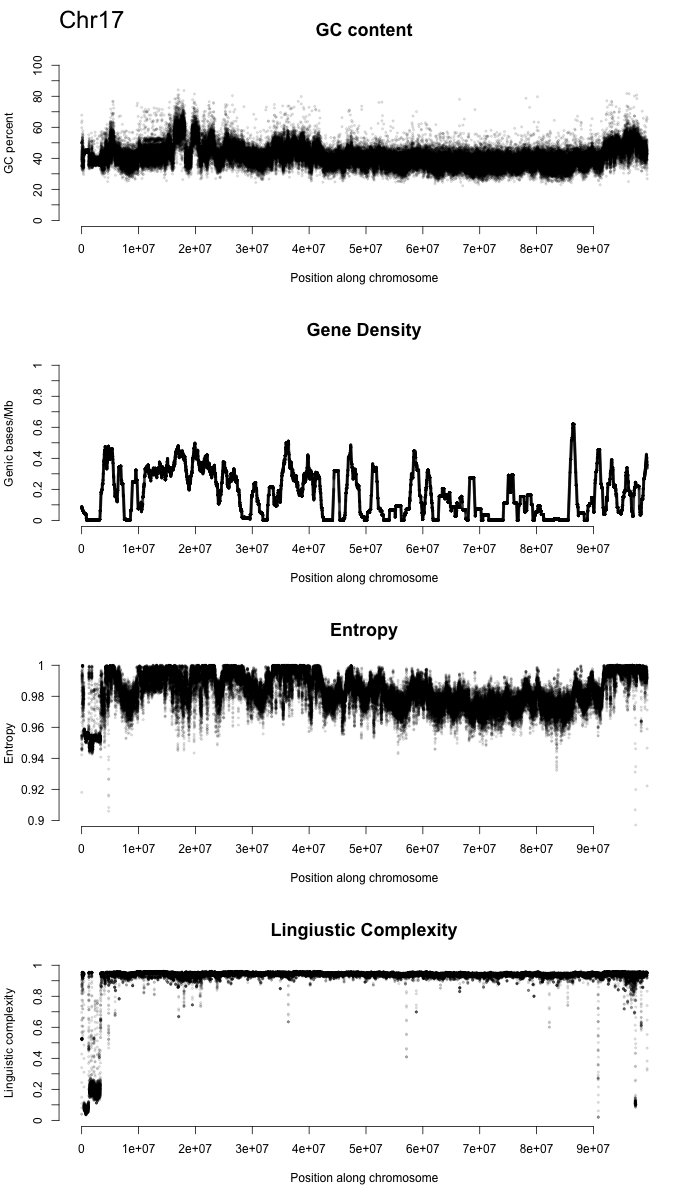

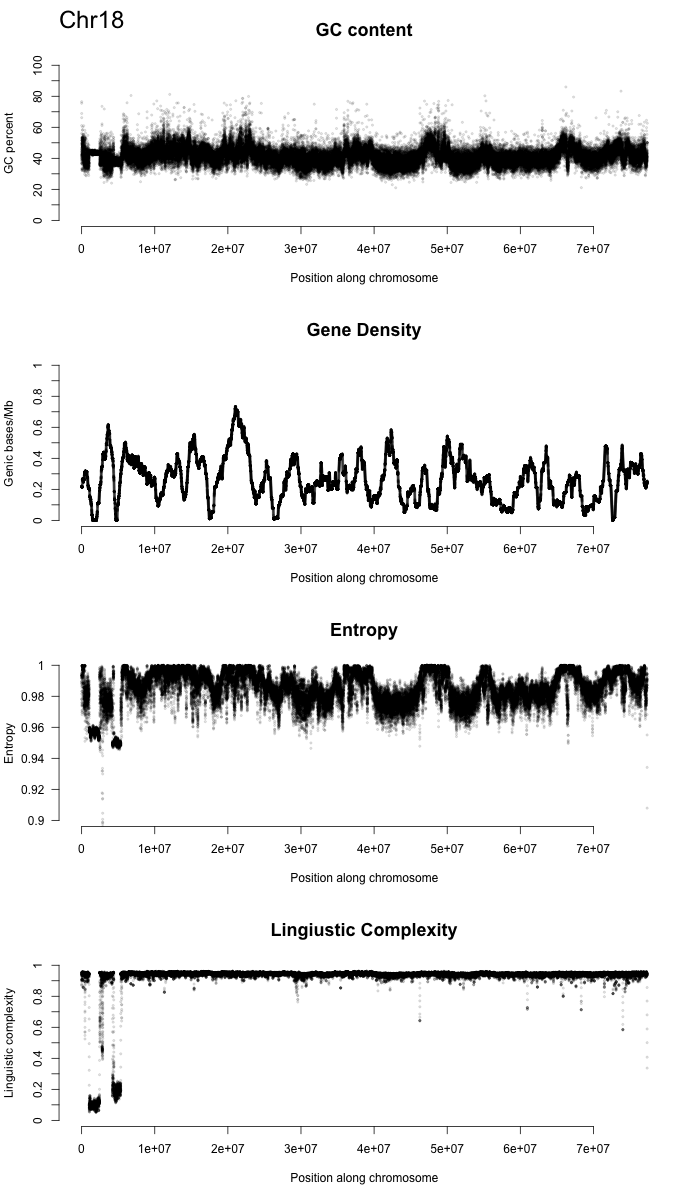

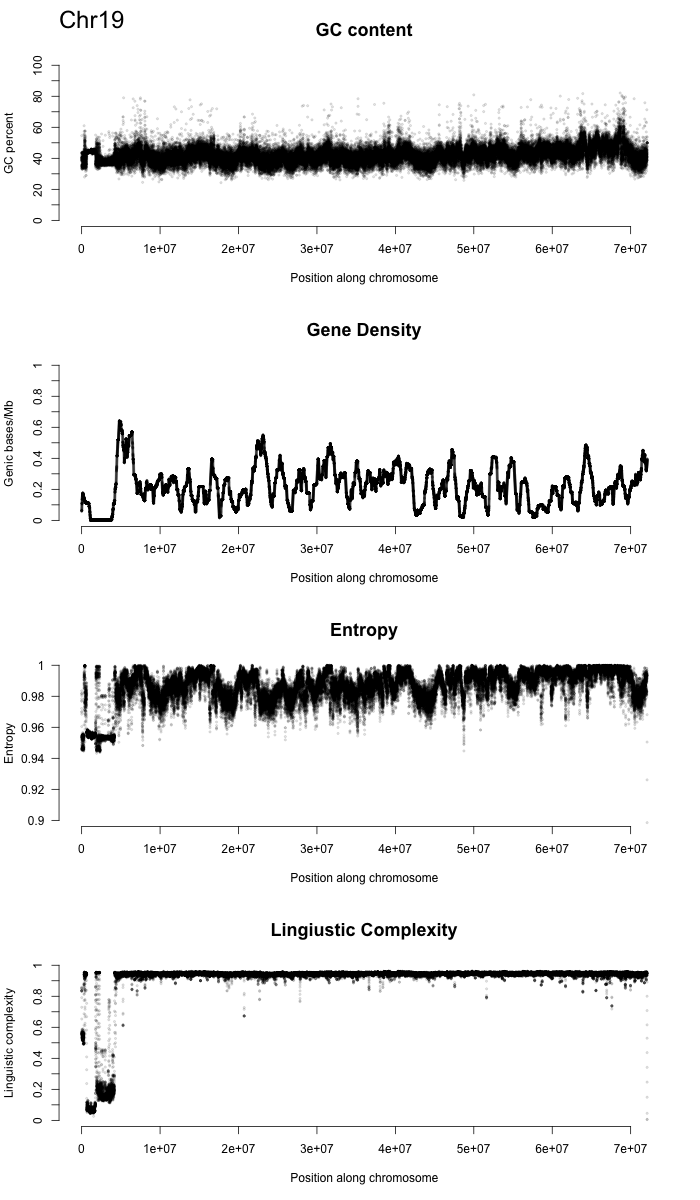

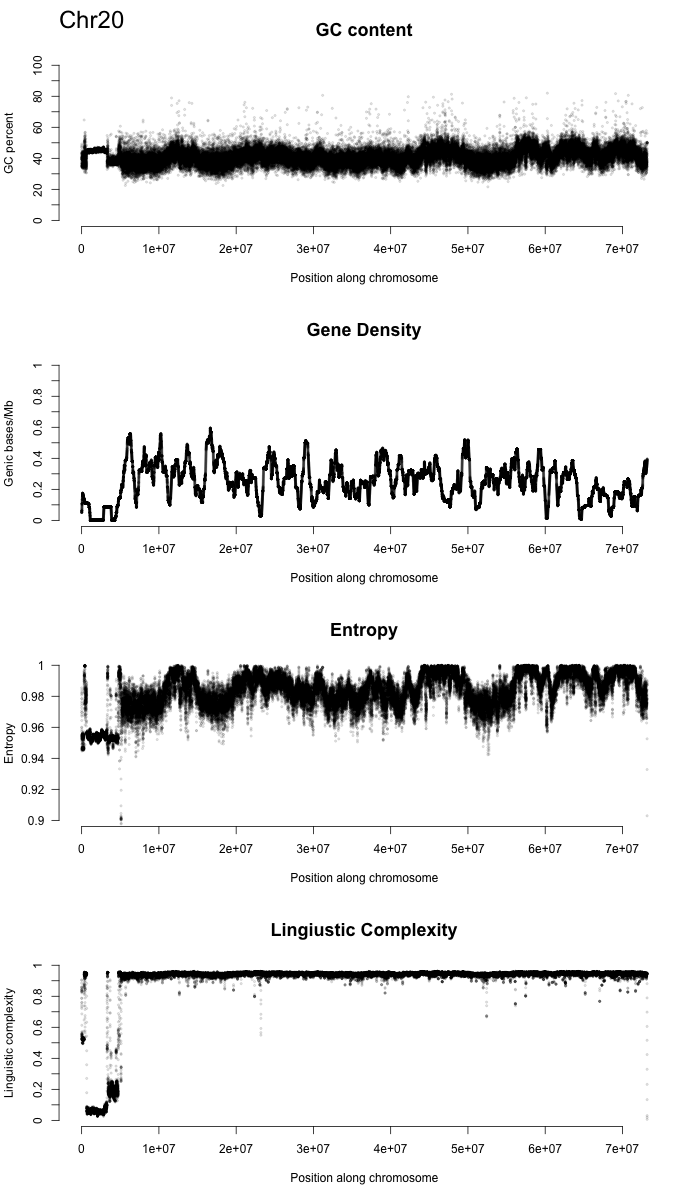

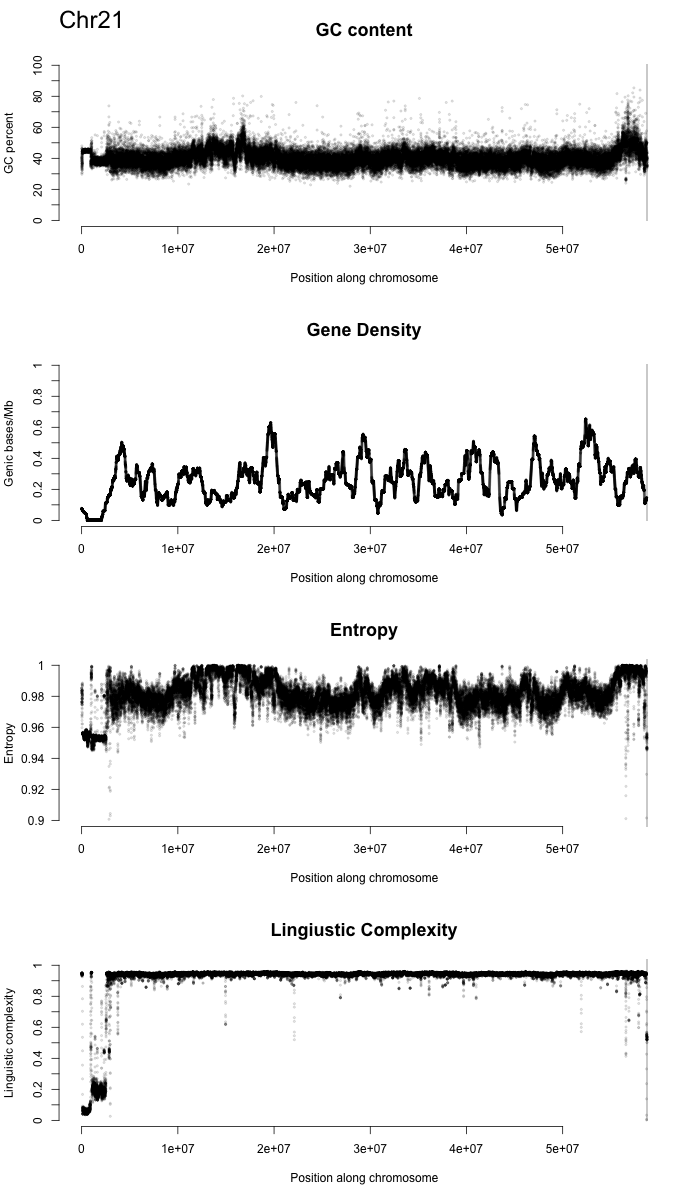

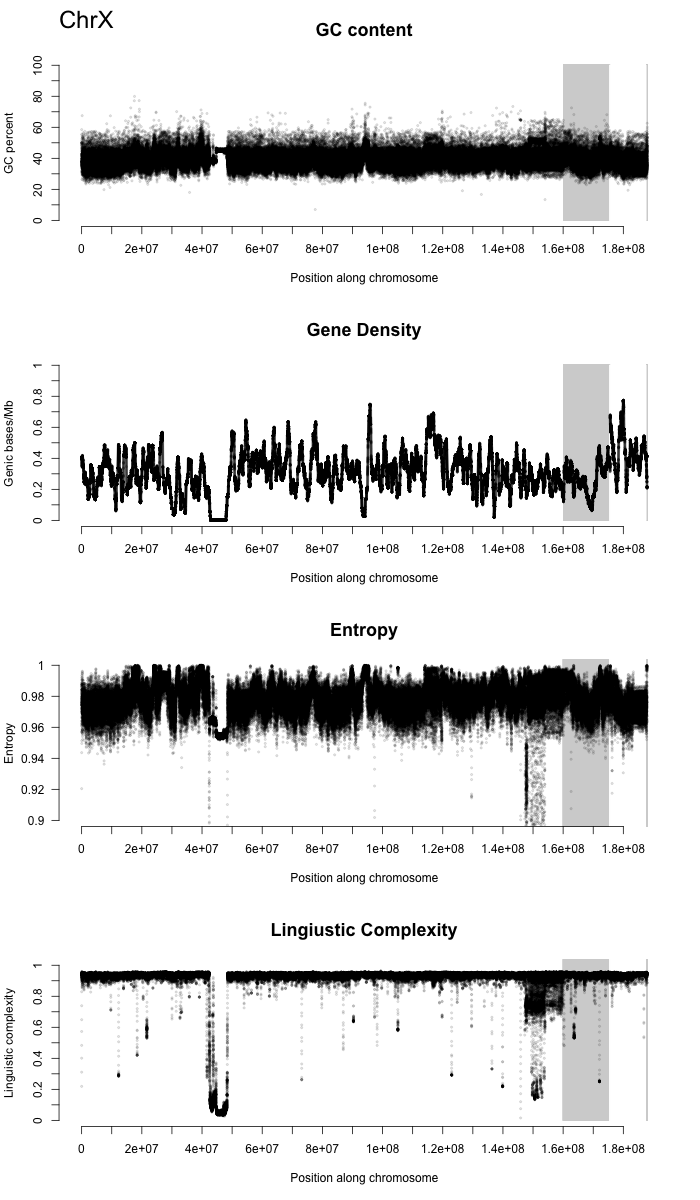

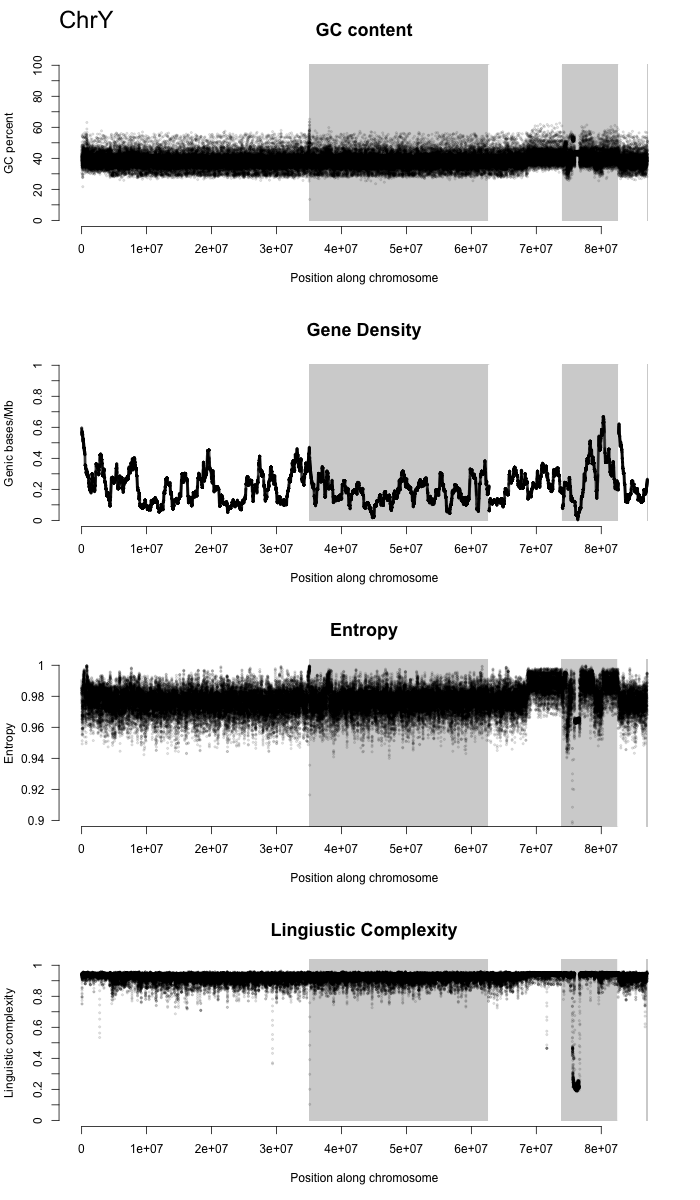

Figure S9: Comparison of two chromosome-scale *Meriones* *unguiculatus* genomes. Above is our assembly with centromeres and the expansion on Chromosome 5 marked with grey ovals. Below are the longest 24 scaffolds from DNAZoo’s Hi-C scaffolding of (Cheng et al. 2019)’s genome comprising 90.2% of the sequence. Except for a few large-scale inversions, the two genomes are predominately co-linear. Chromosome 13 is essentially absent from Cheng et al (2019)’s version as is the large repeat expansion on Chromosome 5. The short scaffolds of Cheng et al (2019)’s assembly are likely the unassembled fragments of Chromosome 13 but due to their length and repetitive nature, quality alignments could not be accurately identified.

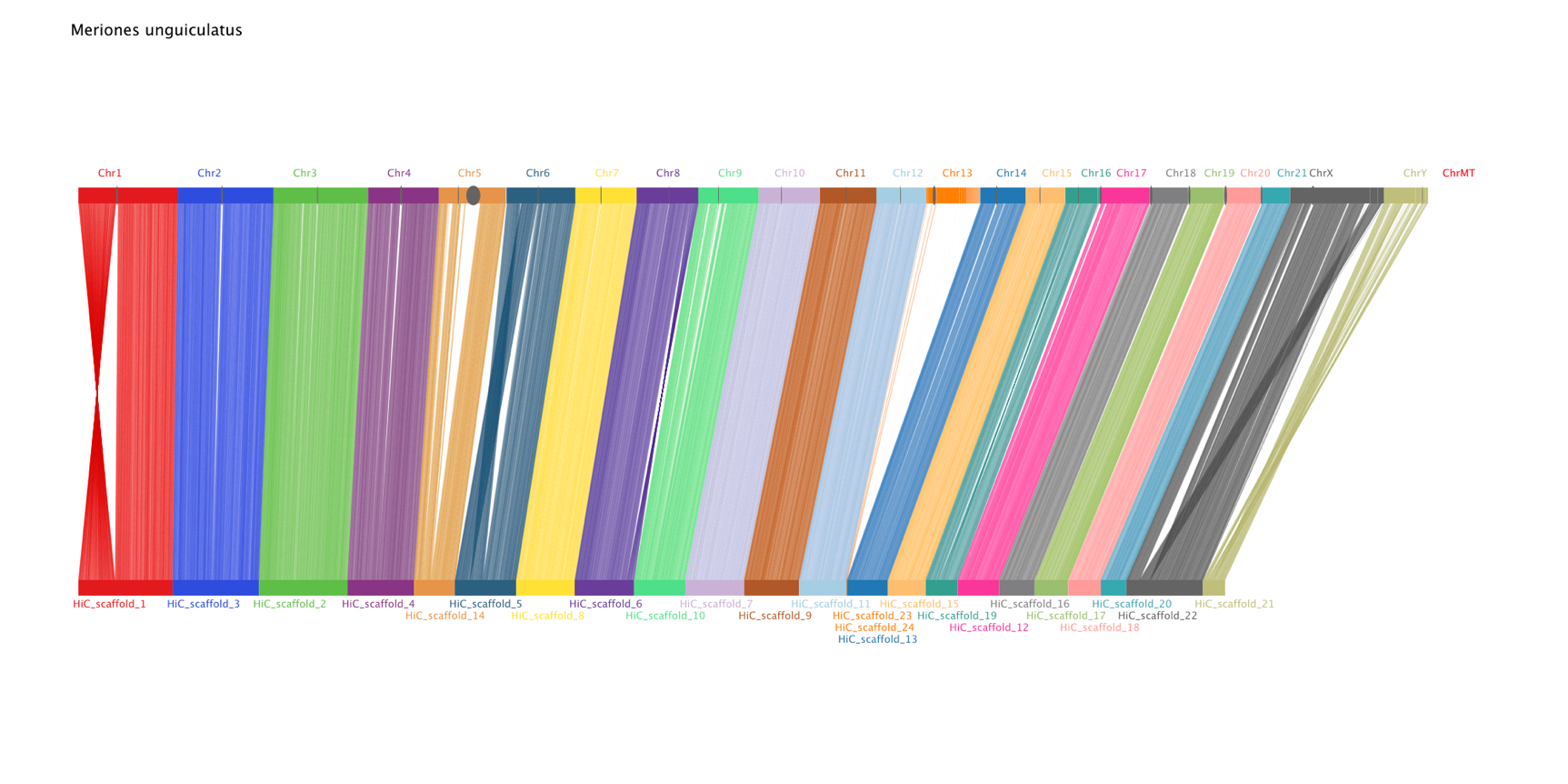

Figure S10: NTRprism of centromeres for each chromosome and the expansion of Chr5. Peaks show the frequency of repeats of each length. Often the program will identify series of repeats of X, 2X, 3X, 4X etc when the length of the base repeat is X. The list of numbers to the right of each plot is the lengths of the repeats with the most common at the top. Multiples of 6 are typically MsatA (the telomere repeat), multiples of 37 are MsatB, multiples of 127 are MsatC, and MsatD is on the Y chromosome at 1747bps. A representative sequence of each Msat are these (though note that there is variation):

>MsatA 6bp

TTAGGG

>MsatB 36bp

GAAGCAAAATTTTTGTGTTCTCAGTTGATTTAGTTG

>MsatC 127bp

TGAGATATATCACTTTTCATTTTTCCTGTATTCCTAATGGAGCGAGCGACGCACATGTGATTTACTTTACAAATTAATGATTTTCTTTTTTGTTCTTTCATTTGAAACTTGTTCTTGTGCTGTACAA

>MsatD 1747bp

AAGTAGAGAGGAGCTCCGTAAAAGGATTGCAGACAGGGCCCAGATACCATAAAGCAGTGTTTCTGAACTAAGCAGAGAAGAAGAGAGAGGGGGATCTCTAATAAACATAGCACAGAGCACAGAAAACTCTATTGTGGATATACGTGTAGGAGAACACAGAATAGTGTTACTGAACTTAGTGCGGAAGGACACCCAGTAGTGTTTCTGTCTGGGTTGATGCAAAAACAATTGTGTTCCTAAATAAAGAGACCATTAAACGAAAGGGAGTTATCCAGAAGCGAAGGAGTGGTTAGCACAGATTTGTTTCCAAGATAAGTACACGATTCCACAGAACGGAGTTTCGGAACTGAGGACAGAAGAAGAGATAGCAGTCAAGACACAGGAAGAATGTTATGTAGCAGAGAAGTTCACAGGAAAATGTTCCCAAGACGAGCACAGTTGAACAGTGTGATCAGGACCAAGGGTGGAAGAACACAGAGCAGTGTTTATTAACCTCGTTCTTCAATAATTTGAAGAAGTGCACTGCAACTAAGTCGGGACGAAAACCGAATTGTTATCTACCCACCACCGAAGAAGAGAGCACAGAAAGTTCTTTCAAACATGAGTACAGATGACCAAGGAACAGTGTGAACTAAGTCTAGCTGAATTAGGGCAGTGACGGTGAGCAAAAGAGAAACAGACAAACAAACAAATGAACAGTGTTAGATAGCAGAGAAGATCCCAGGAGAGTGTTCCCAGATGAGCACAGTGAGAGAGCGCTGCTCTTTTTTTTCTGCACGAAATACAGAGGAGCAGGTAGCAATGTTTTCTGAACCTCAGTTTCCAGAACCCAGGCATGTATCTGAACCAAGGAAGAAGAAAAGCTAAGTGTTGTCTTGCAGAGTCGACAAGAACACAGAAACGTGATTCCACCTTAAGGAAAGAGAGGGCTGAAAAGTGTTTTCAATTTAAGCACCTAAGAAACCACAGCAGTGTTTCTGCAATAAGTATTGCAGGAAAGAGGAGTGAGTGTGAAATCCAACAGAGTGGAGAAGAGCCCCGCCTACTGTGTCCAGGATAGCACAGAAGAGCACAGAAGACTGTTTTTGACCAAAGGAGGGACGAAGACACAGAATGCTTTGTACTATGTTGAGAGTAAAGTATGGAGTGTTTCCACAATCGGGACAGAAACACAGAAGAGGCTTGTTATCTAGCCCAGTAGAGATGAGGCCGAGAAGGGGATCAACGACATGGACCAAAGAGCACAGAACTGAGTCTCCATGATATGTTAGGGAAACAAGAGAAGAGCAGAGTAGACAACAGTTTCCAGGATACTAACAGAAAAGCACTGAGCAGAGTTTCCACAGGAGTACAAAAGATCTCACAGAAGGGTTCCTGAATGAAGTAGAGAAGAAGAAAGGAGATGGTTATTTAGCAAAGAGGGGAGCAGCCCACAGCAGCACTTAAGTGATCACTACAGAAAGAAAAAGACTGAACACTGTTCCTAAACGAAGGCAAGAAGAACTCGCAGCAGTGGATGTGAACAAACAAGAGAGGAAAAGAGAAGAGTGCTATCTAGACGAGAAGATCCCAGGAAAGTGTTTCAAAATTGGTACAGAAGAGAAGGAAGAGACAAGTGTTTCAGAAATGAGGGTATGTCACATGTTATTGTGTCTGAAGTAGGTAGATGAGAAATCTAAAGAGAGAAGGAAGCAGAGGCGTGTTATCTAGAGAAGTAGAGAATAGCGCTGAGAAGGGTTTTAATGGA

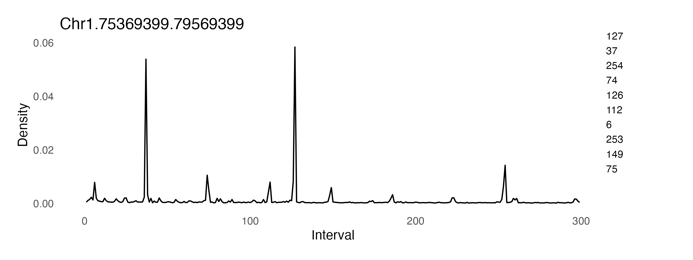

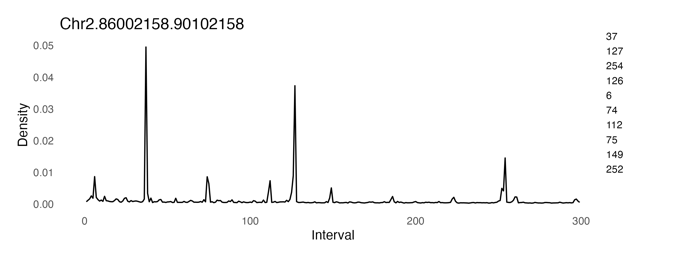

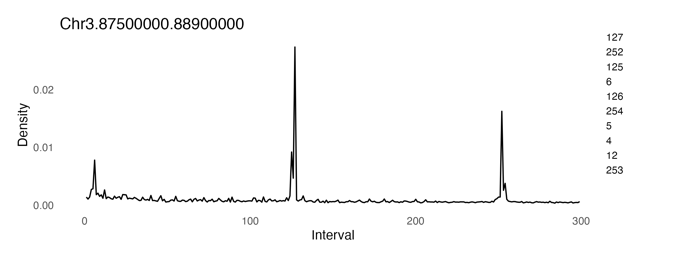

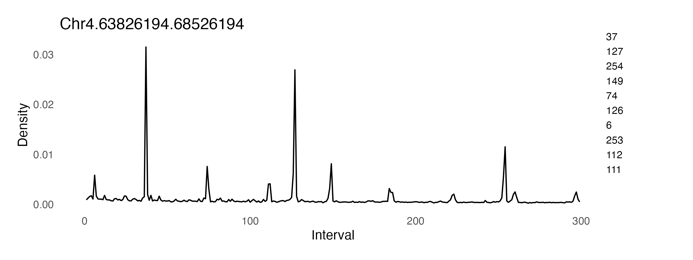

Chr5 centromere:
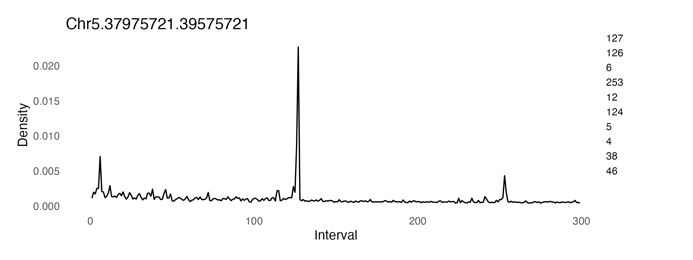

Chr5 expansion:
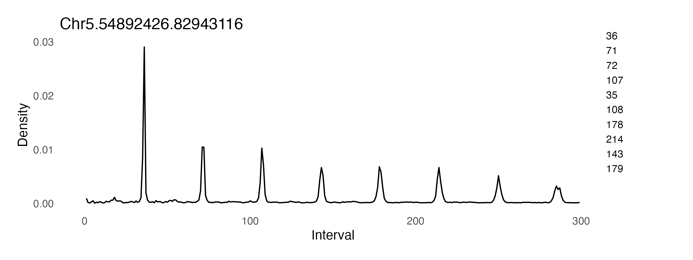

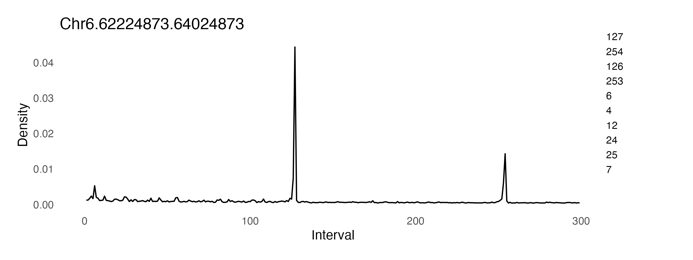

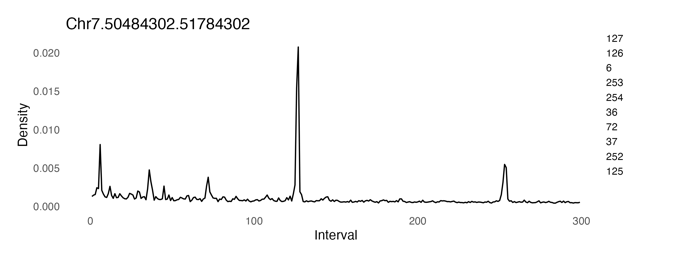

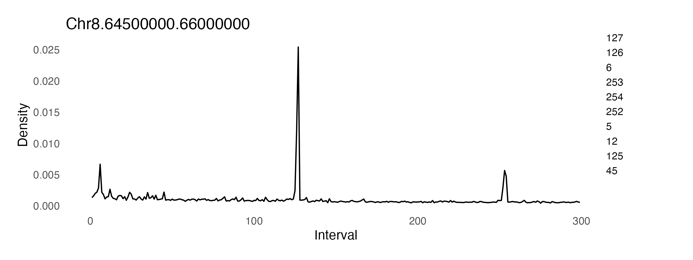

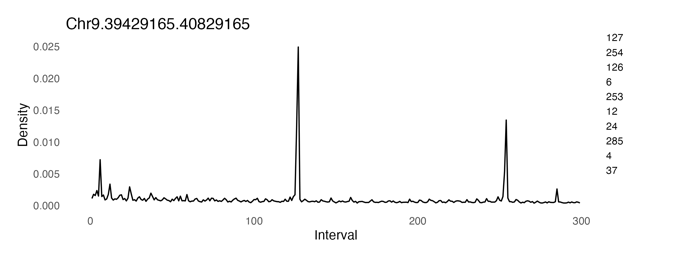

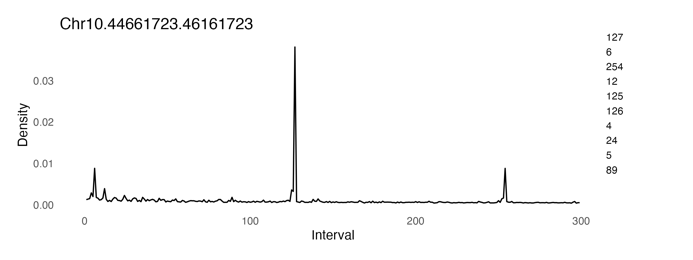

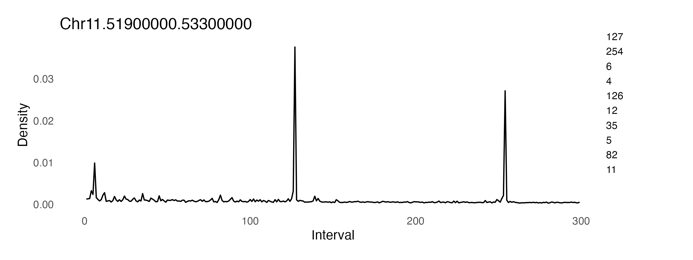

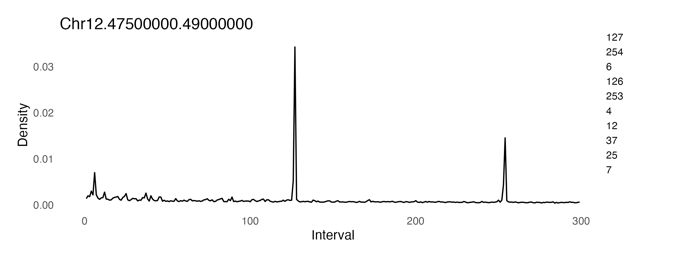

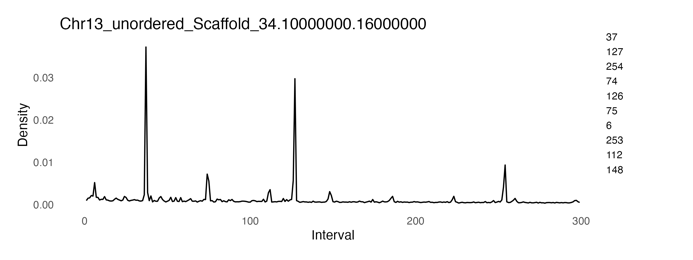

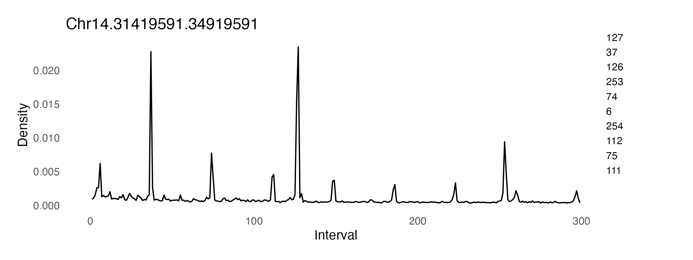

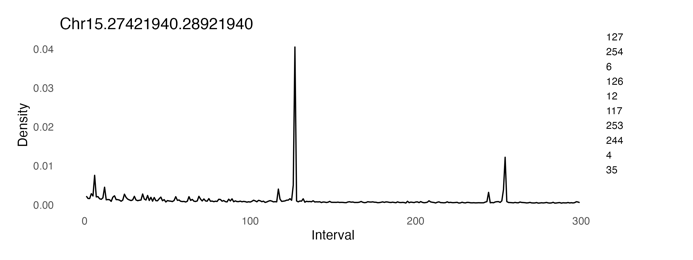

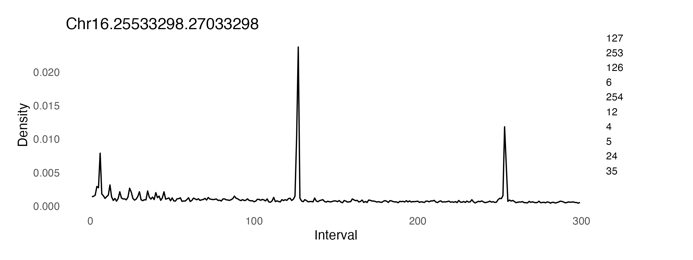

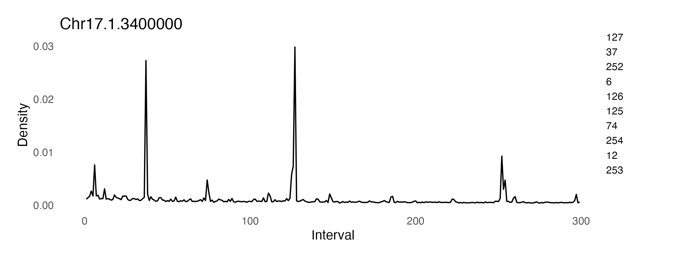

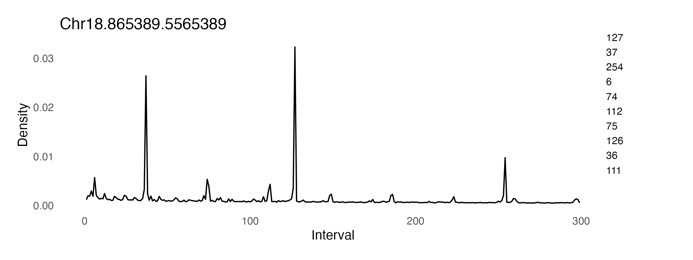

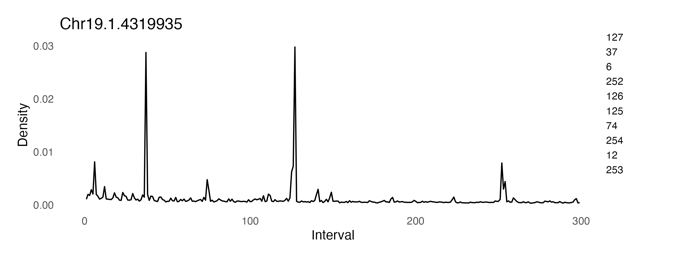

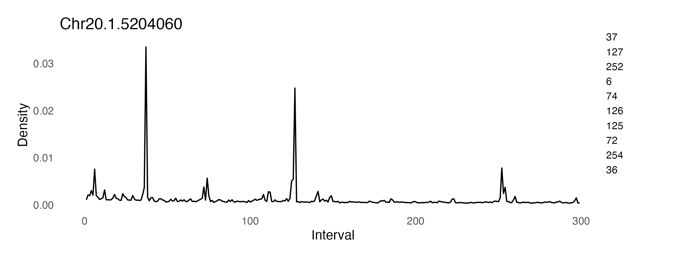

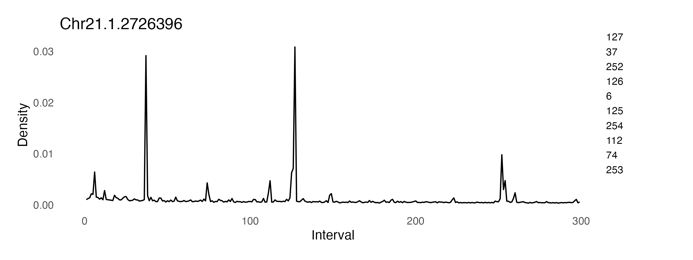

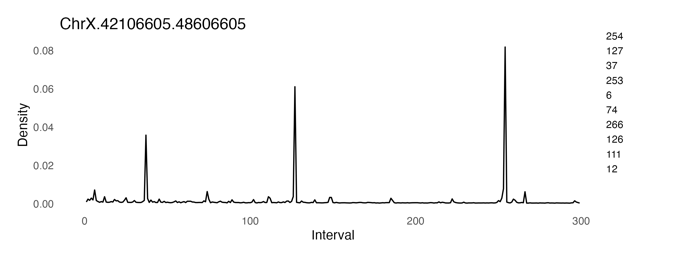

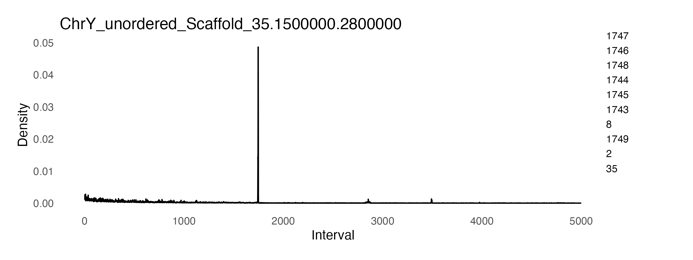

Figure S11: Recombination plots for all chromosomes showing the relationship between physical distance (in Mbp) and recombination rate (in cM/Mb) with recombination hotspots shaded in red.

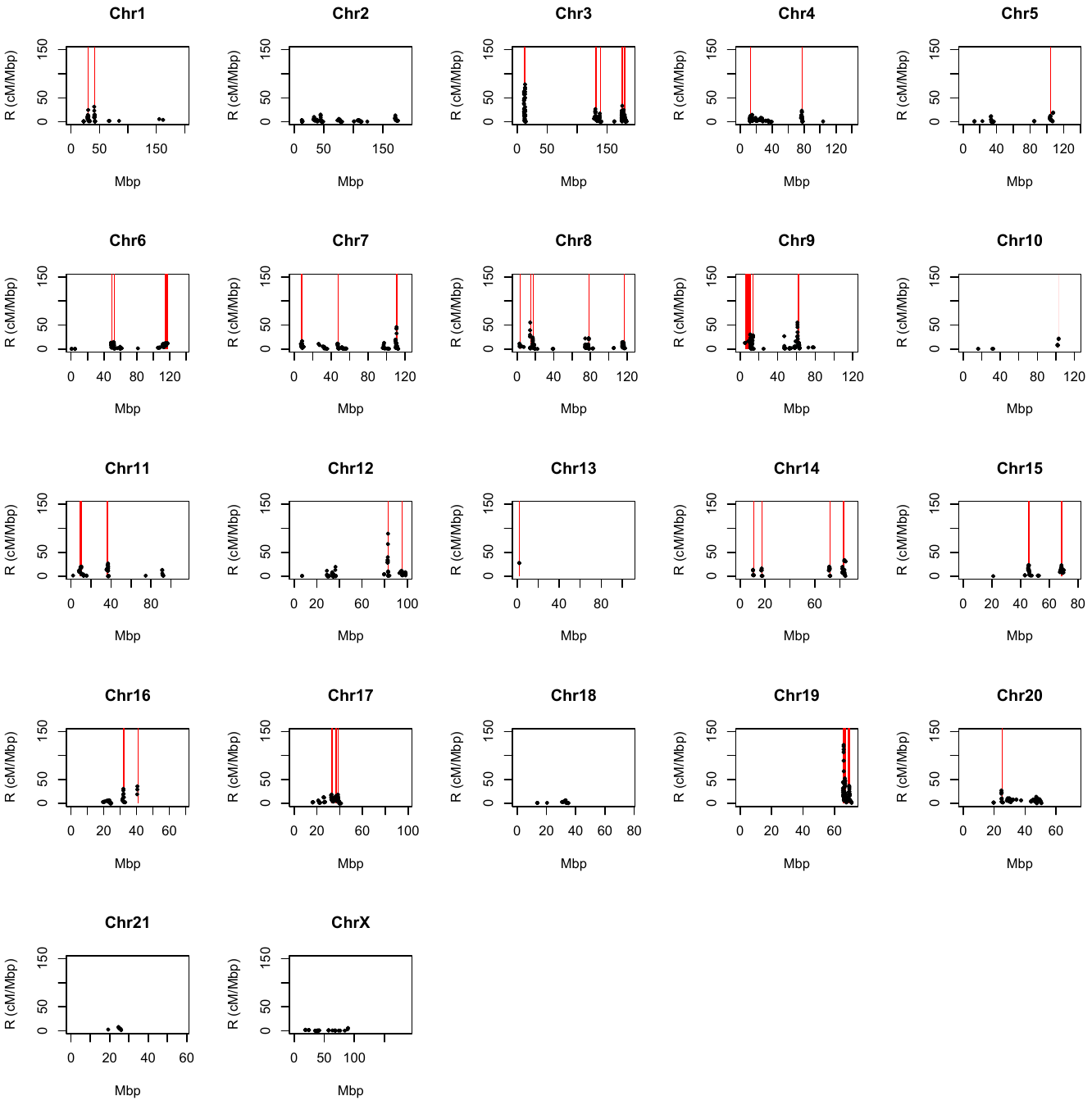

Figure S12: Marey Maps for all chromosomes showing the relationship between physical distance (in Mbp) and genetic distance (in cM) with recombination hotspots shaded in red.

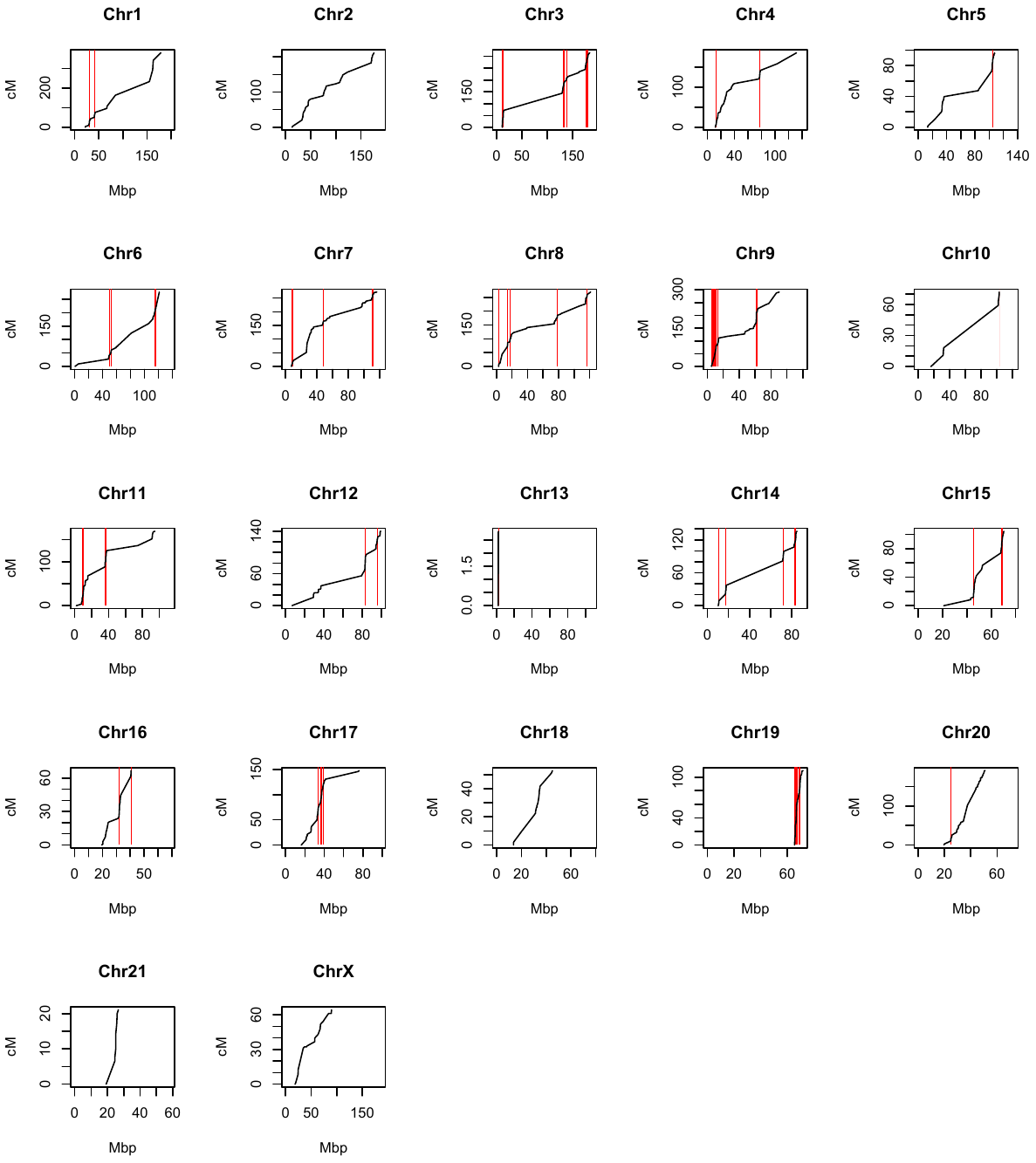

Figure S13:

Distribution of recombination hotspots across chromosomes. We identified 52 recombination hotspots spread across 18 of the 22 chromosomes (average = 2.36 hotspots per chromosome).

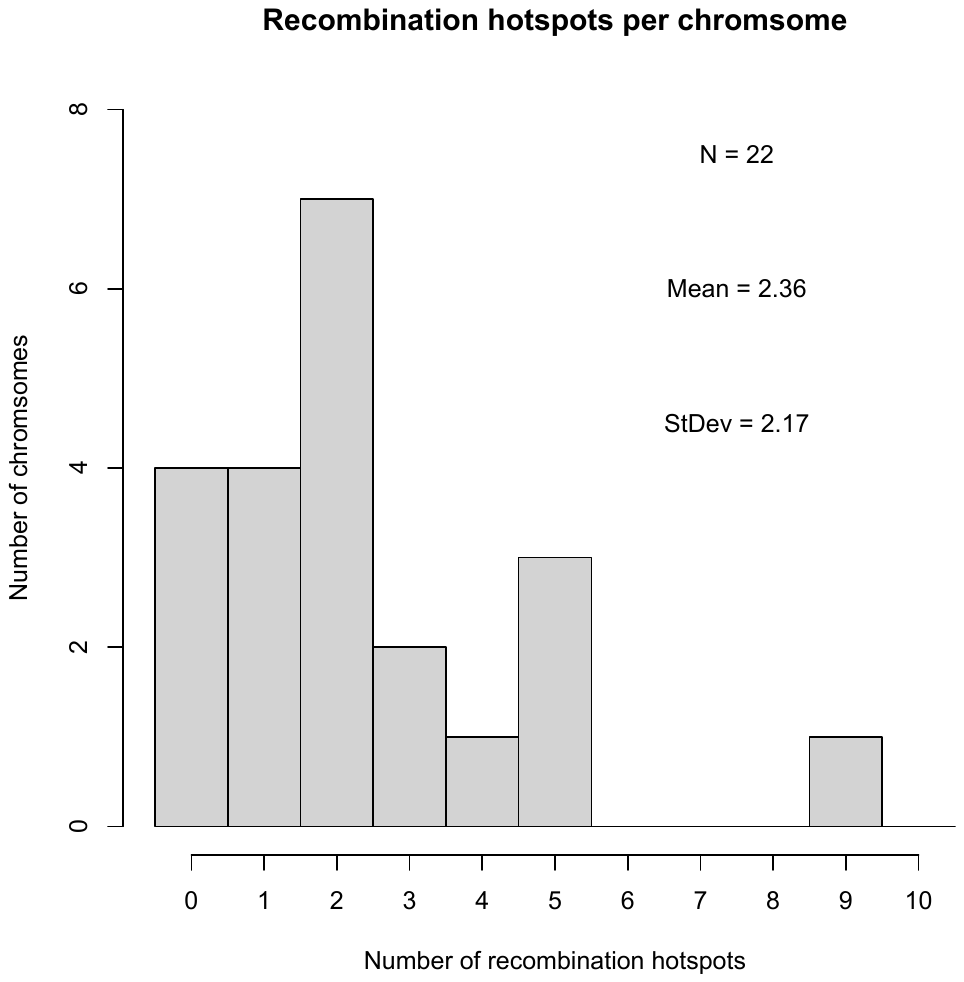

Figure S14: GC-rich genes occur closer to recombination hotspots than expected by chance (t-test, df = 383.2, t = 2.585, p = 0.01012).

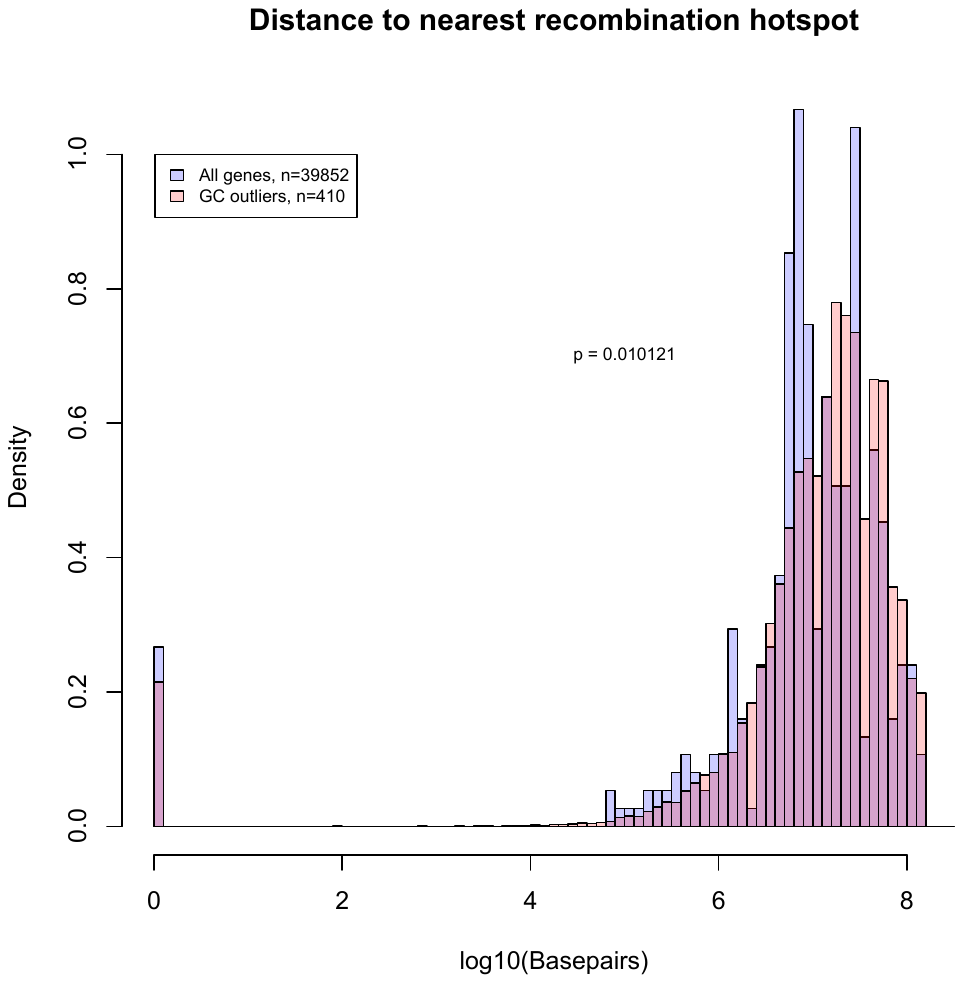

Figure S15: GC rich genes are found closer to the telomere end of chromosome arms than expected by chance t-test, df = 418, t = -14.26, p < 0.000001.

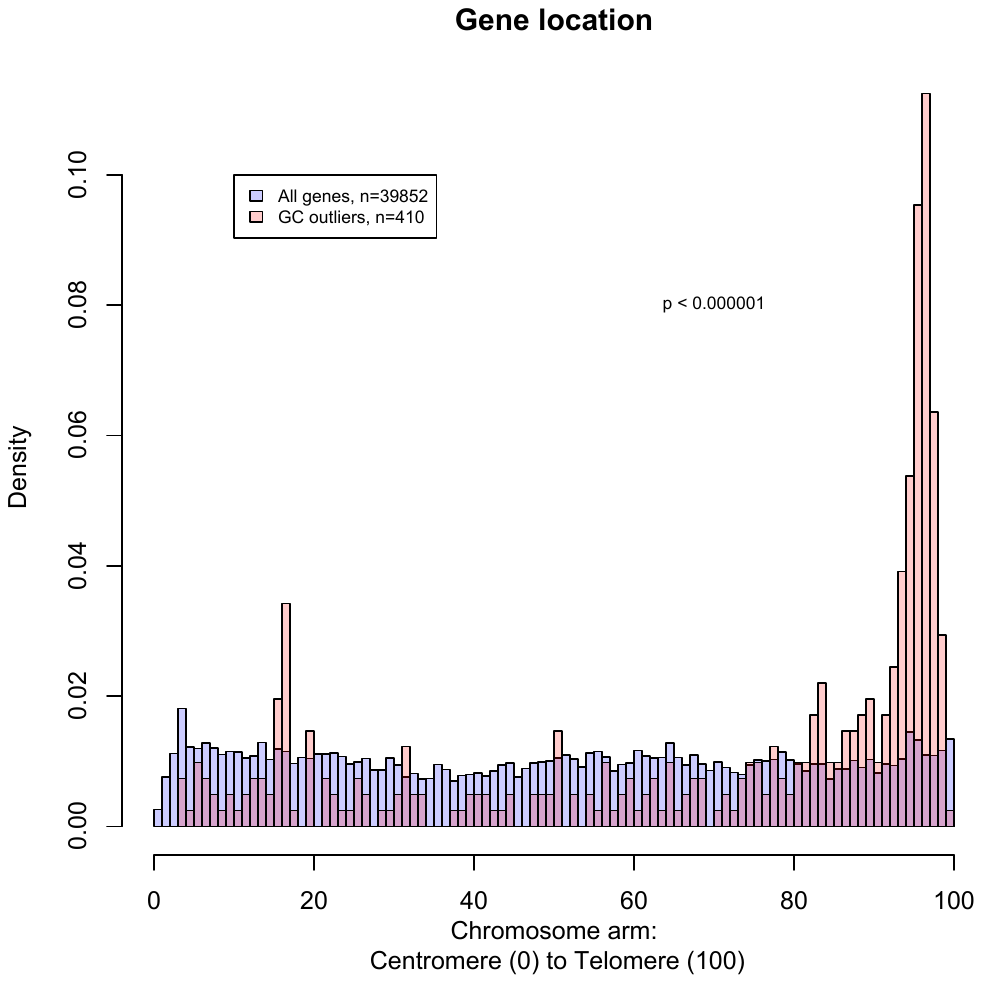

Figure S16: Number of telomere repeats in telomere repeat arrays. As the telomere repeat is only 6bp long it is found throughout the genome. All repeats were identified with the program TandemRepeatFinder and the parameters: 2 7 7 80 10 50 2000 -d -h -l 5. Then telomere repeats were extracted using a search for the telomere repeat sequence “MsatA” which is ‘TAACCC’ and allowing for either orientation and any start position (i.e.: “TAACCC”, and “AACCCT”, and “ACCCTA”, etc…). We defined *interstitial telomere sites* as those with >70 tandem copies of the repeat (to the right of the red line) as that seemed a natural breakpoint in the histogram and coincided with predictions of the number of interstitial telomere sites identified by FISH in (de la Fuente et al 2014).

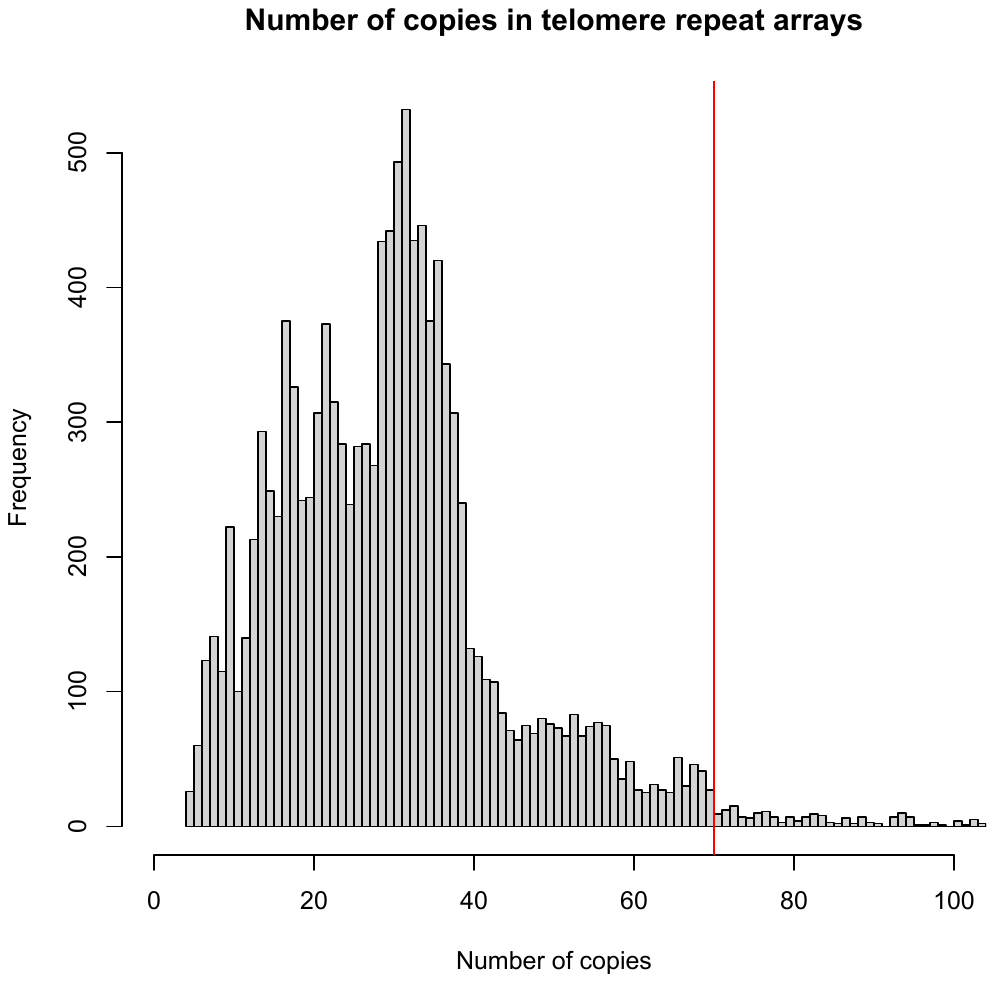

Figure S17: GC-rich genes are clustered nearer telomere repeats (interstitial or otherwise) than expected by chance (t-test, df = 418.6, t = 7.876, p < 0.000001).

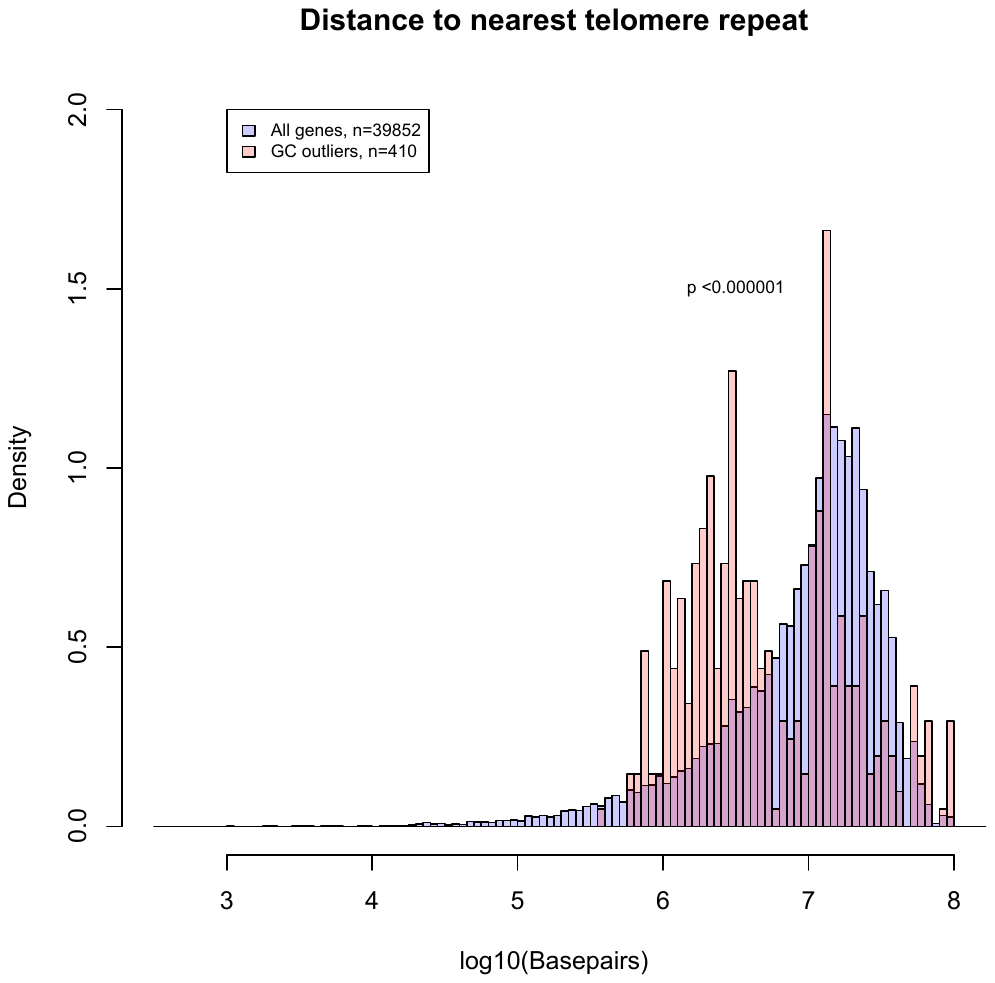

Figure S18: schematic of the gerbil genome with various features marked including centromeres (blue), high GC-content genes (red) recombination hotspots (purple), and interstitial telomere sites with >70 copies in the array (green).
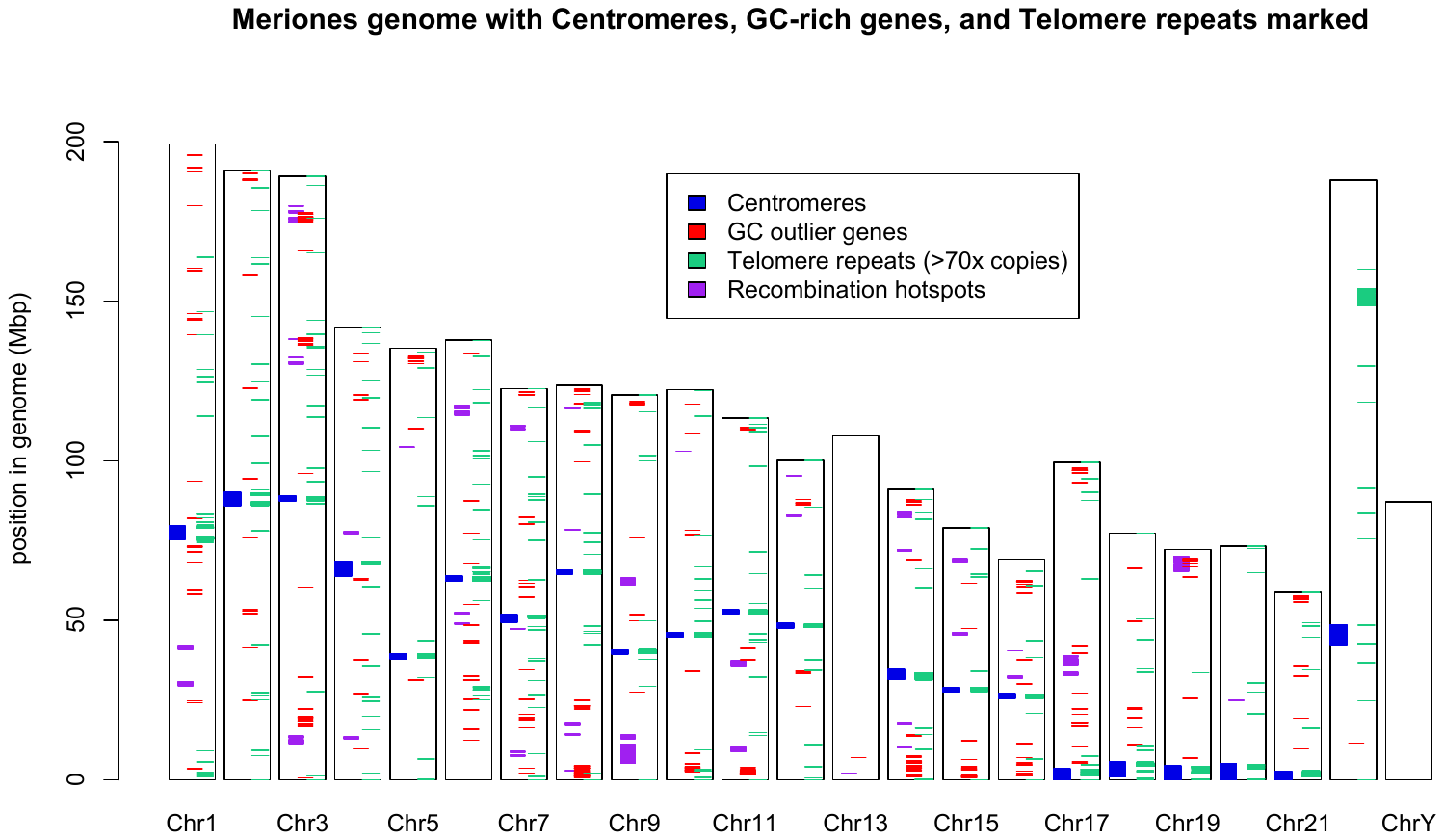

Figure S19: Whole genome alignment showing *Meriones unguiculatus* Chromosome 5 (top) and *Psammomys obesus* Chr 10 (bottom). The location of the centromeres and the repeat expansion are shown as grey ovals. The repeat expansion on *Meriones* Chr5 is at the same location as the centromere of *Psammomys* Chr10. This suggests that the repeat expansion may have been an ancient centromere that has expanded, possibly due to centromere drive. *Psammomys* genome helpfully provided by David Thybert.

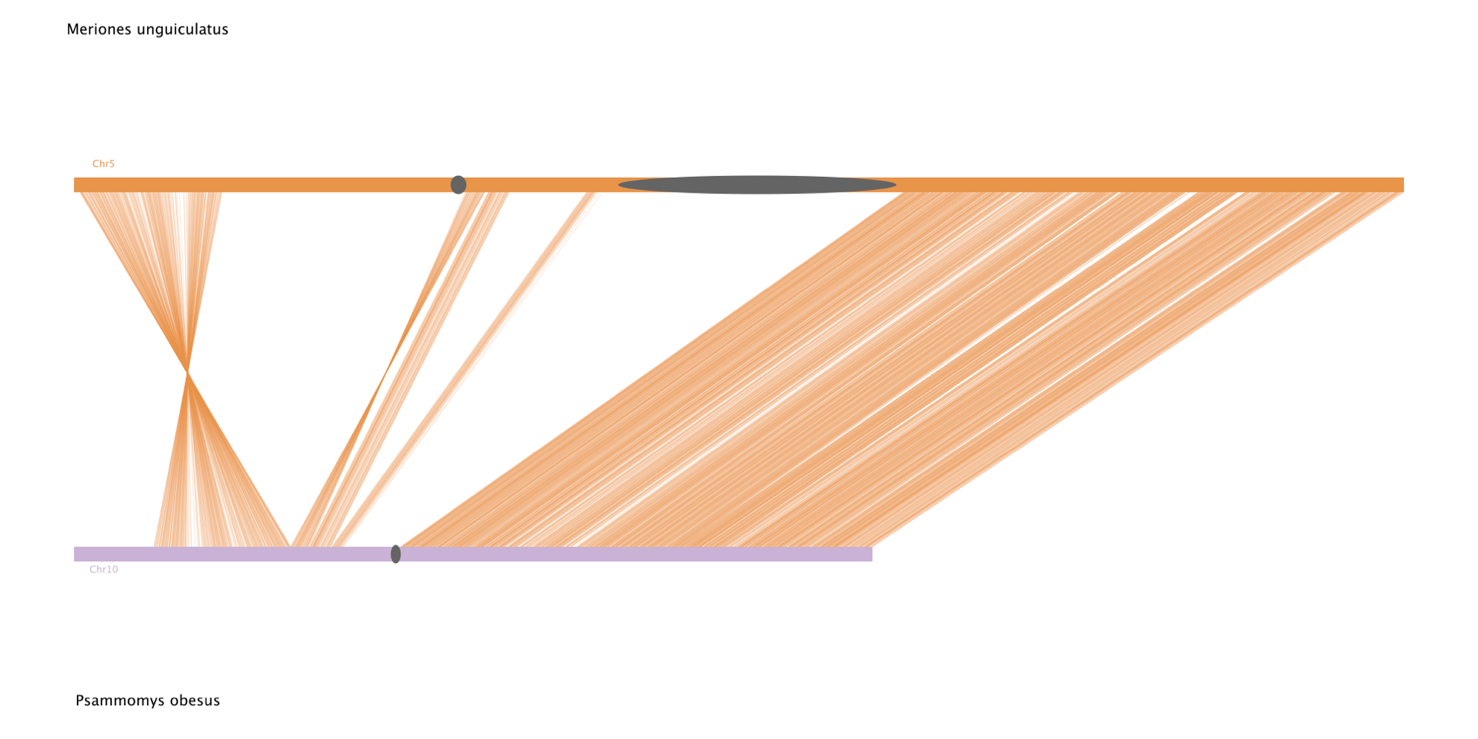

**Detailed Methods**

Animal care and tissue collection

Male Tumblebrook Farm strain Mongolian gerbils from the colony at Bangor University (Brekke et al. 2018) were euthanised using a Schedule 1 method in accordance with EU Directive 2010/63/EU and the Animals (Scientific Procedures) Act 1986. Animal use was reviewed and approved by the Bangor University Animal Welfare and Ethical Review Board. Fresh liver, kidney, and testis were dissected and snap-frozen in liquid nitrogen, then stored at -80°C and whole spleens were dissected into pre-warmed RPMI 1640 buffer (ThermoFisher) for cell culture. Whole blood was collected into EDTA tubes BD Sciences 366450) and stored at 4°C until shipping.

Cell culture and chromosome sorting

Chromosomes were harvested from the gerbil fibroma cell line IMR-33 (ECACC General Cell Collection catalogue number 96020931) after being arrested in mitosis with Colcemid. The cells were lysed in a hypotonic solution and the chromosomes in suspension were sorted using a BD Influx cell sorter (Becton Dickinson, San Jose, CA) as described by (Kuderna et al. 2019). 17 separate clusters were identified and the used for sorting resulting in 6 pools with two chromosomes each and 11 pools with a single chromosome. After chromosomes were sorted, we dialysed each pool using a Pur-A-Lyzer Maxi Dialysis kit (Sigma, PURX50005-1KT) following the manufacturer's instructions to remove any dye that remained bound to the DNA from sorting.

DNA and RNA Sequencing

PacBio HiFi sequencing was performed by the Earlham Institute (Norwich, UK), using DNA extracted from frozen liver tissue. High molecular weight DNA was extracted with the Nanobind Tissue kit (Circulomics), with quality assessed using an Agilent FEMTO-Pulse, with size selection of fragments between 18Kb and 19Kb in size with a SageELF. HiFi sequencing used 3 SMRT cells on a PacBio Sequel II, analysed with the CCS analysis pipeline (SRR18362962). We sequenced 88,071,091,902 base pairs and generated 4,814,749 CCS reads with a quality greater than or equal to Q20. These had an average length of 18,291bp and amount to 34x coverage.

OmniC sequencing was performed by Dovetail Genomics (California, USA), using DNA derived from frozen liver samples. 262,974,243 pairs of 151bp OmniC reads were sequenced (SRR18362944) corresponding to 38x coverage.

Whole fresh blood was sent to the DeepSeq facility (Nottingham, UK) for Oxford Nanopore PromethION sequencing and BioNano Optical Mapping. DNA was extracted in parallel with the Circulomics UHMW DNA extraction protocol (EXT-BLU-001) for PromethION sequencing and the Circulomics HMW DNA Extraction Protocol (EXT-BLH-001) for BioNano optical mapping. We sequenced 1,210,550 ultra-long reads of average length 39387bp and 2,550,000 long reads averaging 19,850bp via the PromethION (SRR18362939).

The sorted chromosomes were sequenced with Illumina MiSeq at Bangor University, and we generated 19,764,484 read pairs of 151bp paired end reads (SRR18362948, SRR18362952, SRR18362951, SRR18362947, SRR18362946, SRR18362945, SRR18362940, SRR18362958, SRR18362956, SRR18362955, SRR18362954, SRR18362949, SRR18362957, SRR18362953, SRR18362960, SRR18362959, SRR18362941).

RNA extraction and sequencing was done by GeneWiz (New Jersey, USA), using kidney and testis from three individuals. We received 177,016,012 151bp paired-end reads (SRR18362961, SRR18362937, SRR18362950, SRR18362943, SRR18362942, SRR18362938).

Genome assembly and annotation

We used the HiFiASM assembly software (Cheng et al. 2020) to build the first iteration genome using the PacBio HiFi reads with the flag -l 1 to lightly purge duplicates. This initial assembly was then scaffolded using the OmniC read-pairs and the HiRise pipeline. To the OmniC-scaffolded assembly we aligned the raw reads from the genetic map (Brekke et al. 2019) using bwa (Li and Durbin 2009) and ran the Stacks2 (Rochette et al. 2019) pipeline followed by R/qtl (Broman et al. 2003; Arends et al. 2014) to build a genome-guided genetic map. This map informed additional merges of scaffolds which we did using the custom script assemble_genome_and_recoordinate_gff.py (Supplemental Material 4). At this point there was a single linkage group per chromosome (omitting the Y) and so the scaffold designation was dropped from the name of those fasta entries. Thus anything named, for instance, simply “Chr13” is tied to a linkage group whereas the scaffolds without genetic markers kept the longer names of the form “Chr13_unplaced_Scaffold_28”. To further assemble the reference, we aligned the Oxford Nanopore ultra-long reads using minimap2 (v2.17)(Li 2018) which also suggested additional merges. We built a hybrid assembly using the Bionano optical mapping data but it did not suggest any further merges and was not used in any analysis.

Our annotation was done on the Omni-C scaffolded version of the genome using the RNAseq data from GeneWiz (New Jersey, USA). Repeat families found in the genome assemblies of *Meriones unguiculatus* were identified de novo and classified using the software package RepeatModeler (v2.0.1)(Flynn et al. 2020). RepeatModeler depends on the programs RECON (v1.08)(Bao and Eddy 2002) and RepeatScout (v1.0.6)(Price et al. 2005) for the *de novo* identification of repeats within the genome. The custom repeat library obtained from RepeatModeler were used to discover, identify, and mask the repeats in the assembly file using RepeatMasker (v4.1.0)(Tarailo‐Graovac and Chen 2009). Coding sequences from *Meriones unguiculatus, Psammomys obesus*, *Mus musculus*, *Rattus norvegicus* and *Peromyscus maniculatus* were used to train the initial *ab initio* model for *Meriones unguiculatus* using the AUGUSTUS software (v2.5.5)(Keller et al. 2011). Six rounds of prediction optimisation were done with the software package provided by AUGUSTUS. The same coding sequences were also used to train a separate *ab initio* model for *Meriones unguiculatus* using SNAP (v2006-07-28)(Korf 2004). RNAseq reads were mapped onto the genome using the STAR aligner software (v2.7)(Dobin et al. 2013) and intron hints generated with the bam2hints tools within the AUGUSTUS software. MAKER, SNAP and AUGUSTUS (with intron-exon boundary hints provided from RNA-Seq) were then used to predict for genes in the repeat-masked reference genome. To help guide the prediction process, Swiss-Prot peptide sequences from the UniProt database were downloaded and used in conjunction with the protein sequences from *Meriones unguiculatus, Psammomys obesus, Mus musculus, Rattus norvegicus* and *Peromyscus maniculatus* to generate peptide evidence in the Maker pipeline. Only genes that were predicted by both SNAP and AUGUSTUS softwares were retained in the final gene sets. To help assess the quality of the gene prediction, AED scores were generated for each of the predicted genes as part of the MAKER pipeline. Genes were further characterised for their putative function by performing a BLAST search of the peptide sequences against the UniProt database. tRNA were predicted using the software tRNAscan-SE (v2.05)(Chan and Lowe 2019). The gff annotation file was then re-coordinated along with each subsequent merging of scaffolds through the remaining assembly steps using the custom python script assemble_genome_and_recoordinate_gff.py (Supplemental Material 4).

The annotation pipeline described above did a perfunctory identification of repetitive elements as a step towards finding a high-quality set of genes, but to assemble and curate a high-quality list of repetitive sequences, we used the EarlGrey pipeline (Baril et al. 2022). This was configured with Dfam (version 3.4) (Hubley et al. 2016) and RepBase (release 20181026) (Jurka et al. 2005; Kapitonov and Jurka 2008), specifying known repeats from *Rodentia* (-r rodentia).

Chromosome Assignment

Scaffolds from the final genome were assigned to chromosomes by the parallel approaches of sequencing each sorted chromosome pool and also using each pool as a FISH probe. Illumina reads from the pool sequencing were aligned to the assembly with bwa (Li and Durbin 2009). For the alignment of each chromosome pool, we counted the number of reads mapping to each scaffold and calculated the reads mapped per scaffold length. Each scaffold then has a read-mapping density from each pool making it possible to associate every scaffold with the pool to which it belonged. We calculated the 99.99% confidence interval of the read mapping density and the scaffold was assigned to the pool that fell outside the confidence interval. In 30 of the 194 cases, a scaffold assigned to either no pool or to multiple pools and we marked these as 'unknown'. Unknown scaffolds comprise 1,588,872 bases, 0.066% of the total genome.

The final link in the chain connecting the karyotype with the chromosomes was made using FISH probes created from the sorted pools. The chromosomes in the gerbil karyotype were named by Weiss (Weiss et al. 1970) who defined the pattern of G-bands on each chromosome. Spleen cells were cultured as per (Yang et al. 2017) in complete RPMI (i.e. RPMI with fetal calf serum, penicillin, and streptomycin) with added EPS to stimulate immune cell growth at 37C and 5% CO_2_ for 48 hours after which colchicine was added to arrest the cells in metaphase. After an hour, the cells were spun down and resuspended in fixative and stored at -20C. Metaphase spreads were made by dropping 14 μl of cell suspension on a glass slide and drying at high humidity while floating in a 55C water bath. We created FISH probes from each chromosome pool and hybridized them to a chromosome spread, thereby linking the banding pattern of each chromosome with a pool. FISH probes were made and hybridized to the metaphase spreads following (Murchison et al. 2012)

Male gerbils have 23 unique chromosomes (21 autosomes, an X, and a Y) and so we expected six of the 17 pools to have multiple chromosomes which is what we found. Eleven of the pools had a single chromosome while six pools included two chromosomes. The methods we used to distinguish the chromosomes in these six pairs are laid out in full in the supplemental material, see Figures S2-S7 and Tables S2 and S3.

Thus, the final version of gerbil genome presented here is the result of PacBio HiFi reads assembled with HiFiAsm (Cheng et al. 2020), scaffolded with OmniC paired reads, annotated with kidney and liver RNAseq data, further scaffolded with a high-density SNP-based genetic linkage map and then Oxford Nanopore ultra-long reads, and assigned to physical chromosomes using chromosome sorting and chromosomal FISH.

Genome Analysis

GC content and gene density were analyzed for every scaffold in the genome in sliding windows using the custom script Calc_R_GC_Gene_density.py (see supplemental code). The window size for GC content is 1,000bp while the window size for gene density is 1Mb in both cases the windows progressed by 1kb across each scaffold. Recombination rate was estimated by taking a sliding window of 8 genetic markers and calculating the slope of the regression of their genetic position against their physical position. As the sliding window progressed by a single marker, each inter-marker region had eight rates associated with it and these were averaged to get the recombination rate of each inter-marker region. The genome-wide recombination rate was calculated by taking the mean of the rate of each region weighted by the length of region. Recombination hotspots were identified by identifying every region whose rate was greater than 5 times the genome average and adjacent regions were merged. Entropy and Linguistic complexity were calculated using the program NeSSie (Berselli et al. 2018) using a sliding window with size 10k and a step of 1k as recommended.

We identified centromere location by visually identifying the trough in the linguistic complexity data of each chromosome. To understand the fine-scale structure of each centromere, we extracted the region and used the program NTRprism (Altemose et al. 2022) to identify the lengths of the different repeats and TandemRepeatFinder(Benson 1999) to identify the sequence of repeats of each length, the data from which forms the basis of the coloured centromere call-outs in Figure 3D. TandemRepeatFinder also served to identify the location of telomere repeats along the length of the chromosome. We identified interstitial telomere sites as those with at least 70 tandem copies of the telomere repeat.

The locations of the 387 GC outlier genes in *Meriones* identified by (Pracana et al. 2020) were extracted from the annotation file. Due to some gene duplications this resulted in 410 genomic locations in our assembly. We tested whether these locations were more clustered than chance by drawing 410 random genes from the genome 1,000,000 times and calculating the average that separated each from its closest neighbor in the set, and calculated a p-value by taking the proportion of the permutations that had a lower average distance than the observed set. We calculated how close they were to recombination in two ways, first by a t-test comparing the distribution of all genes’ proximity to recombination hotspots with the distribution of the outlier genes and second by randomly drawing 410 genes 1,000,000 times and calculating the average distance to the nearest hotspot for the draw. A p-value was calculated for the permutation test as described above. A similar pair of tests was done to evaluate whether the outlier genes were non-randomly placed along a chromosome arm and whether they were more closely located to telomere repeats (interstitial and normal) than expected by chance. For the location along a chromosome arm we transformed physical position of each gene in to a percentage going from the centromere at 0 to the telomere at 100 in order to compare chromosome arms of different sizes.

A self alignment was made for each scaffold that assigned to chromosome 13 as well as a few selected autosomes (10, 16, and 21) and the Y chromosome. The self-alignment was done with mummer4 (Kurtz et al. 2004) using the “maxmatch” and “nosimplify” parameters to identify repetitive elements. Mummer was also used to compare our genome with both the other chromosome-scale *Meriones* assembly (Cheng et al. 2019)and an unpublished chromosome-scale *Psammomys obesus* assembly (provided by David Thybert) using the “mum” parameter.

Meiotic chromosome preparation and immunofluorescence

We obtained preparations of male meiotic chromosomes as previously described (Peters et al. 1997; Fuente et al. 2007) Briefly, a cell suspension was made in PBS from whole testicles by rubbing seminiferous tubules with the help of two forceps. Then, cells were transferred to a 10mM sucrose solution and spread over glass slides previously covered with paraformaldehyde 1% in distilled water (pH 9,5) containing 0.15% of Triton X-100. After two hours standing horizontally on a humid chamber, slides were washed in distilled water with 0.04% Photoflo, air dried and stored at -80ºC until use. For immunofluorescence, slides were incubated overnight at 4ºC with the following primary antibodies diluted 1:100 in PBS: goat anti-SYCP3 protein of the synaptonemal complex (Santa Cruz 20845); rabbit anti histone H3 trimethylated at lysine 9 (H3K9me3) (Abcam 8898), as a marker of heterochromatin; human anti-centromere (Antibodies Incorporated 15-234); and mouse anti-MLH1 (Pharmingen 550838), as a marker of meiotic crossovers. After washing three times in PBS slides were incubated for one hour at room temperature with secondary antibodies conjugated with Alexafluor 350, Alexafluor 488, Cy3 or Cy5, diluted 1:100 in PBS, all of them from Jackson ImmunoResearch Laboratories. After three washes in PBS slides were mounted with Vectashield. Observations were made in an Olympus BX61 microscope equipped with appropriate fluorescence filters and an Olympus DP72 digital camera. Them, images were treated witha Image J.

Bivalents 5 and 13 were identified in pachytene spermatocytes owing to the presence of an interstitial H3K9me3 region or by a complete labeling with this histone marks, respectively. To assess the position of MLH1 foci along the bivalents, we measured the length of the synaptonemal complex of these two bivalents using the free hand tool in ImageJ. The distance of centromeres and MLH1 foci from the tip of the short arm of the bivalents were recorded in the same way. Then the position of MLH1 foci was normalized against the length of the corresponding bivalent, yielding a position between 0 (the proximal end) and 1 (the distal end). The position of all foci was presented in cumulative frequency chart. A total of 83 spermatocytes from two different individuals were scored.

Data availability

All sequencing data and the genome is available under SRA BioProject PRJNA397533.

PacBio HiFi: SRR18362962.

Illumina OmniC: SRR18362944.

Oxford Nanopore PromethION: SRR18362939.

Illumina MiSeq sorted chromosome sequencing: SRR18362948, SRR18362952, SRR18362951, SRR18362947, SRR18362946, SRR18362945, SRR18362940, SRR18362958, SRR18362956, SRR18362955, SRR18362954, SRR18362949, SRR18362957, SRR18362953, SRR18362960, SRR18362959, SRR18362941.

Illumina RNAseq from testis and kidney: SRR18362961, SRR18362937, SRR18362950, SRR18362943, SRR18362942, SRR18362938.

This Whole Genome Shotgun project has been deposited at DDBJ/ENA/GenBank
under the accession JAODIK000000000. The version described in this paper is version JAODIK010000000.

The genetic map, a vcf of the genetic markers and their genotypes in the mapping panel, the gff of the gene annotations, the gff of the repetitive element annotations, and “Supplemental_Material 3_codebase.zip” can be found in the Dryad repository here: Brekke, Thomas D (2022), Data for "The origin of a new chromosome in gerbils", Dryad, Dataset, <https://doi.org/10.5061/dryad.1vhhmgqws>.

Reviewers may find these data files ahead of publication here: https://datadryad.org/stash/share/R5vtycW8DE6euNZJEe26JvJIVTmCEaJVo9SQpfXWAJk
