## Supplementary figures and images for "A chromosome-assigned Mongolian gerbil genome with sequenced centromeres provides evidence of a new chromosome"

### Supplemental material 2 - genome sketch

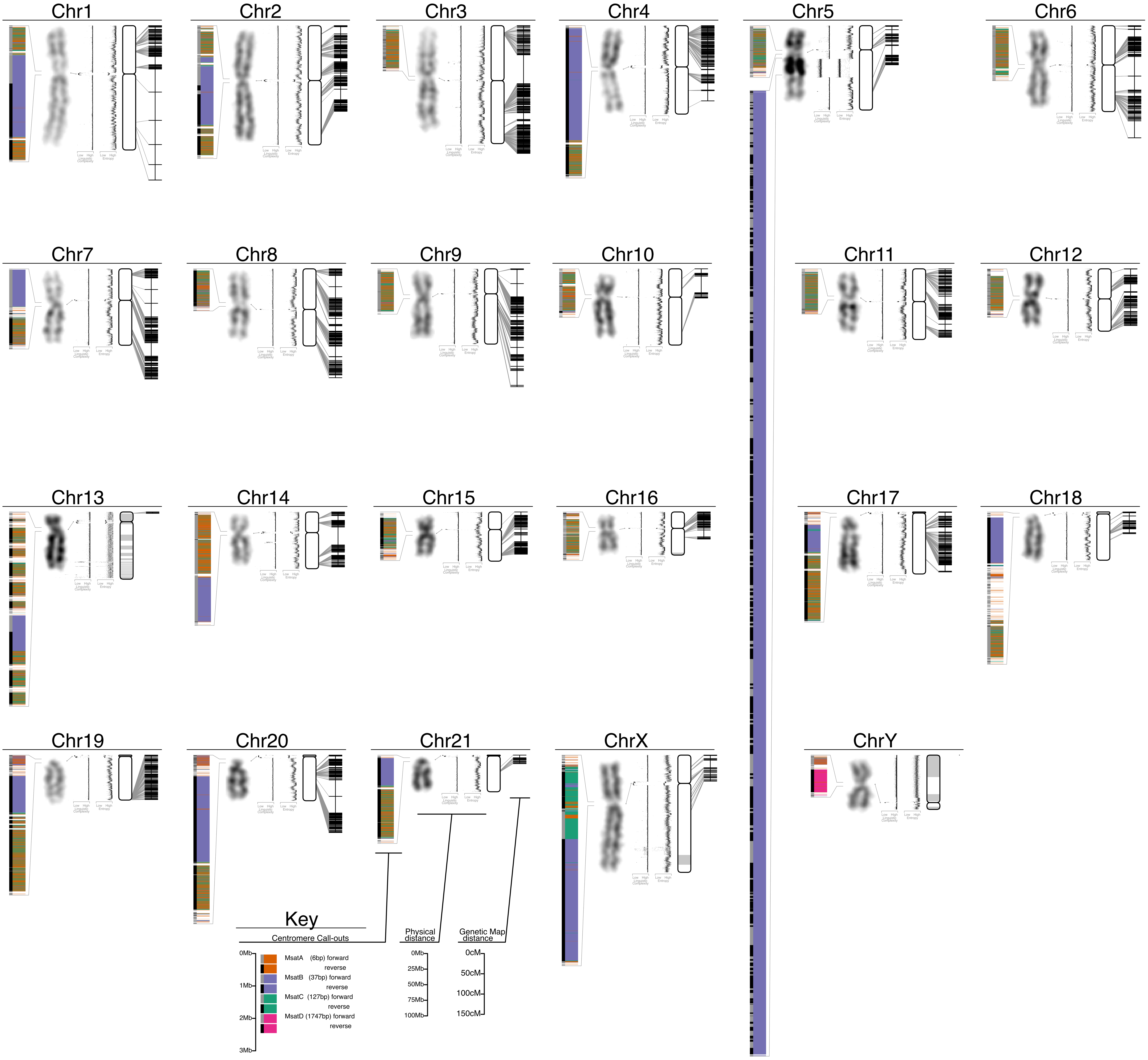
